# Predicting cellular responses to perturbation across diverse contexts with State

**DOI:** 10.1101/2025.06.26.661135

**Authors:** Abhinav K. Adduri, Dhruv Gautam, Beatrice Bevilacqua, Alishba Imran, Rohan Shah, Mohsen Naghipourfar, Noam Teyssier, Rajesh Ilango, Sanjay Nagaraj, Mingze Dong, Chiara Ricci-Tam, Christopher Carpenter, Vishvak Subramanyam, Aidan Winters, Sravya Tirukkovular, Jeremy Sullivan, Brian S. Plosky, Basak Eraslan, Nicholas D. Youngblut, Jure Leskovec, Luke A. Gilbert, Silvana Konermann, Patrick D. Hsu, Alexander Dobin, Dave P. Burke, Hani Goodarzi, Yusuf H. Roohani

## Abstract

Cellular responses to perturbations are a cornerstone for understanding biological mechanisms and selecting drug targets. While machine learning models offer tremendous potential for predicting perturbation effects, they currently struggle to generalize to unobserved cellular contexts. Here, we introduce State, a transformer model that predicts perturbation effects while accounting for cellular heterogeneity within and across experiments. State predicts perturbation effects across sets of cells and is trained using gene expression data from over 100 million perturbed cells. State improved discrimination of effects on large datasets by more than 30% and identified differentially expressed genes across genetic, signaling and chemical perturbations with significantly improved accuracy. Using its cell embedding trained on observational data from 167 million cells, State identified strong perturbations in novel cellular contexts where no perturbations were observed during training. We further introduce Cell-Eval, a comprehensive evaluation framework that highlights State’s ability to detect cell type-specific perturbation responses, such as cell survival. Overall, the performance and flexibility of State sets the stage for scaling the development of virtual cell models.

## 1. Introduction

Therapeutic discovery relies on accurately predicting the impact of cellular perturbations. Ranging from genetic interventions such as CRISPR or RNAi, to chemical treatments with small molecules or biologics, these perturbations serve not only to induce desired phenotypes, but are also central to establishing causal relationships between genes, pathways, and cellular outcomes, thus uncovering deeper insights into cellular function. By selectively disrupting specific components of cellular systems, scientists can identify causal drivers of phenotypes, an essential step in both target identification and drug development. Experimental perturbation technologies enable researchers to probe the effects of interventions along two main axes: the type of perturbation applied and the cellular or biological context. Both factors profoundly influence the system’s response. Advances in functional genomics now enable large-scale screening in specific cellular contexts, often through approaches like pairing pooled CRISPR perturbations with transcriptome-wide readouts at the single-cell level (Dixit et al., 2016; Datlinger et al., 2017; Przybyla and Gilbert, 2022; Replogle et al., 2022; Norman et al., 2019; Feng et al., 2024). However, these assays remain cost-prohibitive and labor-intensive to scale across many contexts. Improving our ability to generalize perturbation response predictions across diverse biological contexts would greatly accelerate causal target discovery, deepen our understanding of cellular function and disease, and in turn facilitate the design of context-specific interventions, creating a foundation for personalized treatment predictions.

A range of computational approaches have been developed to tackle this problem (Lotfollahi et al., 2019, 2023; Bunne et al., 2023; Roohani et al., 2024a; Cui et al., 2024; Hao et al., 2024; Ji et al., 2021). However, despite the rapid growth of perturbation datasets in size and scope, proportional gains in predictive capabilities have not been achieved (Wu et al., 2024; Chevalley et al., 2022; Li et al., 2024b,a; Wenteler et al., 2024). Current deep learning methods do not consistently outperform linear models when generalizing perturbation effects across cellular contexts (Wu et al., 2024; Li et al., 2024b). We argue that this is primarily caused by two major sources of noise that mask true perturbation effects in single-cell perturbation datasets: biological heterogeneity within the studied population that is not explained by experimental covariates, and technical or experimental variation across different perturbation datasets (**Fig. 1A** and **Eq. 1**).

**Figure 1:**
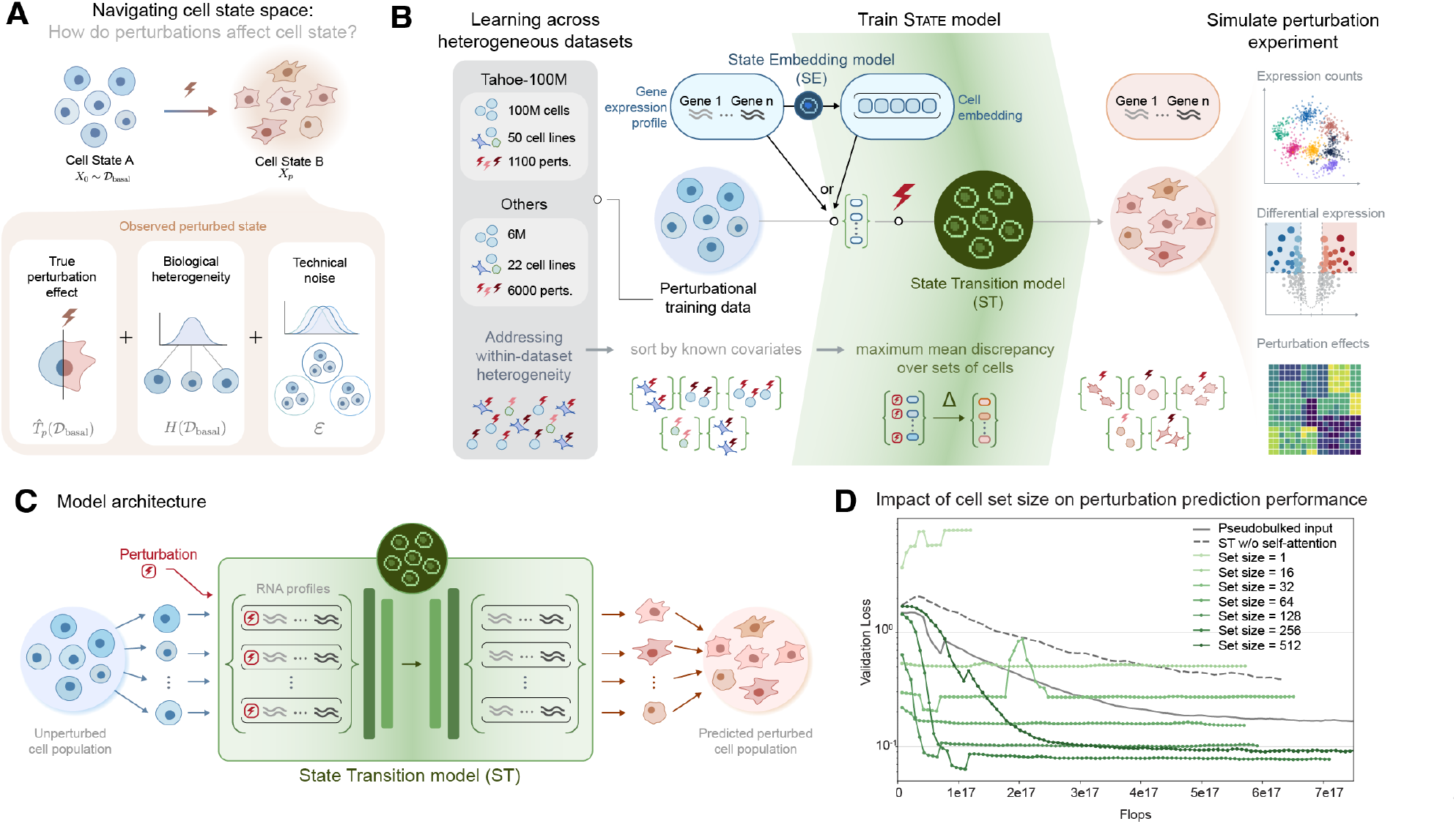
State: A transformer-based model for predicting perturbation effects across sets of cells. **(A)** Modeling perturbation effects at single-cell resolution requires disentangling biological signals from confounding variation introduced by noise, batch effects, and heterogeneity across similarly treated cells. **(B)** State is a multi-scale machine learning architecture that operates across genes, individual cells, and cell populations. The core State Transition model (ST) learns perturbation effects by training on sets of perturbed and unperturbed cell populations grouped by shared covariates (e.g., perturbation type, cell context, and batch). ST can operate directly on gene expression profiles or on compact cell representations from the State Embedding model (SE), which learns information-rich embeddings from large-scale observational data. This multi-scale architecture allows ST to effectively simulate perturbation experiments *in silico* and support downstream analyses such as expression quantification, differential gene expression analysis, and estimation of perturbation effect sizes. **(C)** ST is a transformer model that takes sets of unperturbed cell populations and perturbation labels as input to predict corresponding perturbed cell populations. When using gene expression profiles to represent cells, ST directly predicts transcriptomes at single-cell resolution. When using State embedding inputs, ST predicts output embeddings that are then decoded with an MLP to predict transcriptomes. **(D)** Increasing the size of cell sets improves validation loss up to an optimal point, with best performance on the Tahoe-100M dataset achieved when covariate-matched groups are chunked into sets of 256 cells. The full ST model significantly outperforms a pseudobulk model (State with mean-pooling instead of self-attention) and a single-cell variant (State with set size = 1). Removing the self-attention mechanism (State w/o self-attention) substantially degrades performance, highlighting the importance of modeling interactions between cells within a set.

The challenge of modeling biological heterogeneity is driven by an inherent limitation of single-cell RNA sequencing: the destruction of cells during measurement prevents observation of their pre-perturbation states and accurate inference of each cell’s specific perturbation response. To address this, perturbation effects are inferred by comparing populations of perturbed and unperturbed cells, while attempting to resolve heterogeneity at the level of cell type, batch, or other population-level covariates. Some approaches assume that within-population heterogeneity is negligible compared to perturbation effects and simply map perturbed cells to randomly selected unperturbed cells with shared covariates (Roohani et al., 2024a), a mapping approach that has also been tested with expressive transformer-based models (Cui et al., 2024; Hao et al., 2024). Although effective in datasets where perturbation effects are strong (Norman et al., 2019), these approaches often fail to generalize when perturbation effects are more subtle and heterogeneity in the unperturbed population may even exceed the perturbation signal. This is particularly evident in cases of variation in cell cycle state, lineage bias, or pre-existing epigenetic programs and even more so when the basal population is itself drawn from diverse cell types such as in *in vivo* studies (Lara-Astiaso et al., 2023; Saunders et al., 2024). Other models treat cell populations as distributions, employing generative approaches like variational autoencoders to learn data-generating distributions or explicitly disentangle labeled and unlabeled sources of variation (Lotfollahi et al., 2023; Piran et al., 2024; Bereket and Karaletsos, 2024; Weinberger et al., 2023; Lopez et al., 2023; Papalexi et al., 2021; Weinberger et al., 2024; Lopez et al., 2018; Song et al., 2025). However, in practice, these models often fail to meaningfully outperform methods that do not explicitly model distributional structure when applied to the prediction of perturbation effects (Wu et al., 2024). Optimal transport-based methods that map unperturbed to perturbed populations have also been proposed, but their applicability has been limited by strong assumptions and poor scalability (Bunne et al., 2023, 2024b; Jiang et al., 2024; Ryu et al., 2024).

The second major source of noise is technical, arising from limitations in the data itself rather than the model. In genetic perturbation experiments, the intended effects, such as gene knockout or knockdown, may not always occur in each targeted cell, leaving cells incorrectly labeled as perturbed (Peidli et al., 2024; Papalexi et al., 2021; Weinberger et al., 2023). Additional variability from experimental conditions, including transduction efficiency, RNA sequencing depth, reagent chemistry, and timing of collection, further complicate data integration across different studies (Bock et al., 2022). Together, these technical confounders dilute the true perturbation-derived signal in the data, thereby constraining the development of models that can generalize robustly across distinct datasets. While single-cell foundation models have emerged as a strategy for learning robust cell representations across datasets (Theodoris et al., 2023; Rosen et al., 2023; Cui et al., 2024; Hao et al., 2024; Ho et al., 2024; Chen and Zou, 2024; Pearce et al., 2025; Heimberg et al., 2016; Szałata et al., 2024), they are currently unable to meaningfully distinguish between subtler variations such as those driven by genetic perturbations as they have generally been optimized to differentiate between broader categories such as cell type (Luecken et al., 2022, 2025).

**Modeling heterogeneity in single cell perturbation experiments**

The observed log-normalized perturbed expression state of each cell (*X*_*p*_) can be modeled based on its unperturbed state. However, since the unperturbed state of the cell is unobservable, we approximate *X*_*p*_ as

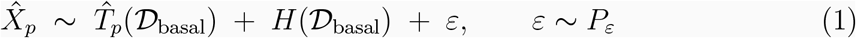

where

- 𝒟_basal_: The distribution of the unperturbed, baseline cell population.
- 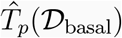: True effect caused by perturbation *p* on the population.
- *H*(𝒟_basal_): Biological heterogeneity of the baseline population.
- *ε*: Experiment-specific technical noise, assumed independent of the unperturbed cell state and 𝒟_basal_.

This 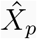 serves as a distributional analogue of *X*_*p*_, that is 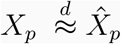, enabling modeling based on observable population characteristics.

To overcome these challenges and advance towards effective virtual models of cell state, we introduce State, a flexible and expressive architecture for modeling cellular heterogeneity and perturbation effects within and across diverse datasets. State is a multi-scale model with two complementary modules: a State Transition model (ST) and a State Embedding model (SE). ST is a transformer that uses self-attention to model perturbation-induced transformations across sets of cells, where each cell is represented either by its raw gene expression profile or a learned embedding. SE is pretrained to generate expressive cell embeddings by learning gene expression variation between cells across diverse datasets (Zhang et al., 2025; Program et al., 2025; Youngblut et al., 2025), yielding representations that are robust to technical variation and optimized for detecting perturbation effects. By leveraging self-attention over sets of cells, ST can flexibly capture biological heterogeneity without relying on explicit distributional assumptions. Together, SE and ST enable State to generalize across many datasets and contexts, improving transferability of perturbation-response modeling.

The multi-scale architecture of State enables it to leverage both 167 million cells of observational data to train its embedding model and over 100 million cells of perturbation data to train a transition model. We evaluate State on several large-scale datasets, including drug-based perturbations (Zhang et al., 2025; Srivatsan et al., 2020), cytokine signaling perturbations (Parse Biosciences, 2023), and genome-scale genetic perturbations (Replogle et al., 2022; Nadig et al., 2025; Jiang et al., 2025; McFaline-Figueroa et al., 2024; Feng et al., 2024). To fully assess the ability of State and other models to simulate cellular perturbations, we present Cell-Eval, a comprehensive evaluation framework that goes beyond conventional metrics based on expression counts to include a suite of biologically relevant and interpretable metrics focused on differential expression prediction and estimation of perturbation strength.

Across all metrics and data scales spanning multiple orders of magnitude, State consistently outperforms both naive and state-of-the-art models. To our knowledge, it is the first model to consistently outperform simple linear baselines in generalizing perturbation effects across cellular contexts. Moreover, we show that modeling perturbations in lower data regimes with the State embedding enables the detection of strong responses in novel cell types, when no perturbation data for those cell types are used during training. For example, we demonstrate that pretraining State on the Tahoe-100M dataset (Zhang et al., 2025) improves the generalization of perturbation effects to unseen cellular contexts. Thus, State presents a scalable approach for learning perturbation effects that transfer across datasets and experimental settings.

Beyond empirical performance, we provide novel theoretical results that connect State to Optimal Transport (OT) theory, a commonly used method for modeling cellular heterogeneity in response to perturbations (Bunne et al., 2023; Chen et al., 2024; Demir et al., 2024; Dong et al., 2023; Bunne et al., 2024b; Ryu et al., 2024; Jiang et al., 2024). Specifically, we prove that, under mild regularity conditions and in an asymptotic limit, the unique continuous OT map between unperturbed and perturbed cell populations lies within the solution family of State. This result positions State as a generalization of OT-based approaches: while it can recover the classical OT solution, it also allows for more flexible modeling of perturbation effects that may not adhere to the assumptions and constraints imposed by standard OT formulations.

## 2. Results

### 2.1. Building the State Transition model for predicting perturbation effects on sets of cells

State is a multi-scale machine learning architecture that predicts downstream transcriptomic responses to cellular perturbations, including gene expression changes, differentially expressed genes, and overall perturbation effect sizes (**Fig. 1B**). It leverages (i) at the molecular level, embeddings that represent individual genes across experiments and species; (ii) at the cellular level, embeddings that capture the transcriptomic state of each individual cell, represented either as the cell’s log-normalized transcriptome or as embeddings generated by the State Embedding model (SE); and (iii) at the population level, the State Transition model (ST) learns perturbation effects across sets of cells. State can leverage both observational and interventional data during training: SE is trained on 167 million human cells drawn from multiple large observational single-cell repositories (Youngblut et al., 2025; Zhang et al., 2025; Program et al., 2025), and ST is trained on over 100 million chemically or genetically perturbed cells from large-scale single-cell screens (Zhang et al., 2025; Parse Biosciences, 2023; Replogle et al., 2022).

The core motivation for ST is to model cellular heterogeneity beyond known covariates, such as cell type and perturbation label, to improve perturbation response prediction. To achieve this, cells are first stratified by known covariates (**Fig. S1**). For each covariate-matched perturbed group, ST constructs non-disjoint cell sets of fixed size, which serve as input during training and are paired with unperturbed control cell sets of equal size and matched covariates. Conditioned on the perturbation, ST uses a transformer backbone to perform repeated bidirectional self-attention and feed-forward operations across control cell sets (**Section 4.3, Fig. S2A**). This enables ST to model heterogeneity within the input cell set while predicting downstream transcriptomic responses to perturbation (**Fig. 1C**).

ST is trained using a maximum mean discrepancy (MMD) loss between predicted and observed transcriptomes of perturbed cells. While ST learns perturbation effects across distributions of cells, it still predicts perturbed cell profiles for individual cells, a feature that is important for learning distributional structure of a perturbed population. Empirical results show that increasing cell set size, up to a threshold, achieves much lower validation loss compared to losses on individual cells, whether they are true samples or pseudobulked across neighboring cells (**Fig. 1D**). Furthermore, removing the self-attention leads to degraded performance (**Fig. 1D**), highlighting the value of flexible set-based self-attention for modeling cellular heterogeneity relevant to perturbation response prediction.

### 2.2. State outperforms baselines in predicting perturbation effects across cell contexts

We tested the State architecture on a generalization task assessing its ability to predict perturbation effects in new cellular contexts, such as unseen cell lines or donors. Specifically, we implemented an underrepresented context generalization task (**Section 4.2.1**), in which each model had access to 30% of perturbations in the test context during training (**Fig. 2A**). We benchmarked performance against several baselines (**Section 4.7.5**), including a simple linear model (Ahlmann-Eltze et al., 2024), two autoencoder-based models CPA (Lotfollahi et al., 2023) and scVI (Lopez et al., 2018), and a single cell foundation model scGPT (Cui et al., 2024). We also included two naive mean-based baselines that explain a significant portion of observed variance in cell-type generalization tasks (**Fig. 2A**). The “context mean” baseline predicts the average expression observed in the training data for a given cell context across all perturbations (Kernfeld et al., 2023), while the “perturbation mean” baseline predicts the average perturbation effect across training cell contexts applied to the basal expression for a given cell context. In our results, we refer to baselines predicting mean expression or mean perturbation effect as “mean baselines” and the other models as “baseline models”. All models (including State) were trained to predict the log-expression of the top 2,000 highly varying genes (HVGs), a commonly used feature space for baseline comparisons.

**Figure 2:**
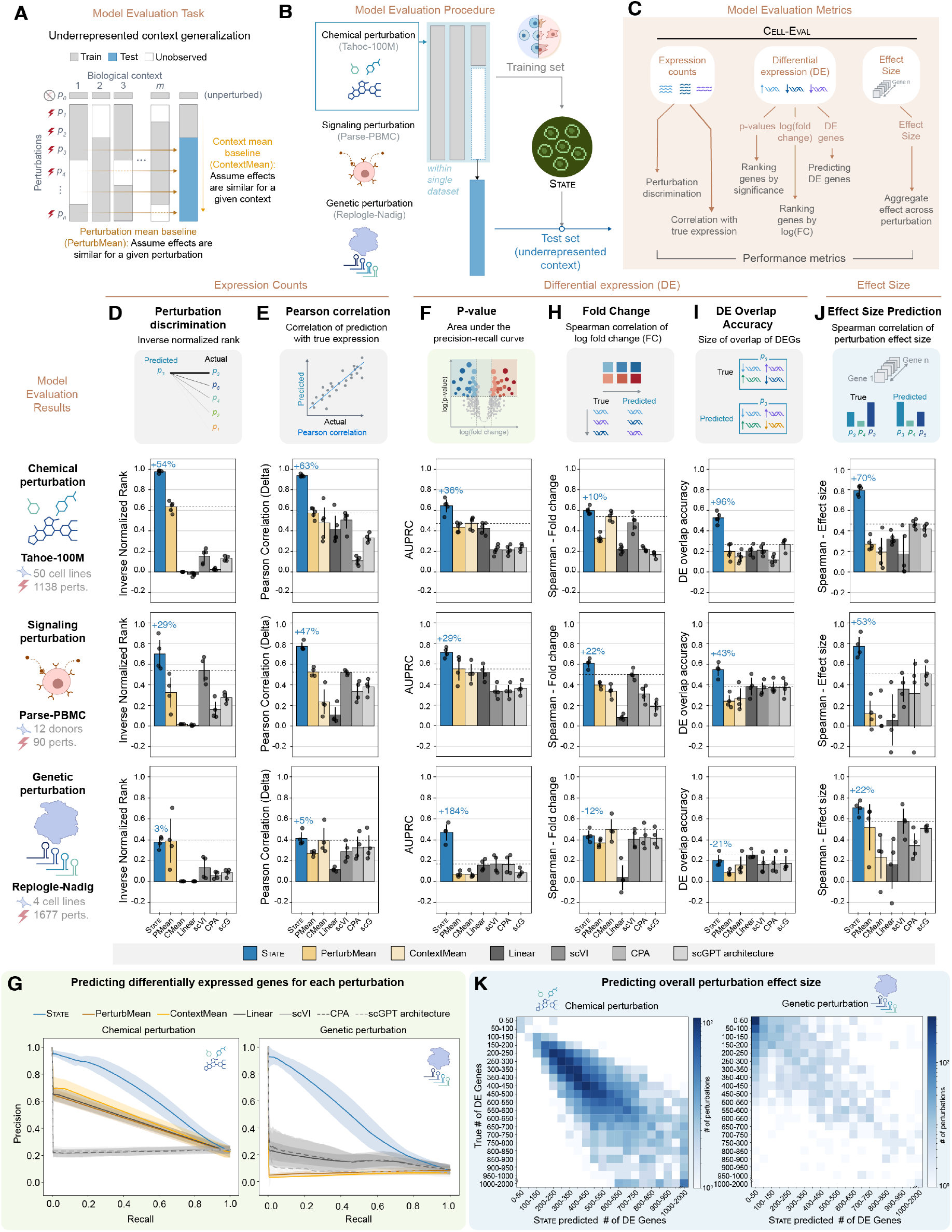
State outperforms existing baselines in predicting perturbation effects across cell contexts. **(A)** Underrepresented context generalization task. Models were trained on perturbation data from one or more cell contexts and evaluated on their ability to predict the effects of the same perturbations in a largely held-out and underrepresented target context. The “perturbation mean” baseline estimates effects by averaging observed differences between perturbed and control states across training cell contexts. The “context mean” baseline uses the average expression profile of the target cell context across all training perturbations. **(B)** Models were trained and evaluated on chemical, signaling, and genetic perturbation datasets (Zhang et al., 2025; Parse Biosciences, 2023; Replogle et al., 2022; Nadig et al., 2025). Training and test contexts were drawn from one dataset at a time. Comparisons included the mean baselines from (A), a simple linear model (Ahlmann-Eltze et al., 2024), autoencoder-based models (scVI (Lopez et al., 2018), CPA (Lotfollahi et al., 2023)), and a foundation model (scGPT (Cui et al., 2024)). **(C)** Performance was assessed using Cell-Eval metrics (**Section 4.7**) on standard Perturb-Seq outputs: expression counts and differentially expressed (DE) genes, with the following highlighted: **(D)** Perturbation discrimination score, measured using inverse normalized rank. **(E)** Pearson correlation between predicted and observed change in post-perturbation log-normalized expression counts. **(F)** Area under the Precision-Recall curve for model predictions of DE genes. **(G)** Precision-Recall curves for model predictions of DE genes. **(H)** Spearman correlation of log fold changes for significant DE genes between predictions and true values. **(I)** Overlap in top DE genes, defined as the percentage of significant genes in observed data that were also predicted as significant in predictions. **(J)** Spearman correlation between predicted and true overall perturbation effect size. **(K)** Confusion matrix comparing predicted and observed perturbation effect sizes, measured by the number of differentially expressed genes per perturbation.

We evaluated State on chemical perturbation data from the *Tahoe-100M* dataset (Zhang et al., 2025), cytokine signaling perturbations from Parse Biosciences (Parse Biosciences, 2023) (abbreviated Parse-PBMC), and genetic perturbation data from Replogle et al. (2022) and Nadig et al. (2025) (abbreviated Replogle-Nadig)(**Fig. 2B**). Tahoe-100M includes perturbation responses from 50 diverse cancer cell lines treated under 1,138 conditions involving 380 distinct drug perturbations. Parse-PBMC contains 90 cytokine perturbation responses across 12 donors and 18 cell types. Replogle-Nadig consists of 2,024 genetic perturbations applied to four distinct cell lines after filtering perturbations with low on-target efficacy. For all datasets, we trained State directly on cell representations derived from highly variable genes (ST+HVG). Training and test contexts were drawn only from one dataset at a time.

To test generalization, we implemented a careful data splitting strategy: for the Tahoe100M dataset, we plotted a PCA using pseudobulked expression values for the fifty available cell lines to visually identify distinct phenotypic clusters. From these, five cell lines were chosen to be in the test set for final model evaluation (**Fig. S3**). No data from these cell lines was observed throughout the model development process. In a separate evaluation, we iteratively held out all cells from 11 distinct organs for testing. For the Parse-PBMC dataset, we held out 4 random donors from the 12 donor cell lines. For each of these held-out contexts, 30% of its perturbations were randomly removed from the test data and included in the respective training data. For the Replogle-Nadig dataset, we conducted an evaluation by iteratively holding out one cell line as a test set. For each iteration, models were trained on the remaining three cell lines plus an additional 30% of perturbations randomly sampled from the test cell line.

Our evaluation framework captures key outputs of a single-cell perturbation experiment which are well represented through three readout categories: (1) gene expression counts, (2) differential expression (DE) statistics, including identification of differentially expressed genes (DEGs) and their log fold changes, and (3) the magnitude of the perturbation effect (e.g., the total number of DEGs) (**Fig. 2B**). To comprehensively assess model performance across these dimensions, we developed a suite of evaluation metrics, Cell-Eval (**Section 4.7, Fig. 2C**). These metrics are designed to be both expressive and biologically interpretable, offering complementary insights. For example, while overlap in DEGs helps link predictions to specific pathways giving them biological significance, it may be less sensitive to fine-grained changes compared to the perturbation discrimination score, which captures the similarity between predicted and true perturbation effects. Moreover, by benchmarking against naive baselines, these metrics provide a clearer assessment on generalization performance versus memorization of training-set effects.

A central goal of perturbation experiments is to identify perturbations that optimally drive desired transcriptomic states. For a model to do this, it must be able to effectively distinguish between different perturbation effects. Using a variant of the perturbation discrimination score adapted from Wu et al. (2024), which ranks predicted post-perturbation expression profiles by their similarity to the true perturbation outcomes, State achieved an absolute improvement of 54% and 29% on the Tahoe-100M and PBMC datasets respectively (**Fig. 2D**). On genetic perturbation datasets, State matched the performance of the perturbation mean baseline and significantly outperforms all other baseline models.

To directly assess the accuracy of predicted gene expression counts, we computed the Pearson correlation between observed and predicted perturbation-induced expression changes. On this metric, State outperformed baselines by 63% on Tahoe-100M, 47% on Parse-PBMC and 5% on Replogle-Nadig. For the genetic perturbation dataset, this task is more challenging due to the subtler effects of perturbations. Notably, on this dataset, the best-performing baseline with performance comparable to State was the context mean rather than the perturbation mean. This highlights that State’s predictions were not trivially similar to either mean baseline (**Fig. 2E**).

To evaluate State beyond global ranking and correlation metrics, we conducted a systematic differential-expression (DE) analysis. Using a Wilcoxon rank-sum test, we identified differentially expressed genes post-perturbation, calculating both their log fold changes and adjusted p-values (false discovery rate). We decomposed our DE analysis into assessments of each component (p-value and log fold change) independently as well as in combination (DE gene overlap). To evaluate p-values for model-predicted DE genes, we first computed true significantly DE genes using the experimentally observed perturbation data while setting an FDR threshold of 0.05. P-values derived from model predictions were then compared to true significance levels using a precision-recall curve. Measuring the area under the precision recall curve, we found that State consistently outperforms all baselines across datasets (**Fig. 2F**). Notably, State’s AUPRC is 184% higher than the next best approach for the genetic perturbation dataset (**Fig. 2G, Fig. S4**).

For evaluating log fold change, we limited our analysis to true significant DE genes to limit confounding from predicted significance levels. The Spearman correlation was computed between the predicted and true log fold changes for each of these genes. Some machine learning baselines such as scVI showed strong performance on this metric, yet State’s performance was still over 10% higher than baselines for both Tahoe-100M and Parse-PBMC datasets (**Fig. 2H**). To simulate a practical DE analysis workflow, we selected DE genes using an FDR threshold of 0.05 applied to model-predicted p-values, then ranked this set by log fold change and compared it to the equivalent set derived from true p-values and fold changes. Using different setting of overlap size (*k* = 50, 100, 200), we observe strong performance by State across all three datasets and all three settings of *k* (**Fig. S5A**). For completeness, we also evaluated the model on a variable sized overlap by setting *k* to be the same size as the number of true differentially expressed genes. State is twice as good as the next base baseline (scGPT) on the Tahoe-100M dataset and 43% better than the corresponding best baseline (Linear) on the Parse-PBMC dataset (**Fig. 2I**). To assess a more practically relevant scenario of minimizing false positives, we also measured the proportion of predicted top *k* DE genes that were significant at all in the observed experimental data (precision at *k*). Across datasets and settings of *k*, State showed much stronger performance than baselines (**Fig. S5B**).

Moving beyond identification of individual DE genes, we assessed the accuracy of models in predicting overall perturbation effects by counting the total number of DE genes predicted for each perturbation. State accurately ranked perturbations by their relative effect sizes, achieving Spearman correlations 53% higher on Parse-PBMC and 22% higher than baselines on Replogle-Nadig, and 70% higher on Tahoe-100M approaching an absolute correlation of 0.8 (**Fig. 2J**). Looking at the trend across individual perturbations, we observe that State can predict perturbation effects across both datasets with large effect sizes (i.e. drug perturbations in Tahoe-100M) as well as those with subtler magnitude of effects, such as the genetic perturbation dataset Replogle-Nadig (**Fig. 2K**). These results suggest that even when the specific genes predicted to be differentially expressed after genetic perturbation do not always match those observed experimentally (**Fig. 2I**), State can still accurately estimate the overall size of the DE gene set. Moreover, because the set of experimentally observed DE genes may be influenced by experiment-specific factors, and because overlap-based metrics can be overly stringent for perturbations with subtle effects, the size of the DE gene set offers a more robust and complementary indicator of model performance in genetic perturbation experiments, even though it is less directly interpretable.

Finally, to evaluate generalization under a more challenging data split, we assessed performance on a held-out tissue. In this setting, State was trained on data from all tissues except one, and evaluated on the held-out tissue. Across multiple metrics, State consistently outperformed the perturbation mean baseline (**Fig. S6**).

While State consistently outperformed all baseline models across datasets with few exceptions, the performance gains over mean baselines were notably larger on Tahoe-100M, which includes 100 million cells spanning thousands of perturbations across dozens of baseline cellular contexts, and Parse-PBMC, which includes 10 million cells across 12 donors and 18 cell types, as compared to the genome-scale genetic perturbation datasets conducted in just a few cell lines. This highlights State’s ability to leverage data scale and context diversity more effectively than existing models, which do not display proportionate gains in performance despite more data. Moreover, even in the case of the genetic perturbation dataset, where some baselines showed occasional benefit over State on certain metrics (Linear model outperformed on DE overlap by 20% and the context mean baseline outperformed on fold change prediction by 12%), these models were unable to consistently outperform across multiple metrics. In contrast, State demonstrated the most consistent performance overall.

### 2.3. State embeddings enhance zero-shot perturbation prediction across contexts

One of the goals in developing virtual models of cell state is to create general-purpose predictive models that can be applied to new contexts, even in the absence of perturbation data for those contexts (Bunne et al., 2024a). These models should also be able to learn cell regulatory information from one dataset and transfer it effectively to other datasets regardless of perturbation modality, such as chemical or genetic interventions. However, gene expression counts are subject to context-specific variability (e.g., sequencing depth and experimental platform), and do not always generalize well across studies.

To address this, we developed a unified cell representation that can be shared across datasets and experiments, enhancing perturbation prediction capabilities in previously unperturbed cellular contexts. The State Embedding model (SE) complements ST by learning cell embeddings that are optimized to capture cell-type specific gene expression patterns (**Fig. 3A**). When used with ST, the embedding enables a smoother landscape over cell states, learned using a vast repository of observational single-cell data (Program et al., 2025; Zhang et al., 2025; Youngblut et al., 2025). SE enables us to indirectly leverage observational single-cell data to improve perturbation response predictions, especially in cases where interventional data for a particular context is scarce or noisy.

**Figure 3:**
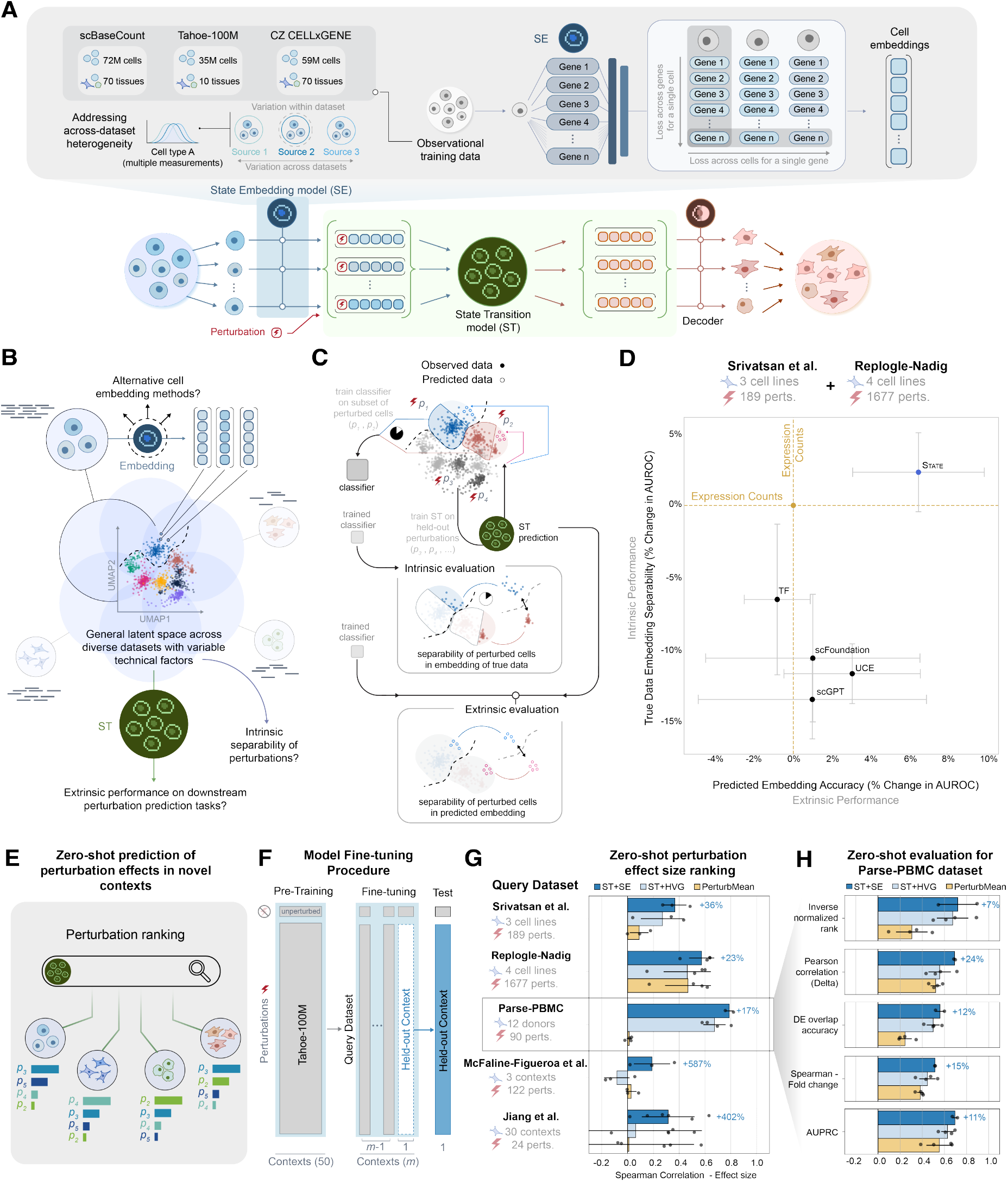
State embeddings enhance zero-shot perturbation effect prediction across datasets, experiments, and modalities. **(A)** The State Embedding model (SE) learns rich, generalizable representations of transcriptomic information across diverse datasets. Given a control (unperturbed) cell population, SE computes cell embeddings. ST then predicts how those embeddings shift in response to a specified perturbation, effectively modeling the distributional effect of the perturbation in latent space. Finally, a learnt decoder maps the predicted embeddings back into gene expression space. **(B)** Understanding the impact of a shared latent space across datasets for modeling perturbation effects across diverse cell contexts and perturbations. **(C)** Embedding quality is evaluated using both intrinsic and extrinsic metrics: intrinsic performance reflects classification accuracy of perturbed cells in the embedding generated using observed data; extrinsic performance measures the classification accuracy over perturbed embeddings predicted by ST trained on the cell embeddings. **(D)** Over two held-out perturbation datasets, using the intrinsic and extrinsic metrics, we evaluate State embeddings against cell embeddings generated by scFoundation (Hao et al., 2024), scGPT (Cui et al., 2024), Universal Cell Embedding (UCE) (Rosen et al., 2023), and Transcriptformer (TF) (Pearce et al., 2025). State embeddings consistently outperform comparable model embeddings, even passing the performance achieved using the original expression counts. **(E)** State embeddings enhance prediction of perturbation effects zero-shot, e.g. in novel cell contexts not seen perturbed at the time of training. **(F)** ST with State embeddings is pretrained using Tahoe-100M and fine-tuned using a query dataset consisting of one or more contexts, of which one is held out for zero-shot testing. ST is evaluated without any training using perturbations in the held-out cell context (**Section 4.2.2**). **(G)** Zero-shot performance in ranking perturbations by overall effect size in previously unseen cell contexts over 5 query datasets. **(H)** Zooming into Parse-PBMC, State embeddings also improve performance over other metrics from Cell-Eval. More datasets are shown in **Figure S7**.

Architecturally, the SE encoder is a dense, bidirectional transformer trained to predict log-normalized gene expression (**Section 4.4**). The SE decoder is a smaller, specialized MLP that predicts gene expression from a combination of the learned cell embedding and the target gene embedding (**Fig. S2B**). This architectural asymmetry encourages the learning of generalizable representations of cell state in a single vector embedding. SE is trained with a loss computed along two axes: it predicts expression across genes within each cell, and for each gene across cells in each minibatch. This dual-axis formulation encourages the model to capture relative variation in gene expression both within individual cells and across the population (Lal et al., 2024; Ding et al., 2025; Fischer et al., 2024). The loss enhances the model’s sensitivity to perturbation effects by preserving the inter-cellular variability necessary for accurate differential expression modeling.

By learning a general-purpose embedding that captures subtle cell-to-cell variation, SE addresses a core challenge in perturbation modeling: defining a transferable feature space across single-cell datasets (**Fig. 3B**). When SE and ST are used together, ST learns to predict perturbed cell embeddings, while simultaneously learning to decode those predicted embeddings to log expression space (**Fig. S2B**). To assess the quality of the embeddings produced by SE, we evaluated their ability to distinguish between perturbations. We measured intrinsic quality by testing how well the embeddings of observed cells cluster by perturbation label, and extrinsic quality by examining how well the predicted embeddings from ST preserve this separation (**Fig. 3C**). We compared SE against using gene expression counts directly, as well as cell embeddings generated by scFoundation (Hao et al., 2024), scGPT (Cui et al., 2024), Universal Cell Embedding (UCE) (Rosen et al., 2023), and Transcript-former (TF) (Pearce et al., 2025) across two held-out perturbation datasets not seen during SE training (Srivatsan et al., 2020; Replogle et al., 2022). To measure separability, we train a simple linear probe on the embeddings to predict the perturbation label called for that cell. State embeddings more effectively separated between perturbation phenotypes compared to all other foundation models and the original expression counts, surpassing even the performance achieved using the original data representation (**Fig. 3D**). This suggests that SE is, in some cases, capable of denoising Perturb-seq data. In the extrinsic evaluation, State embeddings also led to over a 6% absolute improvement in downstream perturbation classification accuracy compared to all baselines.

Projecting gene expression into a shared latent space also enables zero-shot identification of strong perturbation effects in new cell contexts (i.e., without explicit training in the new cell context) (**Fig. 3E**). We assessed the robustness of embeddings from SE by pretraining State (ST+SE) on Tahoe-100M and fine-tuning the model on smaller datasets with new cell contexts, which we denote as *query datasets* (**Fig. 3F**). To evaluate the model, we held out one cell context at a time from the query datasets, thus focusing on zero-shot context-level generalization within a dataset rather than zero-shot dataset transfer. In predicting perturbation effects for previously unperturbed contexts, State trained with the State embedding (ST+SE) consistently ranked perturbation by their effect sizes more accurately than both the perturbation mean baseline and State models trained directly on gene expression (ST+HVG). This evaluation was performed using five datasets - which included two genetic perturbation datasets (Jiang et al., 2025; McFaline-Figueroa et al., 2024) and a drug perturbation dataset (Srivatsan et al., 2020) in addition to datasets used for the previous analyses, producing a total of 2,102 perturbations (**Fig. 3G**) across 5 datasets and 40 cell contexts. For larger datasets like Parse-PBMC and Replogle-Nadig, using the State embedding achieved more than 17% improvement with an absolute Spearman correlation greater than 0.5. In smaller genetic perturbation datasets (e.g., McFaline et al. and Jiang et al.), where baseline performance was near zero, embeddings yielded several-fold improvements.

Dataset-specific processing and quality differences also affected other metrics. Notably, in datasets with strong perturbation effects, zero-shot improvements of using SE were consistent. For example, on the Parse-PBMC dataset, we saw an average of 15% improvement across all five metrics described in **Section 2.2** (**Fig. 3H**). When excluding cases where baseline HVG performance was below 10% (indicating noisy data), the fine-tuning improvements remained largely consistent across all 5 datasets tested (**Fig. S7**). These results show SE’s capacity for transferring learning across datasets where technical variation can confound the biological signal driven by perturbation. ST+SE model performance also consistently benefited from pre-training even for datasets without drug-based perturbations (e.g. genetic or signaling perturbations), highlighting the successful transfer of cell regulatory information across perturbation modalities (**Fig. S8**).

### 2.4. State can detect cell type-specific response to perturbations

To illustrate a practical application of State, we evaluated its ability to detect cell type-specific differential expression (**Fig. 4A**). This analysis focused on five held-out cell lines from the Tahoe-100M dataset (**Fig. 4A**). We identified perturbations with strong cell type specificity by comparing the overlap of DE genes and the Spearman correlation of log fold changes between State’s predictions and two baselines: the context mean and the perturbation mean. Improved performance relative to the perturbation mean baseline suggests that State learns perturbation effects that are specific to a given cell type. Similarly, gains over the context mean baseline indicate that the model can distinguish between different perturbations within the same cell line and is not trivially predicting the average expression for each cell line. Across perturbations, State consistently displayed superior ability to recover the true ranking of log fold changes for differentially expressed genes, outperforming both the context mean and perturbation mean baselines (**Fig. 4B, C**).

**Figure 4:**
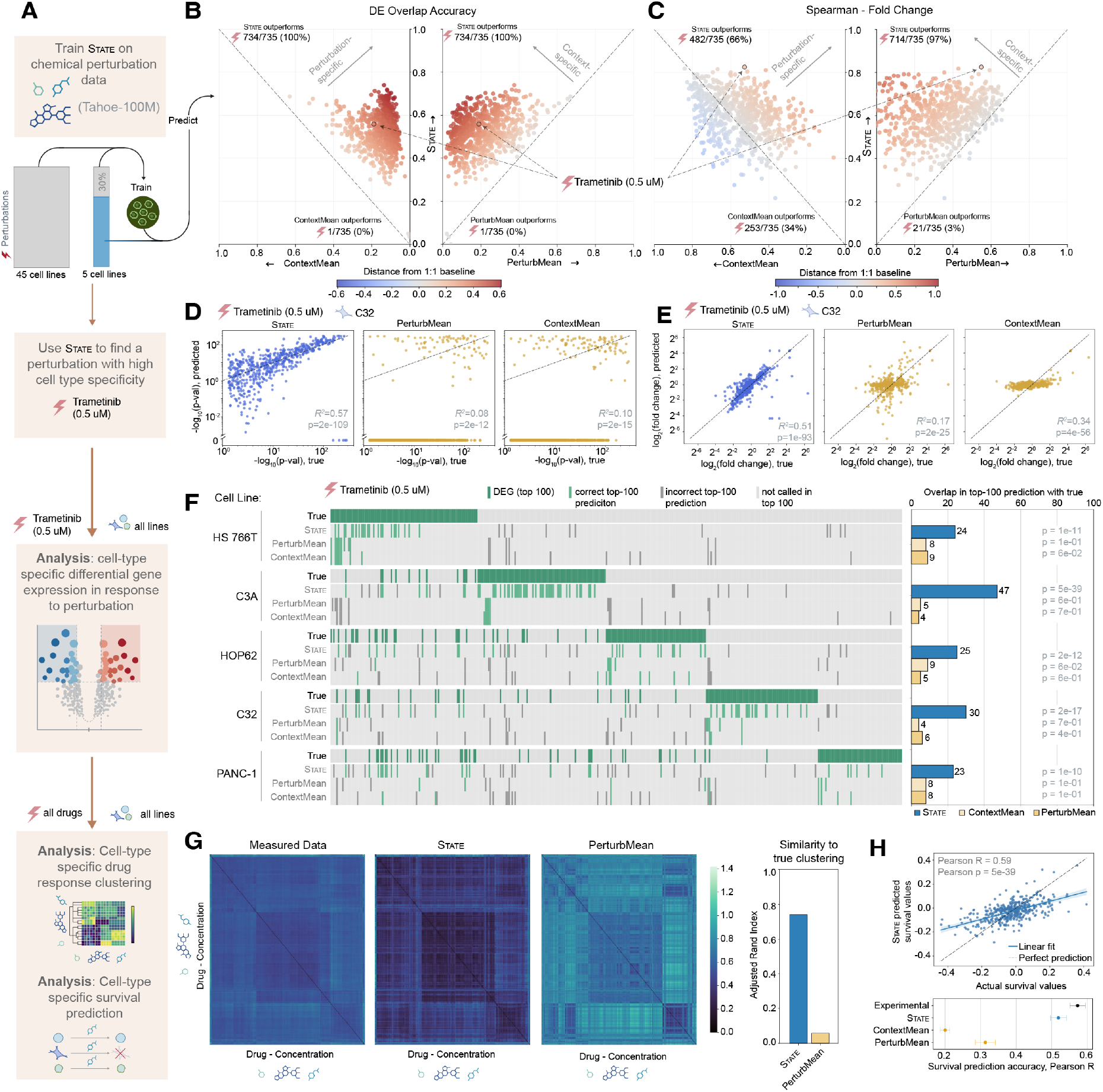
State detects cell type-specific gene expression modulations in response to perturbation. **(A)** Application of State for identifying perturbations with cell type-specific effects. **(B)** DE Gene Overlap between predicted and observed perturbation effects. **(C)** Spearman correlation between predicted versus observed log fold changes for differentially expressed (DE). For **(B)** and **(C)** Left: comparison between State and the context mean baseline. Right: comparison with the perturbation mean baseline. A specific held-out perturbation (Trametinib, 0.05 µM) shows substantially higher correlation for State relative to both baselines, indicating detection of perturbation effects that are both cell type specific but also not trivially predicted by the cell type mean. **(D)** Predicted versus observed significance values for DE genes following Trametinib (0.05 µM) perturbation, across the top 2,000 highly variable genes. State (blue) shows closer alignment with ground truth than baselines (yellow). Log fold changes for observed significant DE genes following Trametinib (0.05 µM) perturbation, comparing predictions from State (blue) and baselines (yellow). **(F)** Detection of cell type-specific DE genes across 5 cell lines following Trametinib (0.05 µM) perturbation. The top row for each cell line corresponds to the top 100 genes with the highest log fold change. Lower rows show the same set as predicted by State and baseline models. For each model, we also report the overlap in predicted DEGs and true DEGs computed with the observed data **(G)** Perturbation similarity heatmap for drug-induced differential expression across all held out cell lines. Each perturbation (row or column) corresponds to a specific drug-concentration pair. Heatmap computed using experimentally measured data, State predictions and perturbation mean baseline. Similarity of heatmaps to clustering in the measured data compared using Adjusted Rand Index. **(H)** State outperforms baselines at predicting cell survival.

To explore the biological relevance of a State-generated prediction, we ranked perturbations by how far they improved performance over the mean baselines, suggesting enhanced sensitivity to context-specific effects. Out of the top two perturbations from over 700 possible choices, one was an FDA-approved drug for BRAF-mutant melanoma and certain other tumors, Trametinib (Lugowska et al., 2015). We chose this perturbation (specifically, 0.5 *µM* Trametinib) since one of the five test cell lines, C32, is a melanoma line known to have the overactive BRAF mutation V600E (Banach et al., 2021). The model was not trained on C32 cells treated with any dosage of Trametinib. Both the predicted significance values for DE genes in C32 following perturbation with Trametinib (**Fig. 4D**) and the log fold changes for the true DE genes (**Fig. 4E**) showed much stronger alignment with ground truth for State relative to mean baselines. Notably, context and perturbation mean baselines exhibited little to no correlation with ground truth DE gene significance due to assigning extremely high levels of significance to the vast majority of genes. This inability to distinguish signal from noise underscores the limitations of simple averaging approaches, and highlights State’s advantage in capturing cell type-specific responses through its modeling of fine-grained cell-to-cell heterogeneity.

To further explore the cell-line specificity of perturbation response prediction, we compared State predictions to perturbation and context mean baselines for all five cell held-out cell lines. For each cell line, we identified the top 100 significantly differentially expressed (DE) genes based on absolute log fold change in the ground truth, and compared them to the top 100 predicted by State, using its own predicted significance values and log fold changes as thresholds (**Fig. 4F**). In every case, State correctly recovered over 30% of the top DE genes, while both baseline methods identified fewer than 7%. On average, State correctly identified *~* 20 additional cell line-specific DE genes relative to the baselines. This performance gap is particularly striking given that the top 100 genes were largely distinct across cell lines, with only an average of 16 DE genes shared within the top 100 across any pair of cell lines. Notably, in cases where genes were shared, State was also able to recover these shared effects, further supporting its ability to capture both unique and conserved perturbation responses across diverse cellular contexts.

To evaluate whether State broadly captures patterns of perturbation similarity, we compared the drug-induced perturbation effects across all drugs in this dataset in different contexts. For each held-out drug-concentration pair, we used the log_2_-fold changes in gene expression across cell lines as a fingerprint of that perturbation’s effect. We then computed pairwise similarities between these fingerprints to construct a perturbation similarity map (**Fig. 4G**). Comparing these predicted similarities to ground truth revealed a clear improvement for State over baseline models, achieving an Adjusted Rand Index (ARI) of over 0.7, compared to less than 0.1 for the perturbation mean baseline (**Fig. 4G**).

Moving beyond transcriptomic responses, we next assessed whether State could capture actual phenotypic outcomes. We trained a regression model to predict cell survival based on State-predicted gene expression profiles. The model was trained on real gene expression data across 10 folds while holding out 15% of the perturbations in each fold. Predicted survival values were strongly correlated with experimentally measured survival, achieving an average Pearson correlation of 0.52. In contrast, predictions based on the context mean baseline showed a low correlation of 0.2, while the perturbation mean baseline achieved a modest correlation of 0.31 (**Fig. 4H**).

While this is just one example of a potential use case for State, it shows the potential for future iterations of the model (or similar models) to provide guidance on the outcome of a particular treatment on a cell type that has not been previously tested with that treatment. This could have important implications for repurposing existing drugs, predicting patient-specific treatment response or understanding potential adverse reactions in cell types beyond the intended drug target.

## 3. Discussion

High-throughput cellular perturbation experiments, combined with recent advances in artificial intelligence, have opened exciting new avenues for developing computational models of cell state and behavior. Just as foundation models at the molecular scale have succeeded in predicting biological traits such as molecular structure and function (Brixi et al., 2025; Nguyen et al., 2024; Hayes et al., 2025; Lin et al., 2023), models at the cellular scale hold promise for uncovering mechanisms underlying disease progression and advancing the discovery of more precise treatments. However, translating these successes to the cellular level has proven substantially more challenging. Single-cell foundation models have yet to achieve the same level of predictive accuracy or generalizability, largely due to the inherent complexity of cells as dynamic, context-dependent systems. Unlike the relatively static and homogeneous nature of individual DNA, RNA, or protein molecules, cellular systems exhibit substantial heterogeneity and are deeply influenced by environmental and temporal factors. Two central challenges for modeling cellular systems are the intrinsic heterogeneity of cell populations undergoing perturbation which cannot be fully resolved using known covariates, and the variability introduced across different experiments and datasets.

Our model, State, presents a scalable approach for learning a foundation model of cell state and behavior across diverse cellular contexts and experimental conditions. The State Transition model (ST) uniquely learns perturbation effects across cell populations while still maintaining single-cell resolution in its predictions, thereby capturing residual heterogeneity not explained by known experimental or biological covariates. While past work has tackled heterogeneity in single-cell data by modeling cells as groups (Persad et al., 2023; Boyeau et al., 2022; Dann et al., 2022), State uniquely leverages a permutation-invariant transformer architecture (Lee et al., 2019; Zaheer et al., 2017) to directly model perturbation effects across unpaired distributions of cells. This frees State from predefined assumptions on distributional structure, unlike approaches based on optimal transport (Bunne et al., 2023; Chen et al., 2024; Demir et al., 2024; Dong et al., 2023). Moreover, our theoretical findings establish that, in the asymptotic limit and under mild regularity conditions, the unique continuous OT map (defined by a fixed quadratic cost) linking unperturbed and perturbed cell populations is provably contained within our model’s solution space (Theorem 2). This means State may learn the OT map when necessary, without explicitly constraining itself to a rigid, predefined OT formulation or requiring specialized architectural components typically associated with explicit OT solvers. Furthermore, by introducing explicit Jacobian-based constraints, we show that State can uniquely identify and learn this continuous optimal transport map (Theorem 3), positioning it as a powerful, theoretically grounded Neural OT solver.

The central goal for State is to reliably simulate single-cell perturbational experiments across diverse cell contexts. We used this objective to guide the design of a comprehensive model evaluation suite, Cell-Eval, focusing on performance along three key dimensions: gene expression counts, differential expression, perturbation effect sizes. To our knowledge, State is the first machine learning model to consistently outperform simple baselines such as the mean or linear models on the cell context generalization task, across almost all metrics and on multiple datasets. Moreover, the inclusion of embeddings from our cell embedding model, SE, enables more effective zero-shot perturbation effect prediction in novel cell contexts, a core goal for developing truly generalizable virtual cells. While State shows strong zero-shot performance when predicting perturbation effect sizes, metrics that reflect subtler changes, such as the accuracy of individual DE genes, are more sensitive to dataset size and quality. With the growth of large-scale perturbational atlases (Rood et al., 2024; Huang et al., 2025a; Zhang et al., 2025; Rood et al., 2025), we expect performance on these subtler metrics to improve, and State is well-positioned to leverage these datasets given its improved robustness to cellular and data variability. Furthermore, we have found that ST attention maps demonstrate sensitivity to cell set heterogeneity, highlighting interpretability as a promising avenue for further understanding gene regulatory mechanisms using State (**Section 6**).

When applied to previously seen contexts, we have found State’s predictions are biologically meaningful, accurately capturing context-specific perturbation effects, such as cell-type specific DE genes. Predicted post-perturbation profiles were also found to capture non-transcriptional cellular phenotypes such as cell survival. These capabilities position State as a step towards realizing the broader vision of a virtual cell–a model capable of exploring the space of possible cell states (Bunne et al., 2024a; Noutahi et al., 2025) and guiding autonomous systems, such as AI agents, for experimental design (Roohani et al., 2024b; Huang et al., 2025b; Rizvi et al., 2025; Wu et al., 2025). State contributes to this vision by advancing the state-of-the-art in perturbation-response modeling and by introducing an architecture that can meaningfully leverage the growing size and diversity of single-cell perturbation datasets. Its flexible transformer architecture can support a range of tasks that involve modeling transitions between distinct populations such as differentiation, development, and reprogramming. State is also agnostic to the choice of cell representation, which enables seamless extension to additional data modalities, including proteomics and morphological measurements, as well as literature-based knowledge. By incorporating mechanisms for perturbation featurization withing State, we expect to further lower the barriers between different modalities such as drugs and gene knockdowns, while also opening the door to combinatorial perturbation prediction. Although State has been trained and evaluated on over 70 cell contexts, we have not tested its usability on entirely held-out datasets for which no cell contexts have been seen during training. Further improvements in SE through training on larger datasets, such as scBaseCount (Youngblut et al., 2025), will be instrumental in generalizing predictions across such diverse settings.

## 4. Methods

### 4.1. Data Generation Process

We assume that the observed log-normalized perturbed expression state of each cell, *X*_p_, is a random variable generated from an *unobservable* unperturbed state, *X*_0_, which itself is a random variable representing the cell’s underlying expression state. *X*_0_ is drawn from a basal cell distribution 𝒟_basal_ for a given set of covariates (e.g., cell line, batch condition, etc), and the perturbation effect can be modeled as follows:

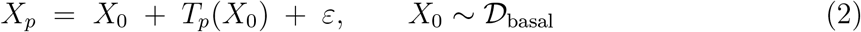

where

- *T*_*p*_(*X*_0_): True effect caused by perturbation *p*.
- *ε*: Experiment-specific technical noise, assumed independent of *X*_0_.

As single-cell transcriptomic measurements destroy the cell, *X*_0_ is unobservable, so directly modeling **Eq. 2** is not feasible. Instead, our method operates on the observable 𝒟_basal_ to predict the perturbed state, denoted as 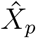. This forms the basis of our model:

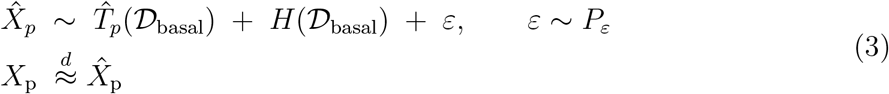

In this approximation, the true effect of the perturbation *T*_*p*_(*X*_0_) is now considered in the context of the entire basal population, denoted as 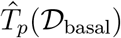. Additionally, we explicitly introduce *H*(𝒟_basal_) to represent the biological heterogeneity inherent in the baseline population. This heterogeneity was implicitly removed when sampling *X*_0_ *~* 𝒟_basal_ in the first equation but is made explicit here to reflect our shift to a distributional view. 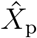 can be seen as a distributional analogue of *X*_p_, allowing us to model the perturbed state based on observable population characteristics rather than unobservable individual cell states.

**Fig. 1A** illustrates these assumptions. State is designed to reflect the assumptions of this data generation process: it directly models 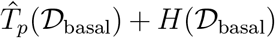 with State Transition (ST) and accounts for *ε* with State Embedding (SE).

### 4.2. Task Description

We now describe the core prediction tasks used for evaluation.

#### 4.2.1. Underrepresented Context Generalization Task

To evaluate the generalization capabilities of perturbation models, we first test the generalization of perturbation effects within Perturb-Seq datasets (**Fig. 2**). Given the dataset of interest, let the collection of biological contexts (cell types or cell lines in some dataset) be 𝒞 and the perturbation catalogue be 𝒫. For every context *c ∈* 𝒞, we write 𝒫_*c*_ *⊆* 𝒫 for the subset of perturbations that were profiled in that context, and we denote by *X*_*p,c*_ *∈* ℝ^*d*^ the gene–expression vector generated for the pair (*p, c*) according to **Eq. 2**. To test the generalization capabilities of the model in an underrepresented context, we fix a test context *c*^*^ *∈* 𝒞. We then choose a proportion *α* = 0.30 and draw a support set

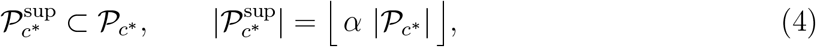

uniformly at random. The remaining perturbations form the target perturbation set 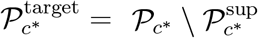. The training and test splits are then given as follows:

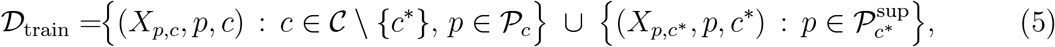

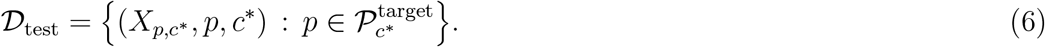

Thus, the model sees all perturbations in 𝒞\{*c*^*^} and only 30% of the perturbations in the test context *c*^*^. If |𝒞 | is large, a diverse fixed subset 𝒞_test_ *⊂* 𝒞 of *n* contexts is designated as test contexts. We form 𝒟_train_ from the 𝒞\𝒞_test_ contexts, as well as 30% of the perturbations from each *c*_*j*_ *∈* 𝒞_test_. We then construct multiple 𝒟_test_, one from each *c*_*j*_ *∈* 𝒞_test_, each containing the remaining unseen perturbations from *c*_*j*_. If |𝒞 | is small, we *iteratively* leave out a single context *c*^*^ = 𝒞\{*c*_*j*_}, form 𝒟_train_ with the remaining *c*_*j*_ contexts as well as 30% of the perturbations from *c*_*j*_ and 𝒟_test_ from the remaining unseen perturbations from *c*_*j*_. This iteration creates multiple training sets, allowing to report performance over separately trained models. In both cases, we report the mean loss across 𝒟_test_, yielding a robust estimate of the model’s ability to extrapolate to unseen perturbations within an underrepresented context. Unperturbed cells from all contexts including the underrepresented context are available to the model during training.

#### 4.2.2. Zero-Shot Context Generalization Task

Next, we evaluate the ability to generalize perturbation effects in a zero-shot manner across biological contexts (**Fig. 3F**). To enhance performance on this task, we include a pre-training step. Each model is initially pre-trained on one or more large perturbation datasets (e.g., Tahoe-100M) and subsequently fine-tuned to a different dataset, referred to as the query dataset, via full fine-tuning. The model is then tested on a held-out context within the query dataset.

Specifically, the model is first pre-trained on one or more datasets that include perturbations across multiple contexts. Then, given a query dataset, we hold out one context and use the remaining contexts for fine-tuning. The contexts used for fine-tuning contain *a superset of the perturbations* found in the held-out context. The fine-tuned model is evaluated on its ability to predict the effects of those perturbations within the held-out context of the query dataset. Formally, the fine-tuning process involves partitioning the set of contexts in the query dataset into 𝒞_fine-tune_ *⊂* 𝒞 and 𝒞_test_ *⊂* 𝒞 with 𝒞 _fine-tune_ *∩* 𝒞_test_ = ∅. The data used in the fine-tuning phase

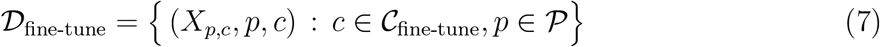

contain all perturbations in 𝒞_fine-tune_. After fine-tuning *f*_*θ*_ on 𝒟_fine-tune_, we evaluate it on the same perturbations *p ∈* 𝒫 but in unseen, held-out cell lines *c ∈* 𝒞_test_. Thus, the test set is

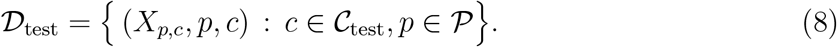

This task directly assesses whether the model has learned generalizable relationships between biological context and perturbation response. Unperturbed cells from all contexts including the held out context are available to the model during training.

### 4.3. State Transition Model (ST)

ST is a deep learning model that learns the transcriptomic responses to perturbations across populations of cells. The model uses a transformer architecture with self-attention across cells to predict perturbation effects between control and perturbed cell distributions. Let

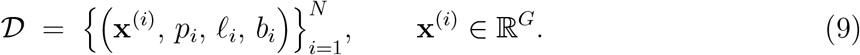

denote a dataset of single–cell RNA-sequencing measurements over the gene set, with *G* the number of genes. Here, **x**^(*i*)^ represents the normalized expression vector for cell *i*, after its raw count values have been processed. Specifically, the raw count value observed for gene *j* in cell *i* after perturbation *p*_*i*_ (if any) is initially 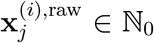. These raw counts are depth-normalized and log-transformed using Scanpy(normalize_total *→* log1p), yielding non-negative, real-valued expression vectors **x**^(*i*)^. Each cell is annotated with perturbation label *p*_*i*_ *∈* {1, …, *P*} *∪* {`ctrl`}, a biological context or cell line label *ℓ*_*i*_ *∈* {1, …, *L*}, and an optional batch effect label *b*_*i*_ *∈* {1, …, *B*}.

#### 4.3.1. Formation of Cell Sets

We group cells into sets based on their biological context, perturbation, and batch labels:

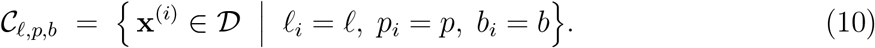

Let *N*_*ℓ,p,b*_ = |𝒞_*ℓ,p,b*_| denote the number of cells in each group. Given a target set size *S*, we partition 𝒞_*ℓ,p,b*_ into non-overlapping subsets of size *S* to construct a collection of tensors 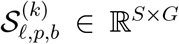, where 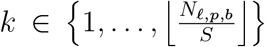. If *N*_*ℓ,p,b*_ is not divisible by *S*, the remaining cells form a final, smaller set, which is padded to size *S* by sampling additional cells with replacement from itself.

#### 4.3.2. Training on Cell Sets

During training, each set 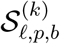, where *p ∈* {1, …, *P*} {`ctrl`}, is paired with a corresponding control set. The control set is constructed by randomly sampling *S* control cells from the same cell line *ℓ*, and optionally same batch *b*. Formally, for each training example 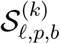, we construct a paired control set via a `map` operation:

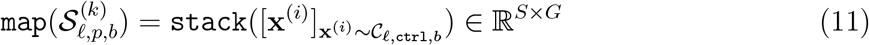

This mapping is applied uniformly, regardless of whether 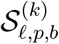 contains perturbed or control cells. The choice of `map` function effectively determines which sources of variation are explicitly controlled for. By conditioning on specific covariates (e.g., cell line *ℓ*, batch *b*), the mapping function reduces known sources of heterogeneity that could otherwise confound true perturbation signals. Different mapping strategies introduce different priors on non-perturbational variation. For instance, sampling control cells from the same batch can reduce technical variation but limit the number of unique cells per set, which may limit model performance.

To construct mini-batches, we collect *B* such set pairs, 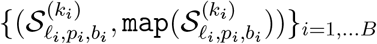, where different pairs may originate from a different combination of cell line, perturbation, and (optionally) batch. These are then arranged into the following tensors:

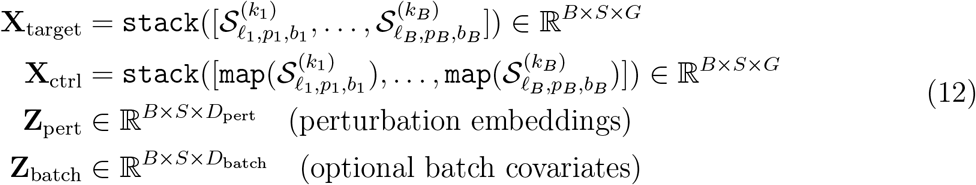

where *D*_pert_ and *D*_batch_ denote the dimensionalities of the perturbation and batch embeddings, respectively. For State, we use one-hot encodings of perturbation and batch, so *D*_pert_ equals the number of unique perturbations across the data, and *D*_batch_ equals the number of unique batch labels. ST takes **X**_ctrl_ as input along with the perturbation embeddings **Z**_pert_ and learns to predict **X**_target_ as output, learning to transform control cell populations into their corresponding perturbed states.

#### 4.3.3. Neural Network Modules

ST uses specialized encoders that map cellular expression profiles, perturbation labels, and optionally batch labels into a shared hidden dimension *d*_*h*_, which serves as the input to the transformer.

##### Control Cell Encoder

Each log-normalized expression vector **x**^(*i*)^ *∈* ℝ^*G*^ is mapped to an embedding via a 4-layer MLP with GELU activations. The MLP *f*_cell_ is applied to each cell independently across the entire control tensor:

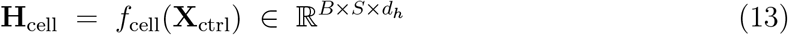

This transforms the input shape from (*B × S × G*) *→* (*B × S × d*_*h*_).

##### Perturbation Encoder

Perturbation labels are encoded into the same embedding dimension *d*_*h*_. For one-hot encoded perturbations, the input vector is passed through a 4-layer MLP with GELU activations:

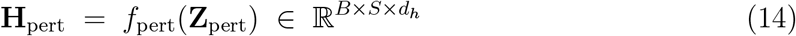

This transforms the input shape (*B × S × D*_pert_) *→* (*B × S × d*_*h*_) (note the perturbation embedding is the same for all cells within the same set of a given batch). Alternatively, when perturbations are represented by continuous features (e.g., molecular descriptors or gene embeddings), the embeddings are directly used in **H**_pert_ and we set *d*_*h*_ = *D*_pert_.

##### Batch Encoder

To account for technical batch effects, batch labels *b*_*i*_ *∈* {1, …, *B*} are encoded into embeddings of dimension *d*_*h*_:

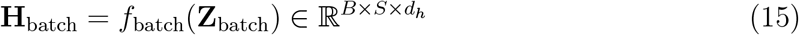

where *f*_batch_ is an embedding layer. This transforms the input shape from (*B × S × D*_batch_) *→* (*B × S × d*_*h*_).

##### Transformer Inputs and Outputs

The final input to ST is constructed by summing the control cell embeddings with the perturbation and batch embeddings:

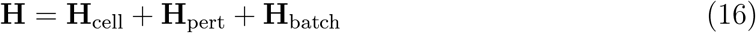

This composite representation is passed to the transformer backbone *f*_ST_ to model perturbation effects across the cell set. The output is computed as:

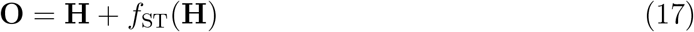

where 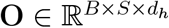 represents the final output. This formulation encourages the transformer *f*_ST_ to learn perturbation effects as residuals to the input representation **H**.

##### Gene Reconstruction Head

When working with inputs directly in expression space, the gene reconstruction head maps the output of the transformer, **O**, back to gene expression space. This is done with a linear projection layer, applied independently to the *d*_*h*_-dimensional hidden representation of each token within each cell set. Specifically, for each cell *b* in the batch, where 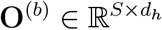 is the transformer output for cell *b*, the reconstructed gene expression 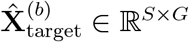 is given by:

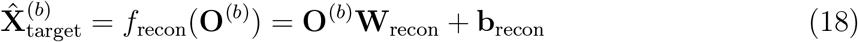

where 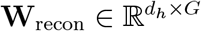 and **b**_recon_ *∈* ℝ^*G*^ are learnable parameters. This operation transforms the hidden representations, yielding reconstructed log-transformed gene expression values for each cell in the batch.

#### 4.3.4. Learning Perturbation Effects with Maximum Mean Discrepancy

ST is trained to minimize the discrepancy between predicted 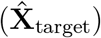 and observed (**X**_target_) transcriptomic responses. This is quantified using the Maximum Mean Discrepancy (MMD) (Gretton et al., 2012), a statistical measure of distance between two probability distributions based on their embeddings in a Reproducing Kernel Hilbert Space (RKHS), via a kernel function of choice. MMD has been applied previously to model single-cell perturbation effects (Zhang et al., 2023).

For each mini-batch element *b*, we consider the set of *S* predicted cell expression vectors of cells within the batch (therefore those sharing the same combination of cell line *ℓ* and perturbation *p*, as described in **Eq. 12**). With a slight abuse of notation, we denote this set as 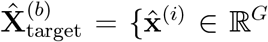 such that 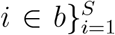, and the corresponding set of *S* target cell expression vectors of cells within the batch as 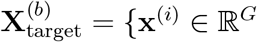 such that 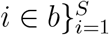. The MMD is then used to minimize the distance between the empirical probability distributions implicitly defined by these two finite sets of vectors.

The squared MMD between the predicted and observed cell sets is computed as:

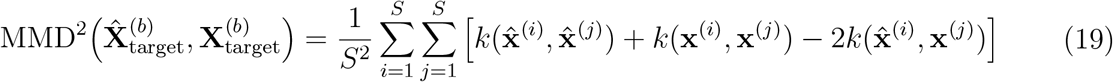

where *k*(*·,·*) denotes the kernel function. The three terms correspond to: (1) similarity within the predicted set, (2) similarity within the observed set, and (3) cross-similarity between predicted and observed sets.

We use the energy distance kernel:

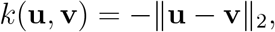

implemented via the geomloss library (Feydy et al., 2019).

For notational convenience, for a training minibatch of *B* cell sets, we define the batch-averaged MMD loss as the average of these MMD^2^ values:

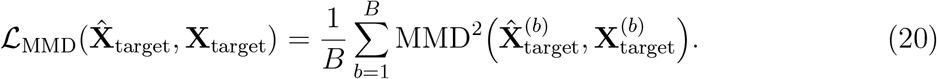

Minimizing this loss encourages the model to generate sets of perturbed cell expression vectors whose overall statistical properties, as captured by the MMD, consistent with perturbation labels and match those of the observed cell sets. The total loss is:

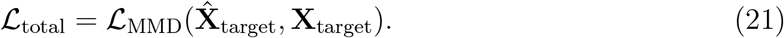

#### 4.3.5. Training ST in Embedding Spaces

ST offers the flexibility to be trained either directly in gene expression space or in a specified embedding space. When trained directly in gene expression space, ST operates on the top 2,000 highly variable genes (HVGs) as identified in our preprocessing. In this mode, the model is referred to as ST+HVG in our results. To enable training in an embedding space, the architecture is modified to include an additional expression decoder. Formally, let *E* denote the dimensionality of the embedding space, where typically *E ≪ G*. The tensors **X**_target_ and **X**_ctrl_, now denoted as 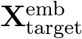 and 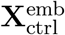, are of dimension *B × S × E*. Accordingly, *f*_cell_ is modified to transform the input shape from (*B × S × E*) *→* (*B × S × d*_*h*_), and *f*_recon_ is modified to transform the output shape from (*B × S × d*_*h*_) *→* (*B × S × E*). This output is denoted as 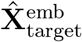. The other encoders and the transformer itself remain unchanged.

To recover the original gene expression of the target cells, **X**_target_, we train an additional decoder head *f*_decode_. This is a multi-layer MLP with dropout that maps from the embedding space back to the full gene expression space:

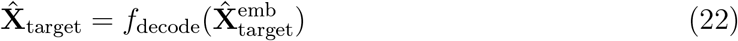

This transforms the predicted embeddings from (*B × S × E*) *→* (*B × S × G*) to recover gene expression profiles.

The ST loss is hence modified to make use of the two target sets: 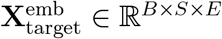 containing the target embeddings, and **X**_target_ *∈* ℝ^*B×S×G*^ containing the target gene expression profiles. Specifically, the model is trained using a weighted combination of MMD losses in the embedding and gene expression spaces:

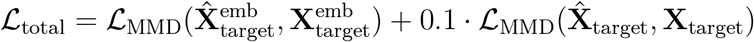

The expression loss is down-weighted by a factor of 0.1 to balance the two terms and avoid overwhelming the primary objective in embedding space. This encourages the model to learn perturbation effects primarily in the embedding space while simultaneously decoding to expression space. Our experiments demonstrate that the embedding space improves learning of perturbational effects across datasets, suggesting that the smoother structure of the embedding space facilitates better modeling of perturbation biology.

### 4.4. State Embedding Model (SE)

ST motivates the development of high-quality cell embeddings that capture relevant biological signal while reducing technical artifacts. For this, we developed the State Embedding model (SE), a self-supervised model trained with a gene expression prediction objective to learn cell representations from single-cell RNA sequencing data. The embeddings produced by SE serve as inputs to ST, enabling more robust transfer across datasets and biological contexts.

#### 4.4.1. Gene Representation via Protein Language Models

Our dataset (**Eq. 9**) consists of cell expression measurements **x**^(*i*)^, where each entry 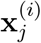 represents the normalized expression level of gene *j* in cell *i*. Beyond these expression values, the SE model incorporates rich information about genes themselves through pretrained protein language model embeddings **g**_*j*_.

Following recent work (Rosen et al., 2023; Pearce et al., 2025), we leverage pretrained protein language models to encode gene features. Gene embeddings are computed with ESM-2 (`esm2_t48_15B_UR50D` (Lin et al., 2023)), first by computing transcript embeddings by averaging per-amino-acid embeddings for each protein-coding transcript in the gene, and then by averaging across all the transcripts in the gene. This featurization captures evolutionary and functional relationships between genes. Genes without Ensembl IDs are mapped using the MyGene API (Wu et al., 2013) and Ensembl REST API (Yates et al., 2015), with manual curation for ambiguous cases. Non-protein-coding genes are excluded from SE.

Each gene embedding **g**_*j*_ *∈* ℝ^5120^ is projected into the model’s embedding dimension *h* via a learnable encoder:

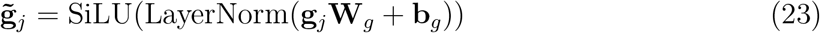

where **W**_*g*_ *∈* ℝ^5120×*h*^ and **b**_*g*_ *∈* ℝ^*h*^ are learnable parameters, and SiLU denotes the Sigmoid Linear Unit activation function (Elfwing et al., 2018).

#### 4.4.2. Cell Representation

We represent each cell *i* as a sequence of its most highly expressed genes in **x**^(*i*)^. For each cell, we construct an “expression set” by selecting the top *L* = 2048 genes ranked by log fold expression level. Empirically, increasing *L* beyond 2048 yielded diminishing returns in model performance. The expression set is then augmented with two special tokens:

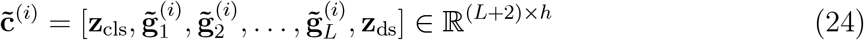

where **z**_cls_ *∈* ℝ^*h*^ is a learnable classification token used to aggregate cell-level information, and **z**_ds_ *∈* ℝ^*h*^ is a learnable dataset token that helps disentangle dataset-specific effects. Here, 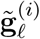 represents the projected embedding of the *ℓ*-th most highly expressed gene in cell *i*, with *ℓ ∈* {1 …, *L*}. The gene selection is cell-specific: different cells may have different sets of highly expressed genes, leading to different gene embeddings in their respective expression sets. If a cell expresses fewer than *L* genes, the expression set is padded to length *L* by randomly sampling from the pool of unexpressed genes. This maintains a fixed-length input for transformer processing.

#### 4.4.3. Expression-Aware Embeddings

Although genes in the cell expression set are sorted by expression level, their magnitudes are not explicitly encoded or used by the model. Instead, SE incorporates expression values directly using an expression embedding scheme inspired by soft binning (Hao et al., 2024). For the *ℓ*-th most expressed gene in cell *i*’s expression set (which corresponds to gene ID 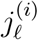) with log-normalized expression value 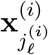 in cell *i*, we compute a soft bin assignment:

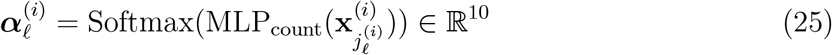

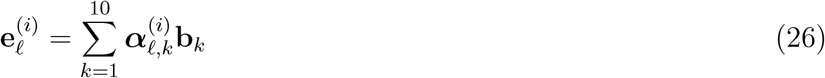

where MLP_count_ : ℝ *→* ℝ^10^ consists of two linear layers (dimensions 1 *→* 512 *→* 10) with LeakyReLU activation, and 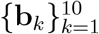 are learnable bin embeddings of dimension *h*. The resulting expression encodings 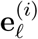 are added to the corresponding gene identity embeddings 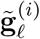:

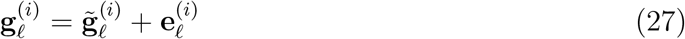

#### 4.4.4. Transformer Encoding

The input expression set, composed of expression-aware gene embeddings and special tokens, is passed through the transformer encoder *f*_SE_:

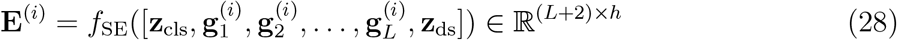

where each 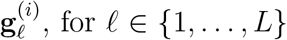 is as in **Eq. 27**. Note that the expression encodings are set to zero for the special tokens `[CLS]` and `[DS]`.

The output is a sequence of contextualized embeddings. The cell embedding is then extracted from the `[CLS]` token at position 0 and normalized:

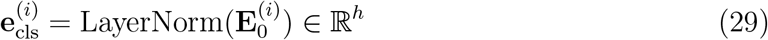

This embedding serves as a summary representation of the cell’s transcriptomic state. Similarly, we extract the dataset representation from the `[DS]` token at position *L* + 1:

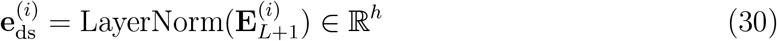

This embedding is used to capture and account for dataset-specific effects during training.

The final embedding is the concatenation of these two quantities:

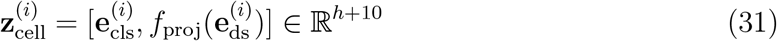

where 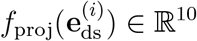 is a learned linear projection of the dataset embedding from *h →* 10. This cell embedding 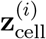 serves as the input representation for individual cells in ST, enabling the multi-scale State architecture to leverage rich cellular representations for population-level perturbation modeling.

#### 4.4.5. Pretraining Objectives

SE is trained using a self-supervised learning framework with two complementary objectives: (1) a gene expression prediction task, and (2) an auxiliary dataset classification task that helps disentangle technical batch effects from biological signal.

##### Gene Expression Prediction

During training, the model receives the complete input cell expression set (as described in **Eq. 24**), and is tasked with predicting expression values for a selected set of 1,280 genes per cell. To ensure coverage across the expression dynamic range, the target genes for prediction are drawn from three categories:

- 𝒫 ^(*i*)^, a set of |𝒫^(i)^ | = 512 highly expressed genes (from the top *L* genes in the cell expression set *i*), 𝒩 ^(*i*)^, a set of | 𝒩^(i)^ |= 512 unexpressed genes (randomly sampled from genes not in the top *L* of cell *i*),
- ℛ a set of |𝒩^(i)^ |= 256 genes randomly sampled from the full gene set, *shared for all cells in the batch*.

This results in 1,280 genes for which expression values must be predicted per cell. This strategy encourages the model to reconstruct expression values across a wide dynamic range and to learn meaningful representations of both expressed and silent genes, while having access to the complete transcriptomic context, through self attention, during prediction.

##### Expression Prediction Decoder

To learn from gene expression prediction across different datasets with varying read depth, we use an MLP decoder that combines multiple sources of information:

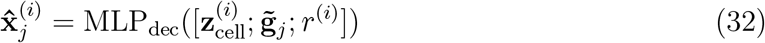

where 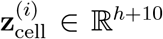 is the learned cell embedding (**Eq. 31**), 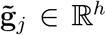 is the embedding of the target gene (**Eq. 23**), and *r ∈* ℝ is a scalar read depth indicator, computed as the mean log expression of expressed genes in the input expression set for each cell *i*. These are concatenated and passed through MLP_dec_, which consists of two skip-connected blocks followed by a linear output layer that predicts the log expression for the target gene *j* in cell *i*.

For each cell *i* in a training batch, let 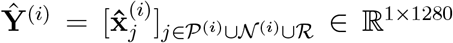 denote the row vector containing the set of predicted expression values for the genes in 𝒫 ^(*i*)^ *∪* 𝒩 ^(*i*)^ *∪* ℛ, and let 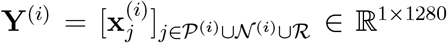 denote the row vector with corresponding true expression values. The tensors are stacked across cells in the batch: 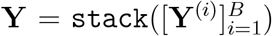 and 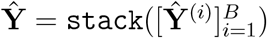, with resulting shape (*B ×* 1 *×* 1280). The gene-level loss is:

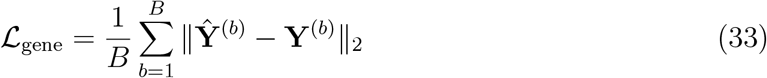

and therefore it effectively measures the similarity between predicted and true gene expression patterns within each cell *b* (which is then averaged across cells in ℒ_gene_).

To capture variation in gene expression across cells in the mini-batch, we also compute a cell-level loss using the shared subset of genes ℛ. Let 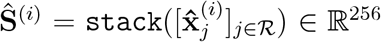 and 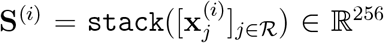 denote the predicted and true expression values in cell *i* for the shared genes ℛ. These tensors are stacked across cells in the batch and transposed: 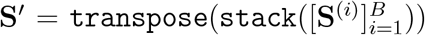 and 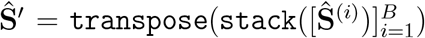, resulting in shape (|ℛ| *×* 1 *× B*), with |ℛ| = 256. We then compute the distance between the concatenated predictions and targets across all cells in the batch for each gene. Specifically, we consider the r-th gene (row) of the transposed tensors, denoted as **Ŝ**^′(*r*)^ and **S**^′(*r*)^ (each being a 1 × *B* vector):

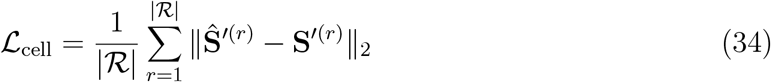

and therefore it effectively measures the similarity between predicted and true gene expression across cells in the batch for each gene in ℛ (which is then averaged across genes in ℒ_cell_).

The final training loss for expression prediction combines both axes:

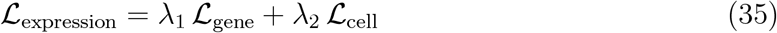

This dual-axis reconstruction loss captures both gene-wise reconstruction fidelity within each cell and the consistency of expression patterns across cells for shared genes.

##### Dataset Classification Modeling

To disentangle dataset-specific technical effects from biological variation, we introduce an auxiliary dataset prediction task. Using the `[DS]` token embedding, the model predicts the dataset of origin:

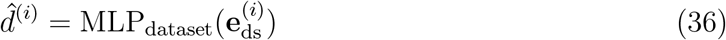

with 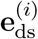 as in **Eq. 30**. We employ cross-entropy loss:

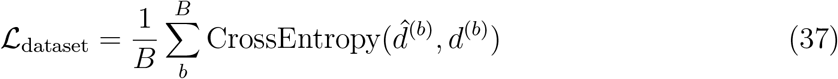

where *d*^(*b*)^ denotes the true dataset label for cell *b* in the batch, and 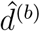 is the predicted label.

In our implementation, cells within the same AnnData file are treated as originating from the same dataset for this classification task. For our SE model trained on Arc scBaseCount, CZ CELLxGENE, and Tahoe-100M, this task becomes a multi-class classification problem over approximately 14,000 datasets. This auxiliary objective encourages the model to pool the relevant information in this token position, disentangling it from true biological signal. Empirically, since datasets in scBaseCount are organized by Sequence Read Archive experiment identifiers (SRX), which typically correspond to individual cell types or experimental conditions, the dataset token captures cell type and cell line information alongside technical batch effects. Since this token embedding is derived exclusively from the expression set representation without incorporating metadata, the trained model can be used at inference time without requiring explicit specification of technical parameters.

##### Total Loss

The SE model is trained using a combination of both losses:

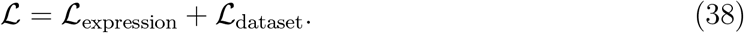

### 4.5. Datasets

#### 4.5.1. Datasets Used for ST Training

We used several single-cell perturbation datasets in this study: Tahoe-100M dataset (Zhang et al., 2025), the Replogle-Nadig dataset (Replogle et al., 2022; Nadig et al., 2025), the Parse-PBMC dataset (Parse Biosciences, 2023), the Jiang dataset (Jiang et al., 2020), the McFaline dataset (McFaline-Figueroa et al., 2019), and the Srivatsan dataset (Srivatsan et al., 2020) (Table 1). All datasets were filtered to retain measurements for 19,790 human protein-coding Ensembl genes and subsequently normalized to a total UMI depth of 10,000. Raw count data were log-transformed using `scanpy.pp.log1p`. For analyses on highly variable genes (HVGs) throughout this work, for each dataset, the top 2,000 HVGs were identified using `scanpy.pp.highly_variable_genes`. Log-transformed expression values for these HVGs were used as gene-level features. PCA embeddings of cells were computed using `scanpy.pp.pca`.

**Table 1:** Summary of datasets used in ST experiments, including total number of cells, perturbations, and biological contexts (e.g., donors, cell types, or conditions).

| Dataset | # of Cells | # of Perturbations | # of Contexts |
| --- | --- | --- | --- |
| Replogle-Nadig | 624,158 | 1,677 | 4 |
| Jiang | 234,845 | 24 | 30 |
| Srivatsan | 762,795 | 189 | 3 |
| Mcfaline-Figueroa | 354,758 | 122 | 3 |
| Tahoe-100M | 100,648,790 | 1138 | 50 |
| Parse-PBMC | 9,697,974 | 90 | 12/18 (Donors)/(Cell Types) |

#### 4.5.2. Additional preprocessing for genetic perturbation datasets

Genetic perturbation datasets were further filtered to only retain perturbations with high knockdown efficacy using the `filter_on_target_knockdown` function from the Cell-Load package. The following three filtering steps were performed:

- **Perturbation-level filtering**: Retain only those perturbations (except controls) whose average knockdown efficiency meets a minimum threshold, residual expression is *≤* 0.30.
- **Cell-level filtering**: Within the selected perturbations, keep only those cells that individually meet a stricter knockdown threshold, residual expression *≤* 0.50.
- **Minimum cell count**: Drop any perturbations that have fewer than a specified number of remaining valid cells (30), while always preserving control cells.

#### 4.5.3. Datasets Used for SE Training

SE was trained on 167 million human cells across the Arc scBaseCount (Youngblut et al., 2025), CZ CELL×GENE (Program et al., 2025), Tahoe-100M (Zhang et al., 2025) datasets (Table 2). To avoid data leakage in our context generalization benchmarks, we trained on 20 Tahoe-100M cell lines separate from the five held out cell lines from (**Fig. 2**). scBaseCount data was filtered to only retain cells with at least 1,000 non-zero expression measurements and 2,000 UMIs per cell. A subset of AnnData files were left out for computing validation loss.

**Table 2:** Summary of datasets used in SE experiments, showing total number of cells.

| Dataset | # Training Cells | # Validation Cells |
| --- | --- | --- |
| Arc scBaseCount (Youngblut et al., 2025) | 71,676,369 | 4,137,674 |
| CZ CellXGene (Program et al., 2025) | 59,233,790 | 6,500,519 |
| Tahoe-100M (Zhang et al., 2025) | 36,157,383 | 2,780,587 |

### 4.6. Training

All models are implemented using PyTorch Lightning with distributed data parallel (DDP) training. We use PyTorch’s automatic mixed precision (AMP) to reduce memory usage and accelerate training. During inference, genes not present in the embedding vocabulary are ignored.

#### 4.6.1. ST Hyperparameters

The ST architecture utilizes a shared hidden dimension, denoted as *h*, across its modules. Each encoders maps its respective inputs to this *h*-dimensional space, which also serves as the internal dimension for the transformer layers. The core transformer module, *f*_ST_, is either based on a LLaMA (Touvron et al., 2023) backbone or a GPT2 (Radford et al., 2019) backbone, provided in HuggingFace. We use a GPT2 backbone for sparser datasets, such as Replogle-Nadig (Replogle et al., 2022; Nadig et al., 2025), due to LayerNorm’s desirable centering transform operations that revive dead neurons in low-data regimes (Lyle et al., 2024). All models are modified to use bi-directional attention. Dimensionality and parameterization of the backbone used in **Fig. 2** are provided in Table 3. Since cell order within each set is arbitrary, no positional encodings are used. Additionally, dropout is not applied within the transformer. Additionally, as we work directly in vector space, we deactivate the model parameters allocated for word embeddings. Before training, most ST weights are initialized sampling from Kaiming Uniform 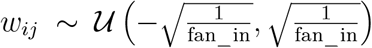 (He et al., 2015) with the exception of the transformer backbone which is initialized from 𝒩 (0, 0.02^2^). Detailed configurations for each neural network module are summarized in Table 4.

**Table 3:** Key model hyperparameters by dataset.

| Dataset | cell_set_size | hidden_dim | n_encoder_layers | n_decoder_layers | batch_encoder | transformer_backbone_key | attn_heads | params |
| --- | --- | --- | --- | --- | --- | --- | --- | --- |
| Tahoe-100M | 256 | 1488 | 4 | 4 | false | LLaMA | 12 | 244M |
| Parse-PBMC | 512 | 1440 | 4 | 4 | true | LLaMA | 12 | 244M |
| Replogle-Nadig | 32 | 128 | 4 | 4 | false | GPT2 | 8 | 10M |

**Table 4:** Architectural details for ST components. *G* denotes the number of genes (**Eq. 13**), *E* the dimension of cell embeddings (if used) (**Eq. 22**), *D*_pert_ the dimension of perturbation features (**Eq. 14**), *D*_batch_ the dimension of batch features (**Eq. 15**), and *h* the shared hidden dimension. All MLPs use GELU activation and Layer Normalization before activation, unless specified otherwise.

| Component | Architecture | Layer Dimensions | Activation | Normalization | Dropout |
| --- | --- | --- | --- | --- | --- |
| $f_{\text{cell}}$ | 4-layer MLP | $(G \text{ or } E) \rightarrow h \rightarrow h \rightarrow h \rightarrow h$ | GELU | LayerNorm | None |
| $f_{\text{pert}}$ | 4-layer MLP | $D_{\text{pert}} \rightarrow h \rightarrow h \rightarrow h \rightarrow h$ | GELU | LayerNorm | None |
| $f_{\text{batch}}$ | Embedding Lyr. | $D_{\text{batch}} \rightarrow h$ | N/A | N/A | None |
| $f_{\text{ST}}$ | LLaMA Transf. | $h$ (input to each of 4 layers) | SwiGLU | RMSNorm | None |
| $f_{\text{recon}}$ | Linear Layer | $h \rightarrow G \text{ or } E$ | N/A | N/A | None |
| $f_{\text{decode}}$ | 3-layer MLP | $h \rightarrow 1024 \rightarrow 512 \rightarrow G$ | GELU | LayerNorm | 0.1 |
| $f_{\text{conf}}$ | 3-layer MLP | $h \rightarrow h/2 \rightarrow h/4 \rightarrow 1$ | GELU | LayerNorm | None |

#### 4.6.2. ST Training Details

All components are trained end-to-end with using the objectives described previously. These components include the control cell encoder *f*_cell_, perturbation encoder *f*_pert_, optional batch encoder *f*_batch_, transformer backbone *f*_ST_, reconstruction and decoding heads *f*_recon_ and *f*_decode_.

For the fine-tuning tasks (**Fig. 3G**), we initialize a new ST model using the pretrained weights while selectively reinitializing specific components that are dataset-specific. In particular, the perturbation encoder *f*_pert_ is reinitialized to enable transfer across perturbation modalities. If ST is trained using cell embeddings, the gene decoder *f*_decode_ is also reinitialized to adapt to differences in gene coverage between datasets. Since different experimental platforms and datasets typically measure distinct subsets of genes, the decoder must be retrained to map from the shared embedding space to the target dataset’s specific gene expression space. All other components — *f*_cell_, *f*_batch_, and *f*_ST_ — retain their pretrained weights and are finetuned on the target dataset.

#### 4.6.3. SE Hyperparameters

SE is a 600M parameter encoder-decoder model designed to learn cell representations by predicting gene expression variability. The encoder consists of 16 transformer layers, each with 16 attention heads and hidden dimension *h* = 2048. Each layer uses pre-normalization with a feed-forward network that expands to dimension 3 *× h*, and uses GELU activation. We apply dropout with probability 0.1 to both attention and feed-forward layers. The decoder is a multi-layer-perceptron (MLP), trained to recover gene expression given learned cell embeddings and target gene embeddings. We used the AdamW optimizer (Loshchilov and Hutter, 2017) with a maximum learning rate of 10^−5^, weight decay of 0.01, and gradient clipping with zclip (Kumar et al., 2025). The learning rate schedule consisted of linear warmup for the first 3% of total steps, followed by cosine annealing to 30% of the maximum learning rate. Before training, all SE weights are initialized sampling from Kaiming Uniform.

#### 4.6.4. SE Training Details

SE was trained on a large-scale corpus of 14,420 AnnData files spanning 167 million human cells across the Arc scBaseCount (Youngblut et al., 2025), CZ CELL×GENE (Program et al., 2025), and Tahoe-100M (Zhang et al., 2025) datasets for 4 epochs. To avoid data leakage, datasets were split into separate training and validation sets at the dataset level. To enable efficient training at scale, we utilize Flash Attention 2 (Dao, 2024), with mixed precision (bf16) training (Kalamkar et al., 2019). Training was distributed across 4 compute nodes, each with 8 NVIDIA H100 GPUs. The model was trained with an effective batch size of 3,072, using per-device batch size of 24 and gradient accumulation over 4 steps.

### 4.7. Evaluation

To comprehensively assess the ability of State to model perturbation effects, we designed an evaluation framework, Cell-Eval, that captures both expression-level accuracy and biologically meaningful patterns of differential expression. Cell-Eval measures not only statistical performance but also the model’s utility in simulating realistic Perturb-seq experiments.

#### 4.7.1. Perturbation Evaluation Metrics

A key goal for perturbation models is to distinguish between different perturbation effects.

Cell-Eval evaluates this using several complementary metrics:

##### Perturbation Discrimination Score

Adapted from Wu et al. (2024), this metric ranks predicted post-perturbation expression profiles by their similarity to ground truth. The score is defined as the normalized rank of the ground truth from the predicted perturbation with respect to all ground truth perturbations. It directly evaluates whether models can recover the relative differences between perturbations. In Wu et al. (2024), the rank ordering is computed using cosine similarity; we compute using Manhattan distances of transcriptomes due to two factors: larger sensitivity to magnitudes (**Fig. S10**) and the lack of normalization in our loss (Steck et al., 2024). Specifically, let *T* be the number of distinct perturbations, *y*_*t*_ the ground-truth profile for perturbation *t*, and *ŷ*_*t*_ its predicted profile. Using a distance *d*(*·, ·*) (here Manhattan or Euclidean), define

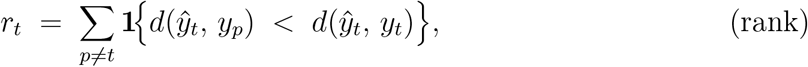

i.e. the number of other perturbations whose ground-truth profile is closer to *ŷ*_*t*_ than the correct one. The per-perturbation score is

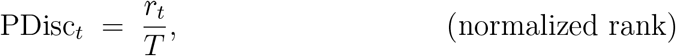

ranging in [0, 1) with PDisc_*t*_ = 0 indicating a perfect match (no closer profiles) and values near 1 signaling poor discrimination. The overall score is the mean

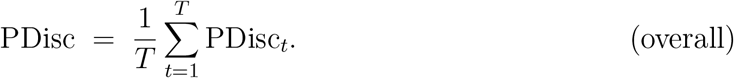

We report the normalized inverse perturbation discrimination score:

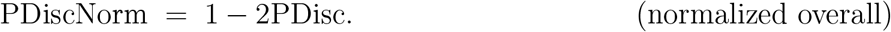

so that a random predictor receives a score of 0.0, and a perfect predictor receives a score of 1.0.

##### Pearson Delta Correlation

This metric calculates a Pearson correlation coefficient between the predicted and observed expression deltas. The expression delta is computed as the difference in transcriptomes between a perturbed pseudobulk and an unperturbed pseudobulk. Specifically, for each perturbation *t* we form a pseudobulk 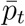 and a control 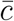 and compute the expression delta: 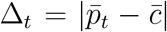, the element-wise absolute difference between the mean expression of perturbed cells 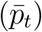 and control cells 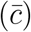. The evaluation score is the Pearson correlation

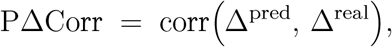

computed with `scipy.stats.pearsonr` across all perturbations.

##### Adjusted Mutual Information (AMI)

AMI is used to assess how well the model preserves the clustering structure of perturbations in the learned embedding space. We compute AMI by first aggregating cells by perturbation, then computing centroids for each perturbation by averaging the embeddings of all associated cells in both the real and predicted datasets. We then perform Leiden clustering on these centroids and compare the predicted cluster labels to the true cluster labels using AMI. The maximum AMI score across all resolutions is reported as the final metric, reflecting the degree to which the model’s predicted perturbation space captures the global structure of biological perturbations. Where ℛ is the set of Leiden clustering resolutions, and 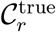 and 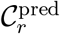 are the clusterings of real and predicted centroids at resolution *r*,

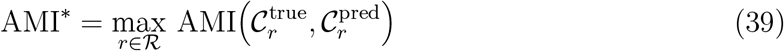

##### Mean Absolute Error (MAE) & Mean Squared Error (MSE)

To assess preservation of shifts we calculate the MAE and MSE of each perturbation between the predicted and ground truth datasets. The MAE/MSE metrics are evaluated over predicted and observed pseudobulks. Formally, for perturbation *t*, let 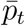 and *p*_*t*_ denote the predicted and observed pseudobulked expressions, respectively. Then,

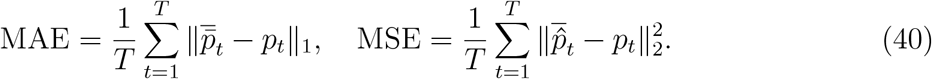

#### 4.7.2. Differential Expression

To evaluate biological relevance, Cell-Eval performs a differential expression analysis using the Wilcoxon rank-sum test and adjusts for multiple hypotheses using the Benjamini-Hochberg procedure, applied to both observed values and model predictions in test cell lines.

##### Notation

Fix a perturbation *t ∈* [*T*] = {1, …, *T*} and let *G* be the set of all genes. Define 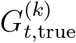 and 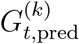 as the top-*k* significant DE genes (*p*_adj_ *<* 0.05) in ground truth and predictions, ranked by log-fold changes 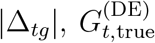 and 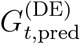 as the complete significant sets. All rank correlations are Spearman’s, denoted by *ρ*_rank_.

Cell-Eval also evaluates model performance using many complementary metrics:

##### DE Overlap Accuracy

For each perturbation, we identify the top *k* differentially expressed genes (*k* = 50, 100, 200, *N*), filtered by adjusted *p*-value and ranked by absolute log fold change. We compute the intersection between the predicted and true DEG sets and report the overlap as a fraction of *k*. When *k* = *N* the *k* value varies by perturbation where *N* is the total number of differentially expressed genes in the true DEG set. When *k* is not specified, we are referring to DE Overlap at *N*. That is,

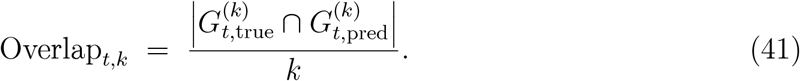

##### Top-*k* precision

For each perturbation, we compute how many of the top *k* DEGs from the ground truth appear in the model’s top *k* predicted DEGs, measuring precision at various thresholds. That is,

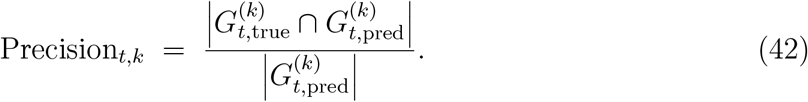

##### Directionality Agreement

For each perturbation, we identify the set of significantly differentially expressed (DE) genes in the ground truth using an adjusted p-value threshold (e.g., *p <* 0.05). We then find the intersection between the predicted and observed DE gene sets. For each overlapping gene, we check whether the predicted and true fold changes have the same direction. The directionality agreement is defined as the fraction of overlapping genes for which the predicted and observed directions match. Formally, where 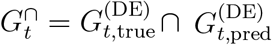 and where log-fold changes Δ_*tg*_ (true), 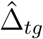 (predicted). That is,

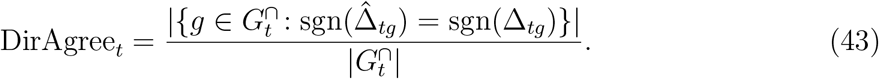

##### Spearman Correlation

We compute the Spearman rank correlation between predicted and observed fold changes, restricted to genes that are significantly DE in ground truth. For each perturbation, we extract the list of significant genes (based on the adjusted p-value threshold) and compute the Spearman correlation coefficient on the log fold-changes between predicted and ground truth. That is,

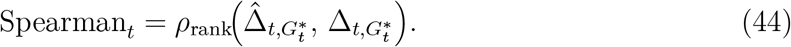

##### ROC-AUC

To assess the model’s ability to distinguish significant from non-significant DE genes, we assign binary labels to each gene in the observed data (1 if *p <* 0.05, 0 otherwise) and use predicted − log_10_ (adjusted p-values) as confidence scores. We compute the true positive rate (TPR) and false positive rate (FPR) at multiple thresholds and calculate the area under the ROC curve using auc. This metric evaluates the model’s ability to separate significant from nonsignificant genes regardless of the classification threshold. That is,

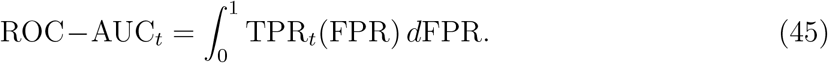

##### PR-AUC

To measure the precision recall tradeoff of the model in identifying significant DE genes, as in ROC-AUC, we use binary labels from the observed and predicted − log_10_(p-values) as scores. This metric reflects the model’s ability to recover true positives with high precision when significant DE genes are sparse. That is,

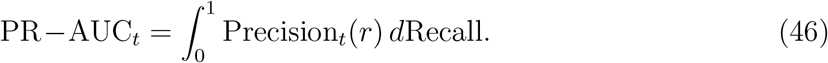

##### Effect sizes

To compare the relative effect sizes of perturbations we calculate Spearman correlation coefficients for each perturbation on the number of differentially expressed genes (adjusted p-value *<* 0.05) between predicted and ground truth. This lets us assess whether models accurately capture relative effect sizes. Formally, for set sizes 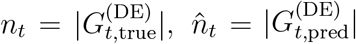, we compute

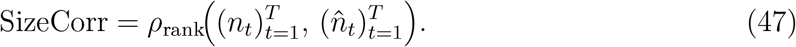

#### 4.7.3. Cell Embedding Evaluation Metrics

We also evaluated cell embeddings using both intrinsic and extrinsic metrics (**Fig. 3D**).

##### Intrinsic Evaluation

To assess the extent to which embeddings capture perturbation-specific information, we trained a multilayer perceptron (MLP) with one hidden layer to classify perturbation labels from cell embeddings, measuring AUROC and accuracy. This classification task was performed separately for each cell line, with data partitioned such that 20% of cells from each perturbation group were randomly assigned to the test set and the remaining 80% to the training set. The MLP was trained using a cross-entropy loss. High classification performance indicates that embeddings maintain distinct representations of perturbation-induced states.

##### Extrinsic Evaluation

To evaluate the downstream utility of various cell embeddings (**Fig. 3E**), we compared the performance of ST when trained with different embedding spaces. This approach captures the idea that embeddings rich in cellular information may not always be optimal for modeling perturbation effects. The ideal embedding space should balance both: it should preserve perturbation-specific information while supporting accurate prediction of perturbation-induced transitions across cell states.

#### 4.7.4. Other Evaluations

##### Drug Similarity Analysis

To evaluate how well different models preserved the relational structure between drug perturbations, we first constructed a representation of each drug as a vector of log_2_-fold changes in gene expression across cell lines. These vectors were aligned to a shared gene *×* cell line space. Pairwise drug-drug similarities were then calculated using cosine similarity. To assess the structural preservation of perturbations, we applied hierarchical clustering with average linkage to both the real and predicted distance matrices and computed the Adjusted Rand Index (ARI) between the resulting cluster assignments (using *k* = 5 clusters).

##### Cell Set Scaling

To evaluate the impact of cell set size in State, we ablated the number of cells per set and the batch size, such that the batch size times the cell set size was always 16,384, and measured the validation loss as a function of floating-point operations (FLOPs) (**Fig. 1C**). These models were also compared to a pseudobulk model that replaces the self-attention in State with mean pooling, and masked attention version of State, where each cell can only attend to itself. As the cell set size increased, the validation loss improved significantly on held out cell lines, up to an optimal value of 256.

##### Survival Prediction Analysis

In order to assess whether the model could be used to predict the survival of each cell line within each drug well, a previously used measure of fitness (Yu et al., 2024), we built a model to predict survival based on the mean expression of the 2K HVG set relative to DMSO for each cell type. Specifically, for the held-out set of cell lines and perturbations, we calculated a vector of true gene expression differences between DMSO and the perturbation. Then, we trained a support vector machine model to predict survival values based on these vectors, with all 3 doses from 35 randomly selected drugs held out. Finally, we used the same SVM model to predict the survival for these held-out treatments, first predicting on the true gene expression counts, followed by the predicted, cell-type mean, and pert-mean. In order to calculate confidence intervals, this process was repeated 10 times, with a different set of drugs being held out each time.

The SVM model was a default scikit-learn support vector regressor model, with an “rbf” kernel. Relative fitness for each condition was calculated as 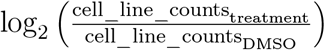, where DMSO is matched to be in the same plate as the treatment.

#### 4.7.5. Baseline Models

We describe and formalize various baseline models used as comparisons to State.

##### Perturbation Mean Baseline

This baseline predicts a perturbed expression profile as the control mean for that cell context plus a global perturbation offset learned from the training data. For each cell type *c* and perturbation *p* we first compute cell-type–specific means

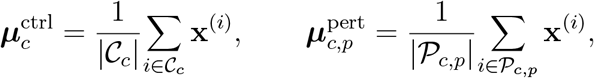

where 𝒞_*c*_ is the set of control cells of type *c* and 𝒫_*c,p*_ the perturbed cells of type *c* receiving perturbation *p*. Their difference is the cell-type offset 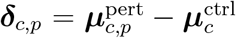. Averaging across all cell types that contain *p* yields a global offset

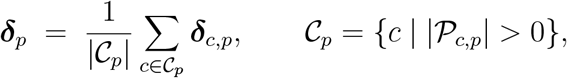

Given a test cell type *t* and a perturbation label *p* the model outputs

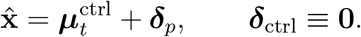

Thus, controls are reproduced exactly, while every non-control perturbation receives the same global shift.

##### Context Mean Baseline

This baseline predicts a cell’s post-perturbation profile by returning the average perturbed expression of *cells of the same cell type* observed in the training set. For every cell type *c* we collect all training cells whose perturbation is not the control and form the pseudo-bulk mean

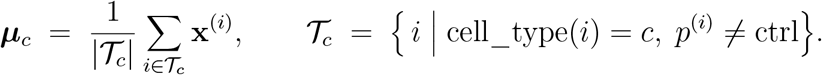

At inference time, for a test cell *i* with cell type *c*^(*i*)^ and perturbation label *p*^(*i*)^ we predict

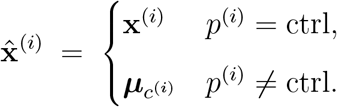

i.e. controls are passed through unchanged, whereas perturbed cells inherit their cell-type mean.

##### Linear Baseline

This baseline treats a perturbation as a low-rank, gene-wide linear displacement that is added to each cell’s own control expression (Ahlmann-Eltze et al., 2024). Let 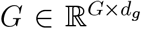 be a fixed gene-embedding matrix (e.g. pretrained protein feature vectors, one row per gene) and 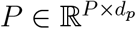 a fixed perturbation-embedding matrix (one row per perturbation, one-hot). From the training set we first build an “expression-change” pseudo-bulk

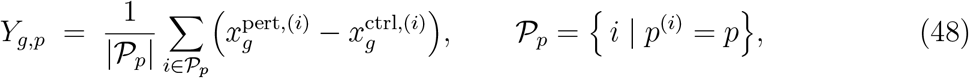

so *Y ∈* ℝ^*G×P*^ stores, for every gene *g* and perturbation *p*, the average change relative to that cell’s matched control. The model seeks a low-rank map 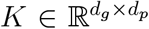 and a gene-wise bias **b** *∈* ℝ^*G*^ such that

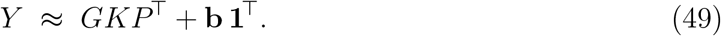

We obtain *K* in a single shot by solving the ridge-regularized least-squares problem

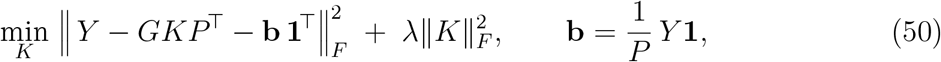

whose closed-form solution is

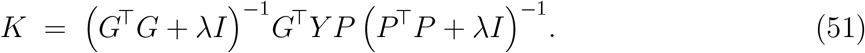

No gradient-based optimization is required, as once *K* is computed the model is fixed. For a test cell *i* with its own control profile **x**^ctrl,(*i*)^ and perturbation label *p*^(*i*)^ the prediction is

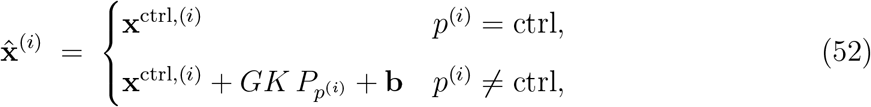

where 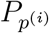 denotes the row of *P* corresponding to perturbation *p*^(*i*)^. Thus each prediction preserves the cell’s basal state exactly and adds a perturbation-specific, low-rank shift learned from the training cohort, with model capacity controlled solely by the embedding dimensions and the ridge parameter *λ*.

##### Deep Learning Baselines

We benchmark State against several baselines that leverage related deep learning architectures for predicting perturbation effects across diverse cell contexts. These include two autoencoder-based models, scVI (Lopez et al., 2018) and CPA (Lotfollahi et al., 2023), as well as the transformer-based scGPT model (Cui et al., 2024). scVI models gene expression distributions while accounting for technical noise and batch effects. CPA learns a compositional latent space that captures the additive effects of perturbation, dosage, and cell type. scGPT leverages generative pretraining on over 33 million cells to support zero-shot generalization across tasks including perturbation prediction. All models are evaluated using the respective datasets and setup described in **Fig. 2B**.

## 5. Analysis: State Transition as Optimal Transport

In this section, we analyze the theoretical capacity of State Transition (ST) in learning optimal transport mappings across cellular distributions. First, we prove that the solution family of ST covers the optimal transport map between control and perturbed cell distributions. Second, we derive explicit constraints on ST for learning the unique continuous optimal transport map. Finally, we discuss the implications of the theoretical analysis and possible future directions.

### 5.1. ST Asymptotic Behavior and Solution Family

Optimal Transport (OT) provides a framework for comparing and aligning probability distributions by finding a mapping or coupling that minimizes a certain fixed cost. Traditional OT methods explicitly define a cost function and solve a linear program or iterative algorithm to find an optimal coupling or map (Cuturi, 2013; Peyré et al., 2019). The advent of deep learning has given rise to Neural Optimal Transport (Neural OT), which leverages neural networks to parameterize and solve OT problems (Bunne et al., 2024b, 2023).

Neural OT methods (Bunne et al., 2024b) typically operate by either: (1) Parameterizing the OT map *T* directly using a neural network designed to enforce OT properties; (2) Parameterizing the OT coupling matrix (**P**) or its dual potentials (e.g., *α, β*) with neural networks; (3) Explicitly minimizing an OT cost function (or its dual objective) as the primary loss. For instance, CellOT (Bunne et al., 2023) explicitly parameterizes the convex potentials of the dual optimal transport problem using Input Convex Neural Networks (ICNNs) to learn an optimal transport map *T*_*k*_ = *∇g*_*k*_ for each perturbation *k*, with *g*_*k*_ one of the dual potentials.

ST, while not explicitly solving an OT problem in the traditional sense, performs a task related to Neural OT: learning a transformation that aligns unperturbed and perturbed cell distributions. This transformation is learned, not explicitly engineered for a fixed cost. However, motivated by rigorous theoretical work that demonstrates how transformers with fixed parameters can provably solve optimal transport by implementing gradient descent through engineered prompts (Daneshmand, 2024), we demonstrate that, in an asymptotic setting, the solution family of our trained and optimized ST contains the unique continuous optimal transport map between cellular distributions when regularity assumptions on the distributions are met.

For this analysis, we focus on a single training mini-batch, denoted as the *b*-th batch. We start by describing the mathematical formulation of ST, which aims to learn the effect of one single perturbation from cell population 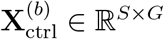 to 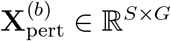. Recall that the input of the transformer for this batch is constructed by summing the control cell embeddings with the broadcasted perturbation and batch embeddings, 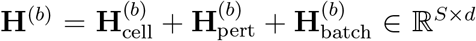 (**Eq. 16**). The output of the transformer is computed as **O**^(*b*)^ = **H**^(*b*)^ + *f*_ST_**H**^(*b*)^ *∈* ℝ^S×d^, where *f*_ST_ is the transformer encoder (**Eq. 17**). We consider *f*_ST_ with *L* layers, where the input to the first layer is **H**^0^ = **H**^(*b*)^:

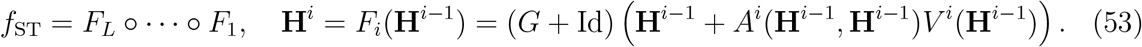

Here *A*^*i*^ denotes the multi-head attention, *V* ^*i*^ denotes the attention value encoder, and *G* + Id denotes the feedforward layer combined with the residual connection. The layer normalization function is absorbed in each constructed function. Finally, for the *b*-th batch, a cell-wise linear transform defines the final output 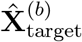 for the *S* cells (**Eq. 18**). We can equivalently rewrite this for each cell *s* as:

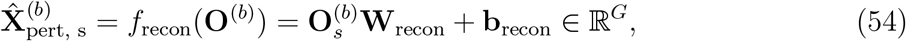

where 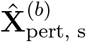 denotes the *s*-th row of 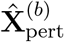, containing the predicted state of cell *s*, obtained by its embedding 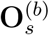 which denotes the *s*-th row of **O**^(*b*)^.

The constants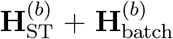, as well as the transforms *f*_cell_, *f*_recon_ can be absorbed in the first and last layers of *f*_ST_, which we therefore define as 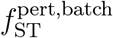. Therefore, we can equivalently rewrite the predicted state of cell *s* as:

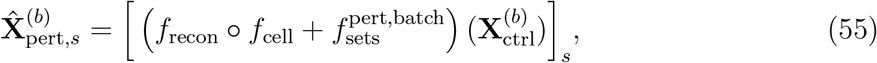

where [*·*]_*s*_ indicates that we access the *s*-th row of the obtained output. To simplify our analysis, we focus on the asymptotic setting where the cell set size *S* tend to infinity. Recall that each cell is sampled from the distribution 𝒟_ctrl_. Then, as the cell number *S → ∞* with certain regularity conditions, the output for a single cell 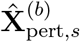 depends solely on 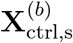 itself and the overall distribution 𝒟_ctrl_, because the attention mechanism effectively processes information from the entire distribution. Thus, the limiting operator can be defined per cell as:

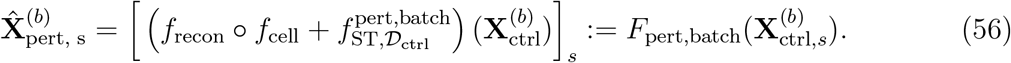

More detailed treatments are provided in recent works considering the limiting mean-field dynamics of transformers (Furuya et al., 2024; Biswal et al., 2024). Notably, **Eq. 56** holds for an arbitrary cell *s* sampled from the distribution 𝒟_ctrl_. The operator **F**_pert,batch_ defines a mapping from 𝒟_ctrl_ to 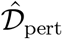. We further assume that the transformer model is expressive enough, such that minimizing the empirical kernel MMD objective is possible: 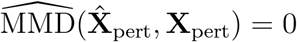 with energy kernel *k*(*u, v*) = − *∥u* − *v∥* _2_. Our following lemma shows that zero empirical MMD indicates distributional matching with probability 1.

#### Lemma 1

(Distributional matching via Empirical MMD). *Suppose that the supports of* 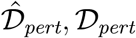 *are bounded. When the cell number S → ∞, if the empirical MMD reaches zero, that is*, 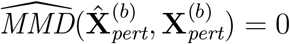, *then with probability 1, we have*

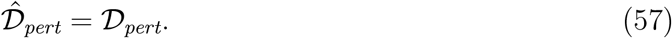

*Reversely, if* 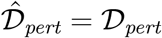, *then with probability 1, we have* 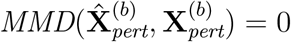.

*Proof*. The empirical MMD used as the ST objective is defined as

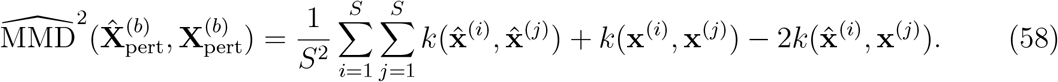

Here we use 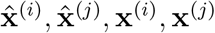 to denote cells in 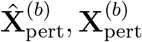 respectively. The corresponding theoretical MMD on distributions 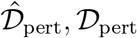 is defined as

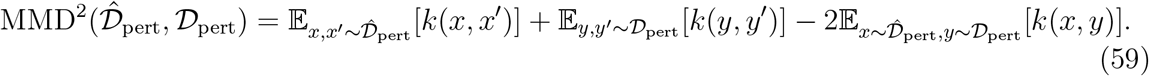

The empirical MMD is a V-statistic (Gretton et al., 2012), thus when the support sets are bounded (thus the kernel value *k* is also bounded), by the result of SLLN for V-statistics (Hoeffding, 1961; De la Pena and Giné, 2012), we have

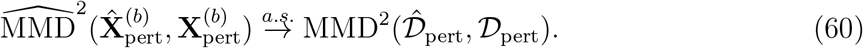

With the energy kernel *k*(*x, y*) = −*∥x* − *y∥* _2_, the theoretical MMD coincides with the energy distance *D*^2^ between distributions:

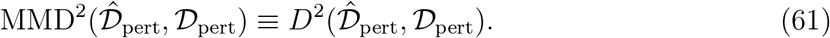

Finally, zero energy distance implies equal distributions and vice versa (Rizzo and Székely, 2016). □

We define the continuous solution families that achieves zero empirical MMD and distributional matching as

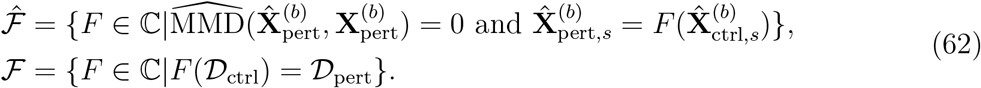

Then, by Lemma 1, we have

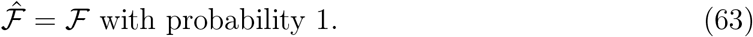

To simplify our analysis, we assume that our model is expressive enough to learn any element in 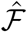, which is identical to ℱ with probability 1. We next demonstrate that, under mild regularity conditions, the unique optimal transport mapping from 𝒟_ctrl_ to 𝒟_pert_ is within the solution family 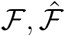.

#### Theorem 2

(Optimal Transport Mapping within the Solution Family of State). *Assume the densities of* 𝒟_*ctrl*_, 𝒟_*pert*_ *are absolute continuous and bounded, and the support sets are strictly convex and compact with ℂ* ^2^ *boundary. Then the continuous optimal transport map T from* 𝒟_*ctrl*_ *to* 𝒟_*pert*_ *associated with the squared distance cost c*(*x, y*) = *∥x* − *y∥*^2^ *satisfies* 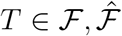 *with probability 1*.

*Proof*. Under the regularity conditions posed for distributions 𝒟_ctrl_, 𝒟_pert_, applying Caffarelli’s theorem (Caffarelli, 1992) shows that there exists an continuous and differentiable optimal transport map *T ∈* ℂ^1^ between 𝒟_ctrl_, 𝒟_pert_. Thus by definition, *T ∈* ℱ. Furthermore by Lemma 1, 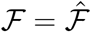 with probability 1, in which case we also have 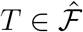. □

While the theorem shows the existence of the continuous optimal transport map *T* and that it is covered by the solution family 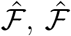 contains infinite additional elements beyond *T*, thus the optimal transport solution is not guaranteed to be learned by the model. This inherent flexibility is advantageous for State, as biological transformations often involve complexities and unmodeled variations that may not align perfectly with a single, fixed optimal transport solution. Nevertheless, in the next theorem, we show that if we impose constraints on the Jacobian in the ST model objective, then the optimal transport mapping is guaranteed to be learned:

#### Theorem 3

(Constrained ST Model for Unique OT Map). *Under the same assumptions as Theorem 2, consider the constrained solution family:*

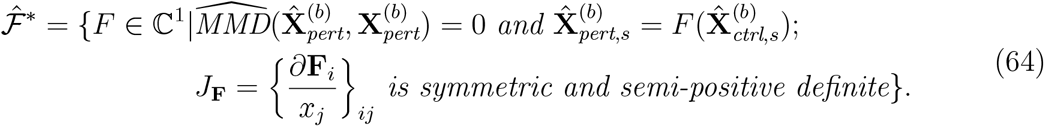

*Then* 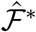 *uniquely contains the continuous OT mapping T with probability 1:* 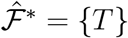.

*Proof*. By Breiner’s theorem (Brenier, 1991), *T* is the unique continuous optimal transport mapping regarding the quadratic cost, if and only if it is the gradient of a convex function *ψ*: *T* = *∇ψ*. As we already have *T ∈* ℂ^1^ by Caffarelli’s theorem, the condition is equivalent to symmetric and semi-positive definite Jacobian in *T*. Further by Lemma 1, 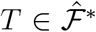 with probability 1, and the uniqueness of *T* implies 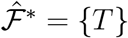. □

Our theoretical analysis can be seen as a corollary of Caffarelli and Brenier’s theorems on continuous optimal transport maps with quadratic cost. In particular, the final theorem hints that, we can enforce the model to learn an optimal transport mapping from 𝒟_ctrl_ to 𝒟_pert_, through imposing adequate regularizations on the Jacobian of the mapping. More precise constraints on the Jacobian may require additional architecture designs that explicitly achieve Jacobian constraints, or directly retrieve gradients of convex neural networks (Makkuva et al., 2020; Bunne et al., 2023).

Despite the large feasible solution set that contains numerous overfitting solutions, neural network models have shown remarkable generalization capacities. An important aspect on neural network generalization capabilities is implicit bias (Soudry et al., 2018; Gunasekar et al., 2018; Vardi, 2023), which means gradient descent favor certain solutions, for instance, those minimally shifted compared with initial weights (Gunasekar et al., 2018). Therefore we hypothesize that the optimization of ST may lead to solutions with smallest “overall shifts” from 𝒟_ctrl_ to 𝒟_pert_, thus resembling a optimal transport mapping with some unknown cost function. Theoretical characterization of the implicit bias in the transition model is out of the scope here and remains promising future research.

## 6. Exploring the Latent Space of State

Initial interpretability analyses reveal that State’s predictions can be better understood through its internal computation patterns, particularly suggesting that the transition model architecture harnesses heterogeneity to enhance perturbation prediction.

### State embeddings display markers of biological knowledge learned from self-supervision

We begin by evaluating cell embeddings produced by SE. Given the embedding model’s impressive performance in a zero-shot setting across datasets, it is instructive to examine State’s embedding structure. As shown in **Fig. S9A**, the principal components of State embeddings capture nuanced distinctions across and within cell types, suggesting a rich representation for transfer. Notably, embeddings for K-562 and Jurkat (two leukemia cell lines) cluster closely in this space, implying similarity under State’s learned representation, following results in Nadig et al. (2025). Importantly, SE was trained without direct knowledge that K-562 and Jurkat share these similarities, and recognizes and leverages such biological concepts purely from self-supervision over gene expression data.

### The State transition model leverages cell heterogeneity through self-attention

Early efforts to reverse-engineer deep machine learning models have commonly focused on visualizing activations, revealing how neural networks encode concepts (Zeiler and Fergus, 2014; Olah et al., 2017; Bau et al., 2017; Clark et al., 2019). Similarly to these works, since all inputs to ST originate from the same cell line, we aim to unpack the model’s inner workings with respect to cell heterogeneity. To accomplish this, norm-based contribution magnitudes and self-attention maps for well-defined cell sets can discover general behaviors in the model’s methods for prediction. Running inference with the Replogle-Nadig dataset, we first find that the largest changes to hidden state happen at the final layer of the transition model, across all attention heads (**Fig. S9B**). A deeper dive into these heads reveals intriguing structural behaviors. We find that the attention heads on the final layer take on extremely different roles. Specifically, we identify major two classes of heads:

1. **Heterogeneity-Sensitive heads**, which exhibit scattered, high-magnitude Query-Key interactions across many cells or only focus on the corresponding Key matrix for the Query (**Fig. S9C&D**).
2. **Heterogeneity-Insensitive heads**, which fixate on either narrow Key matrices, often from just a few cells, and broadly apply across all Queries (**Fig. S9C&D**). These behaviors mimic operations akin to pseudobulking (i.e. averaging across single-cell profiles to mimic bulk measurements) or PerturbMean-style computations, two robust but less expressive baselines to State.

To further dissect heterogeneity-sensitive attention behaviors, we construct an inference-time example with a mixed cell set composed of 50% K-562 and 50% Jurkat cells (**Fig. S9D**). Upon inputting this hybrid set, we observe that certain attention heads adaptively shift their behavior: heterogeneity-aware heads become more polarized, and a novel insensitive head emerges, one that selectively utilizes only the Key matrices from one cell type while computing over Queries from both. This asymmetric attention pattern reveals that State is not merely averaging across cell sets, but actively discriminating between them in a context-sensitive fashion. Such computations suggest that the transition model internalizes representations of heterogeneity and dynamically leverages them during inference for improved prediction performance.

These initial results underscore the interplay of leveraging transformer interpretability to better understand perturbation effects in cells. Future work can apply more sophisticated interpretability methods—beyond attention maps, which have shown limited utility for exact localization (Hase et al., 2023; Chefer et al., 2021)—to better understand State’s prediction mechanisms, building on approaches that optimize for extracting sparse, finer-grained model features (Bricken et al., 2023; Bhaskar et al., 2024; Dunefsky et al., 2024; Ameisen et al., 2025).

## Code and model availability

All code for this project is available at https://github.com/ArcInstitute/State, https://github.com/ArcInstitute/Cell-load, https://github.com/ArcInstitute/Cell-eval, https://github.com/ArcInstitute/State-reproduce.

Model parameters are available on Huggingface: https://huggingface.co/arcinstitute/SE-600M, https://huggingface.co/arcinstitute/ST-Tahoe, https://huggingface.co/arcinstitute/ST-Parse

## Acknowledgments

We thank Joseph Caputo, Alden Woodrow, Julia Kazaks, Jingtian Zhou, Brian Hie, Tony Hua, Paul Datlinger, Nianzhen Li, Brian Yu, David Lara-Astiaso, Kristen Seim, Lorena Saavedra, Rachel Senturia, Scott Newins, Gwanggyu Sun, Tim Hudelmaier, Jason Swinderman, for helpful discussions and assistance with manuscript preparation. We also thank Charlotte Bunne, Michael Bereket, Ansh Khurrana, Ayush Agrawal, Zane Dash, Yanay Rosen and Minkai Xu for helpful discussions and experiments during the early stages of this project. This study was supported by Arc Institute. We also acknowledge the efforts of our colleagues to generate and release large-scale datasets necessary to train and evaluate predictive models.

## Author contributions

A.K.A and Y.H.R. conceived the State project. P.D.H. and S.K. conceived Arc’s Virtual Cell Initiative. Y.H.R., H.G, D.P.B, A.D, P.D.H. and S.K. supervised the project, with Y.H.R. coordinating effort across the team. Y.H.R. designed the machine learning problem. A.K.A. and Y.H.R. developed the ST approach; A.K.A. implemented the data loaders, ST model, and scaling for SE pretraining. A.K.A., D.G., and Y.H.R. iteratively improved the ST architecture; A.K.A., A.I., R.S., R.I., and Y.H.R. improved the SE architecture. N.T. combined ST and SE into a unified State code base. Y. H. R., N.T., A.D., S.K. and B.E. designed evaluation metrics. M.N., S.N. designed and ran baseline perturbation models. D.G. analyzed ST self-attention and optimized metrics and data loaders. B.B. developed the initial theoretical formulation, which M.D. formalized to develop the proof showing that ST can learn optimal transport. B.B, D.G formalized model description. A.I. curated the pretraining dataset for SE and conducted training experiments for both SE and ST, including scaling experiments, pretraining for ST, zero-shot and few-shot evaluations, and biological context generalization. R.S helped curate the pretraining dataset for SE and evaluate SE embeddings. N.T. wrote the evaluation suite. Y.H.R. performed the cell type specificity DE analysis. C.C conducted a survival analysis. V.S. conducted an analysis on drug-drug similarity. A.W. conducted mutant analysis to find cell type specific perturbation effects. S.T. optimized State code for multi-node training. J.S. helped with multi-node training and setting up the compute infrastructure. N.Y. designed a web portal. C.R.T created visuals for main text figures. J.L and L.A.G. provided feedback on experiments and problem formulation. Y.H.R. designed the figures and initial manuscript outline, and Y.H.R., H.G., D.P.B. wrote the first draft of the manuscript. All authors wrote the final draft of the manuscript.

## Competing interests

A.K.A., Y.H.R., H.G. and D.P.B. are inventors on a patent filed by Arc Institute relevant to State. D.G. acknowledges outside interest as part of the founding team of the Autoscience Institute. D.P.B. acknowledges outside interest as a Google Advisor. H.G. acknowledges outside interest as a co-founder of Exai Bio, Tahoe Therapeutics, and Therna Therapeutics, serves on the board of directors at Exai Bio, and is a scientific advisory board member for Verge Genomics and Deep Forest Biosciences. L.A.G. is a cofounder of nChroma Bio and a Scientific Advisory Board member of Myllia Biotechnology. P.D.H. acknowledges outside interest as a co-founder of Monet AI, Terrain Biosciences, and Stylus Medicine, serves on the board of directors at Stylus Medicine, is a board observer at EvolutionaryScale and Terrain Biosciences, a scientific advisory board member at Arbor Biosciences and Veda Bio, and an advisor to NFDG, Varda Space, and Vial Health. Y.H.R. is a scientific advisory board member at QureXR.

## 8. Supplementary Figures

**Figure S1:**
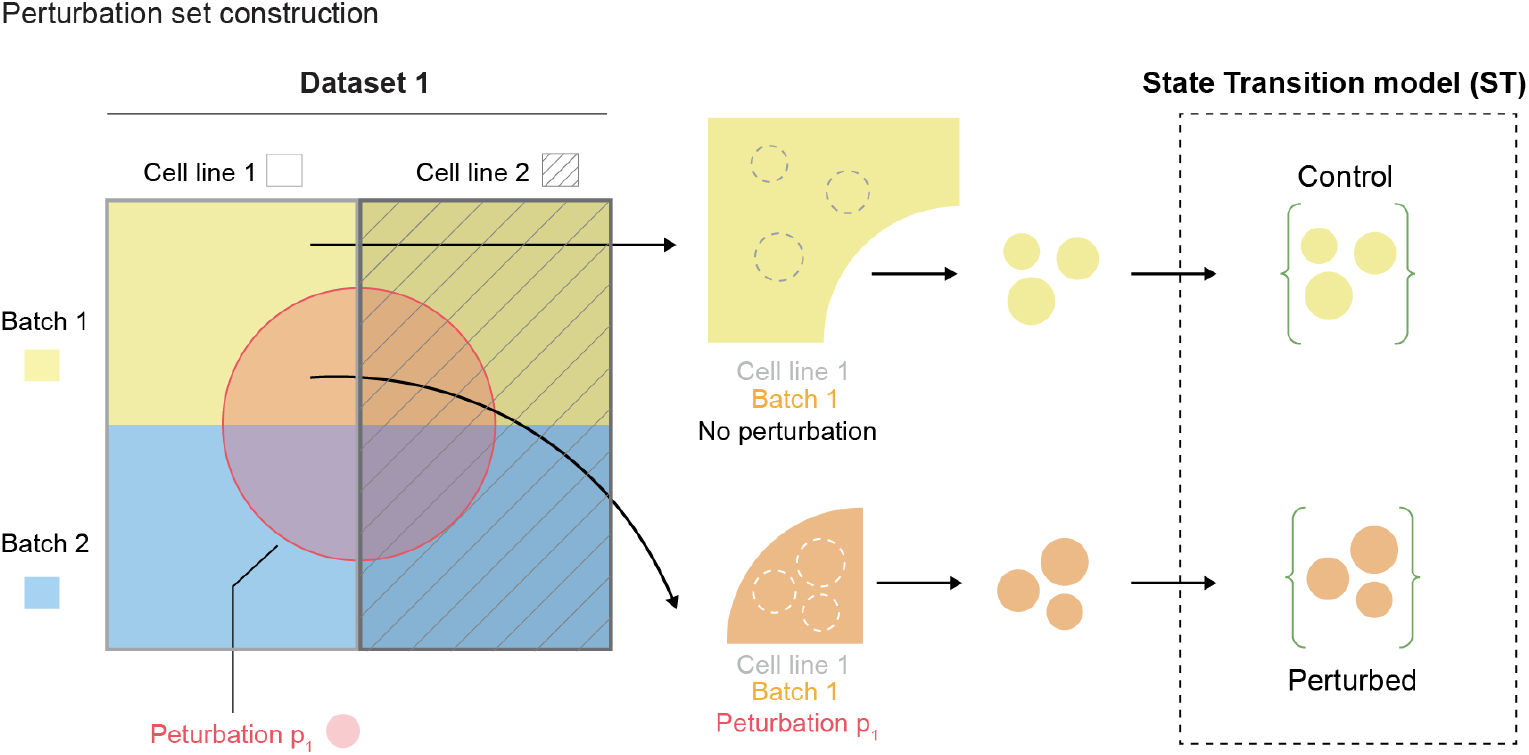
Cell set design for the ST perturbation model. Cells were grouped based on shared features (e.g., tissue, lineage, or batch), enabling the model to learn perturbation effects conditioned on set size and the matched covariates.

**Figure S2:**
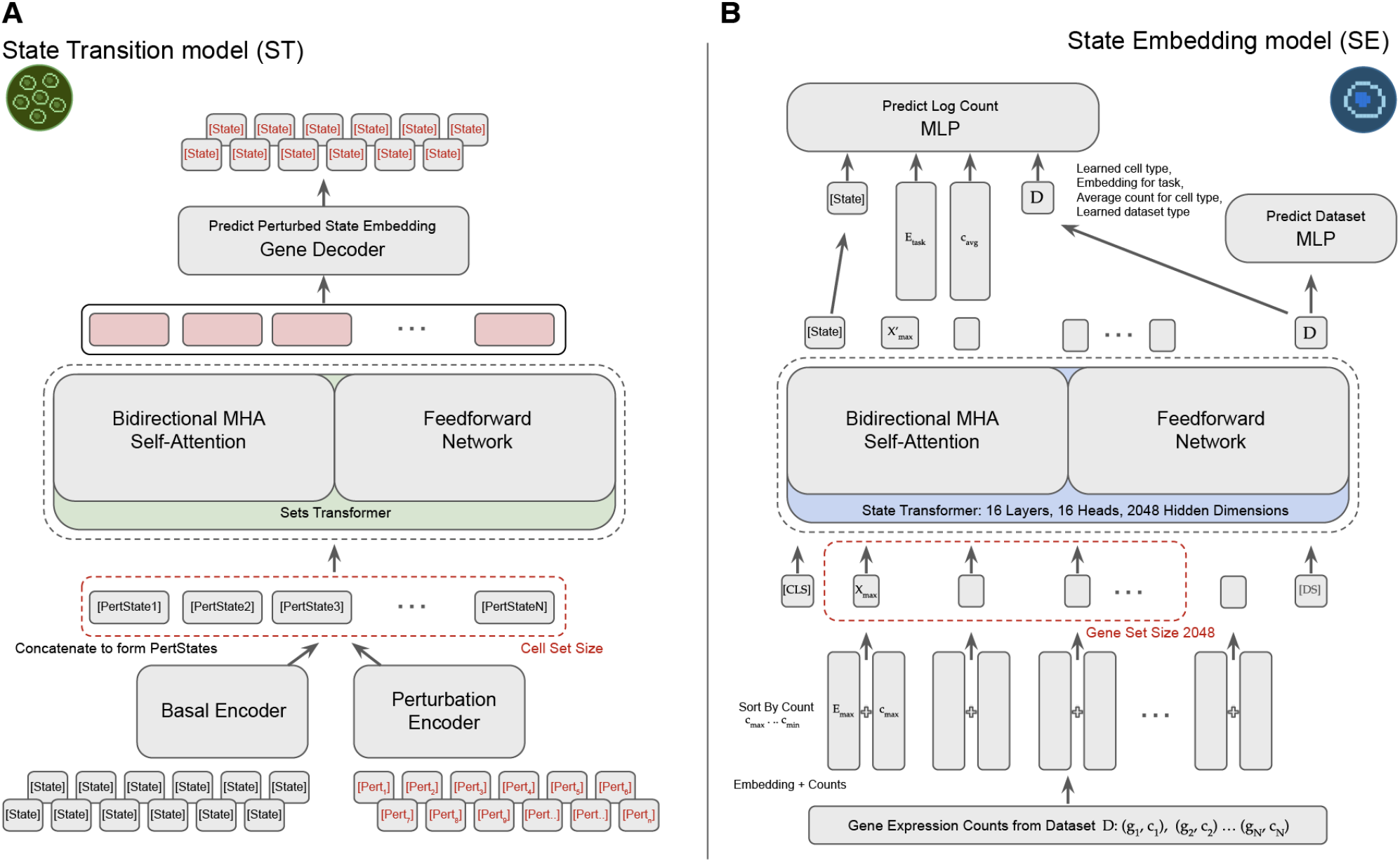
Comprehensive model architecture visualization for State. **(A)** Model architecture for ST with inputs in embedding space. When working directly in gene expression space, the gene decoder is swapped out with the gene reconstruction head, a simple linear layer that projects the transformer outputs back to gene expression space. **(B)** Model architecture for SE. The transformer outputs for the gene set inputted into the model are not used for downstream perturbation prediction. The [CLS] token is transformed into a strong representation of cell state, [State], to predict gene expression variability.

**Figure S3:**
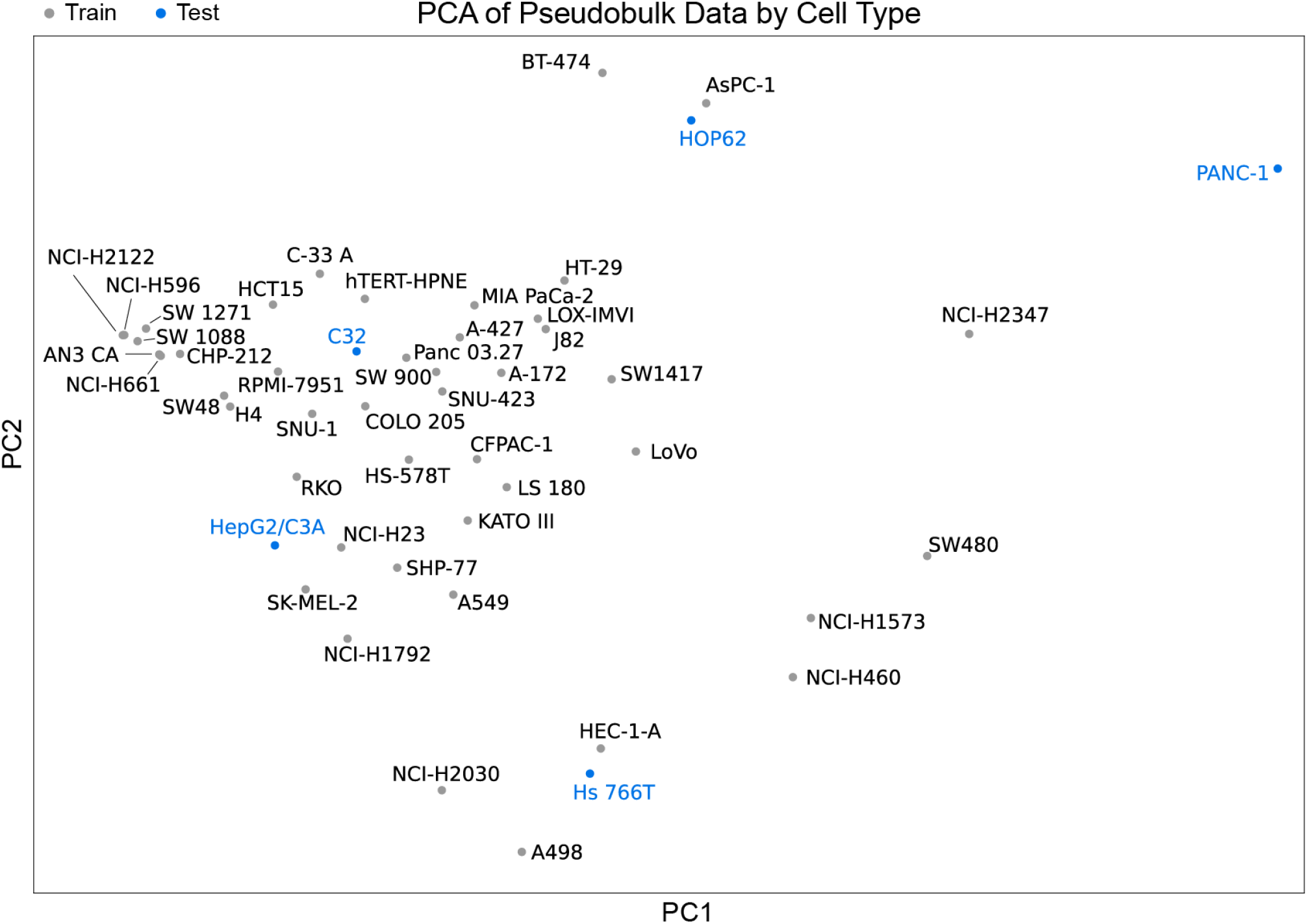
PCA visualization of Tahoe-100M pseudobulks reveals clusters of similar cell types. Pseudobulks were computed by averaging the counts over all cells with matching cell line, perturbation, and plate, and then aggregated into one pseudobulk per cell line. Generalization splits were chosen accordingly to not be too difficult (including some cell types near clusters), nor too easy (including some outlier cell types). The five holdout cell lines used in this analysis are: C32, HOP62, HepG2/C3A, Hs 766T, PANC-1.

**Figure S4:**
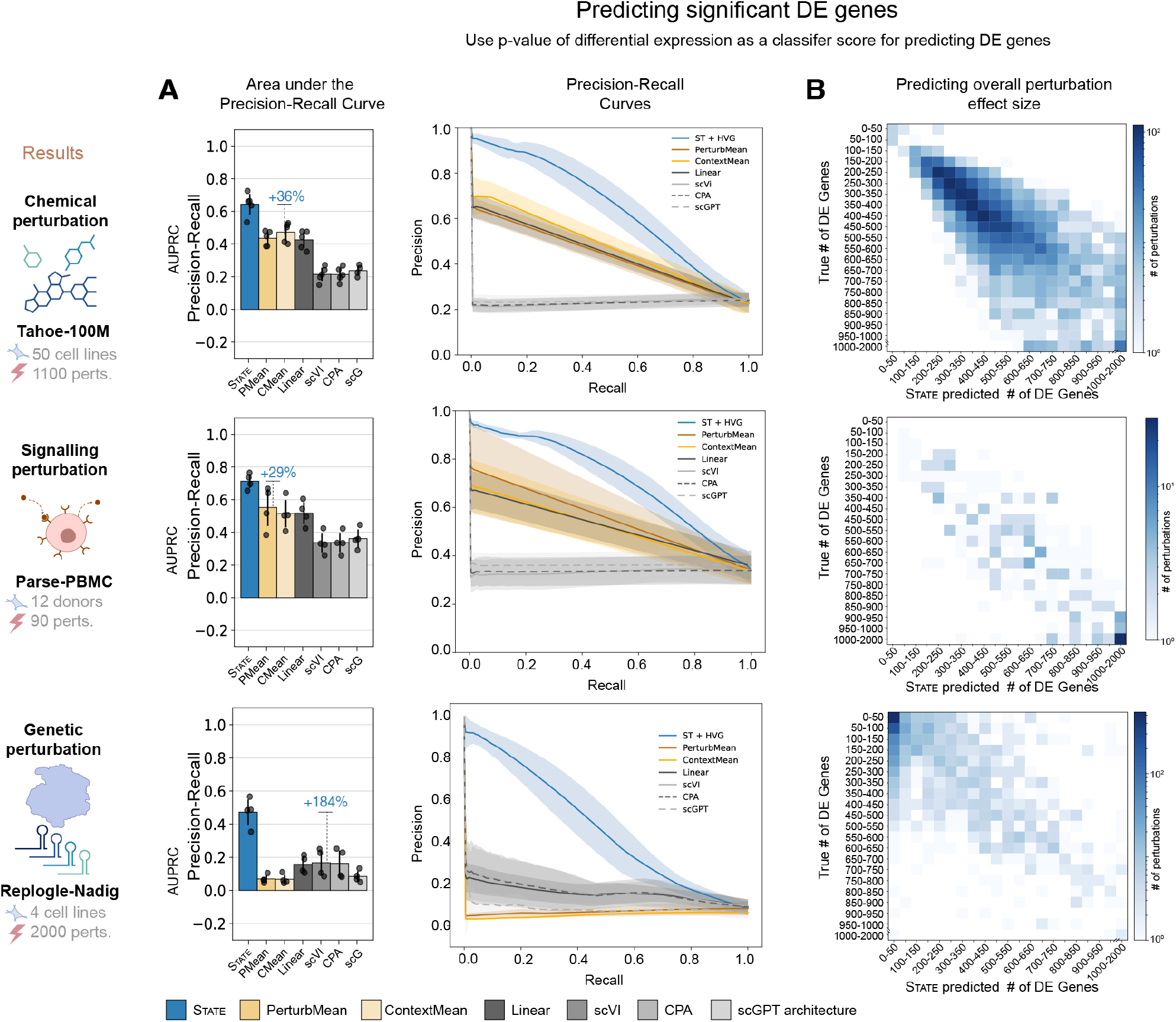
Additional metrics for evaluating identification of significant genes. Models were trained and evaluated on chemical, signaling and genetic perturbation datasets (Zhang et al., 2025; Replogle et al., 2022; Nadig et al., 2025). Comparisons included the mean baselines from (A), a simple linear model (Ahlmann-Eltze et al., 2024), autoencoder-based models (scVI (Lopez et al., 2018), CPA (Lotfollahi et al., 2023)), and a foundation model (scGPT (Cui et al., 2024)). Performance was assessed using standard Perturb-Seq outputs: expression counts and differentially expressed (DE) genes, with the following metrics considered: **(A)** Area Under the Precision-Recall Curve (AUPRC), summarizing model performance in identifying true DE genes, especially under class imbalance. **(B)** Heatmaps comparing the true overall perturbation effects as measured by number of differentially expressed genes with the predicted perturbation effect sizes by State.

**Figure S5:**
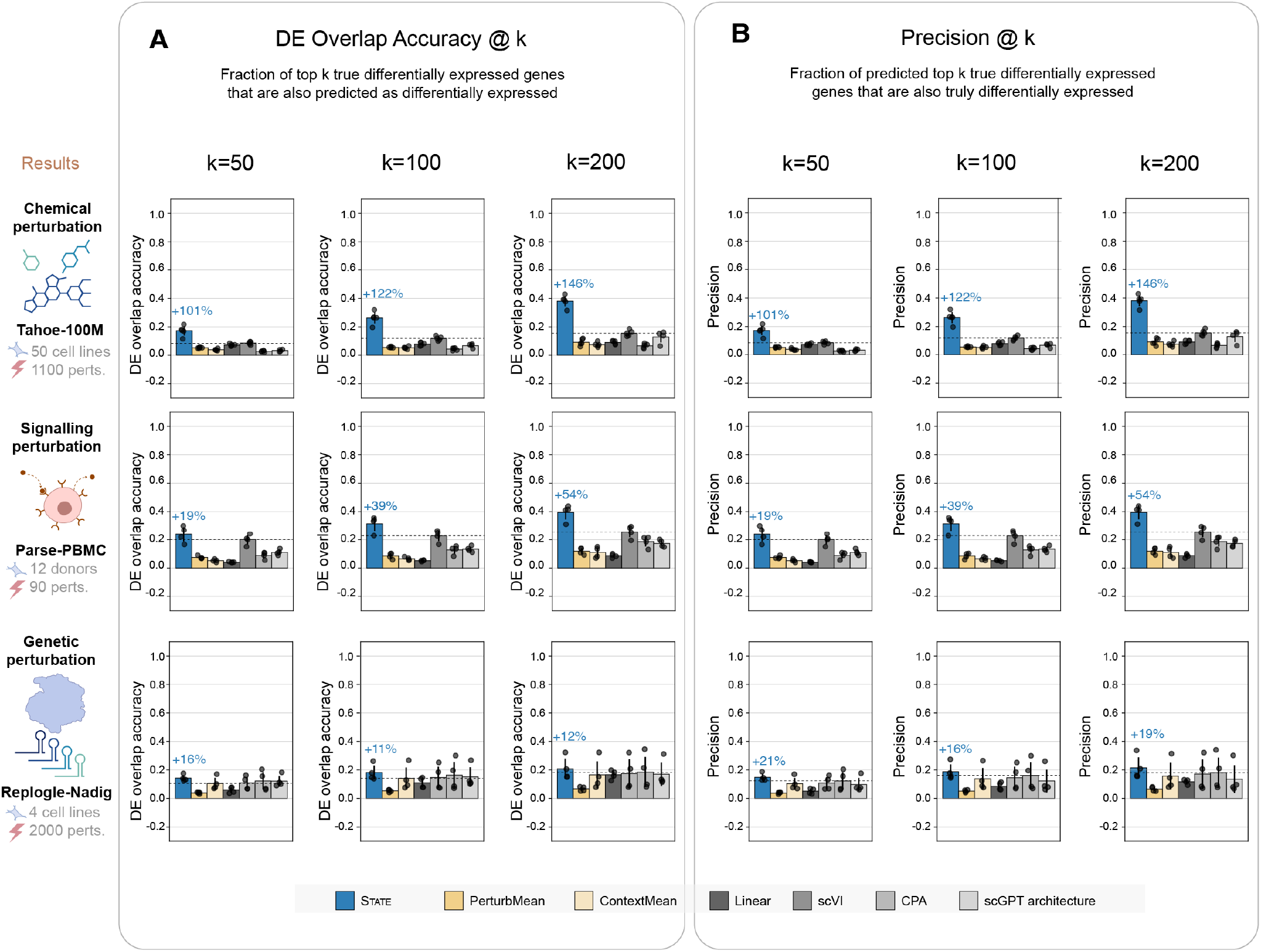
Additional metrics for evaluating top DE gene prediction. Models were trained and evaluated on both chemical, signaling and genetic perturbation datasets (Zhang et al., 2025; Replogle et al., 2022; Nadig et al., 2025). Comparisons included the mean baselines from (A), a simple linear model (Ahlmann-Eltze et al., 2024), autoencoder-based models (scVI (Lopez et al., 2018), CPA (Lotfollahi et al., 2023)), and a foundation model (scGPT (Cui et al., 2024)). Performance was assessed using standard Perturb-Seq outputs: expression counts and differentially expressed (DE) genes, with the following metrics considered: **(A)** Overlap@k: fraction of top-k differentially expressed genes in the observed data that were also identified in model predictions, evaluated across various values of k. **(B)** Precision@k: proportion of top-k predicted DE genes that were truly significant in the observed data, measuring model precision at varying *k* thresholds.

**Figure S6:**
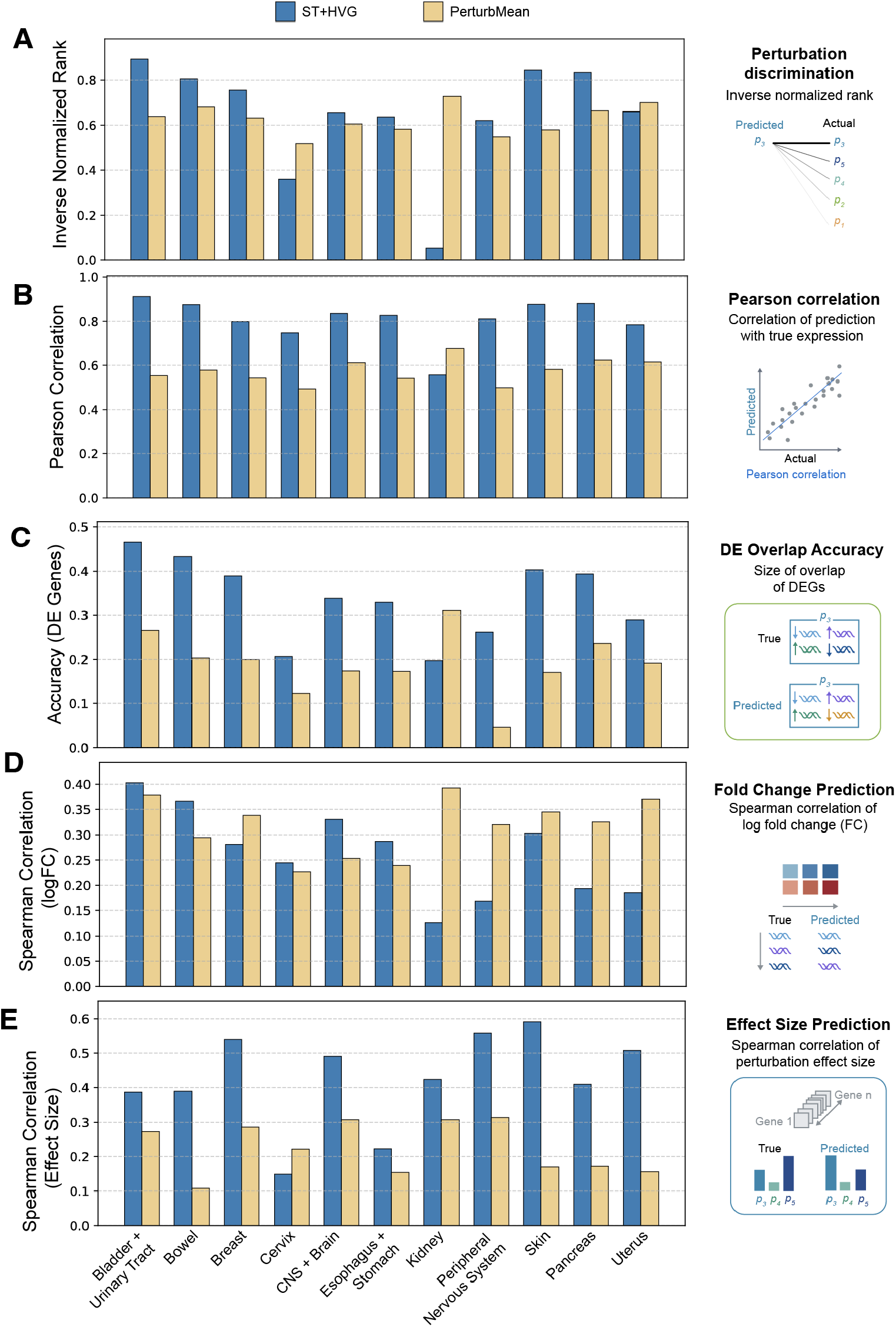
Tahoe performance by tissue. Performance comparison on five core metrics for the State model trained to generalize across tissues. For each split, all cell lines belonging to a specific tissue were included in the test set. 30% of perturbations from the test cell line were moved to the training set. **(A)** Perturbation discrimination score. **(B)** Pearson correlation of predicted and observed expression profiles. **(C)** DE gene overlap. **(D)** Log fold-change Spearman correlation for significant DE genes. **(E)** Effect size correlation.

**Figure S7:**
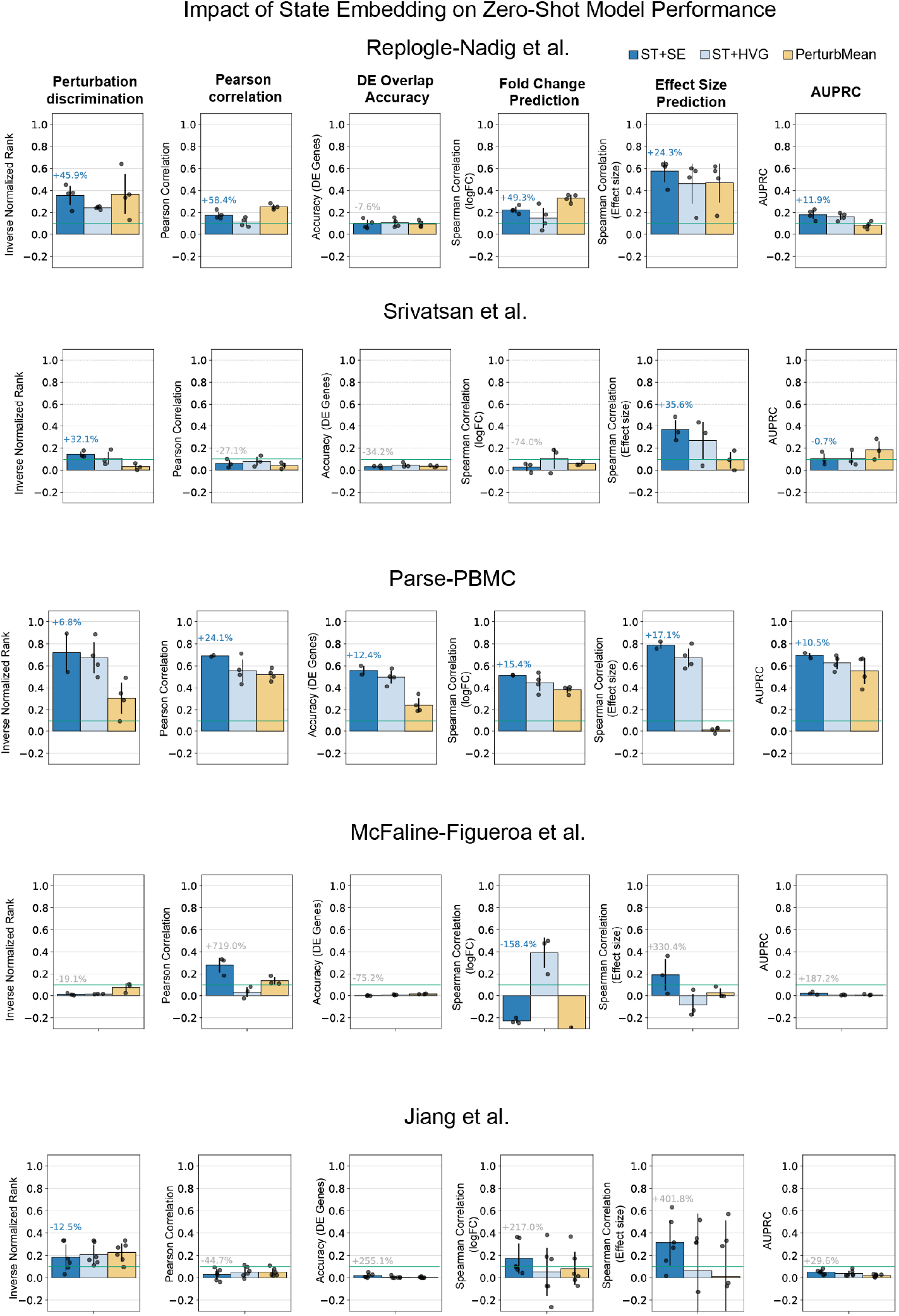
Impact of State embedding on zero-shot model performance. Model performance on perturbation effect prediction on a context for which perturbation data is completely unobserved during training. Performance improvements correspond to change between ST + SE and ST + HVG. Six metrics are considered (i) Perturbation Discrimination Score (ii) Pearson Correlation (iii) AUPRC (iv) Spearman correlation (log fold changes) (v) DE Gene Overlap Accuracy (vi) Spearman correlation (perturbation effect size).

**Figure S8:**
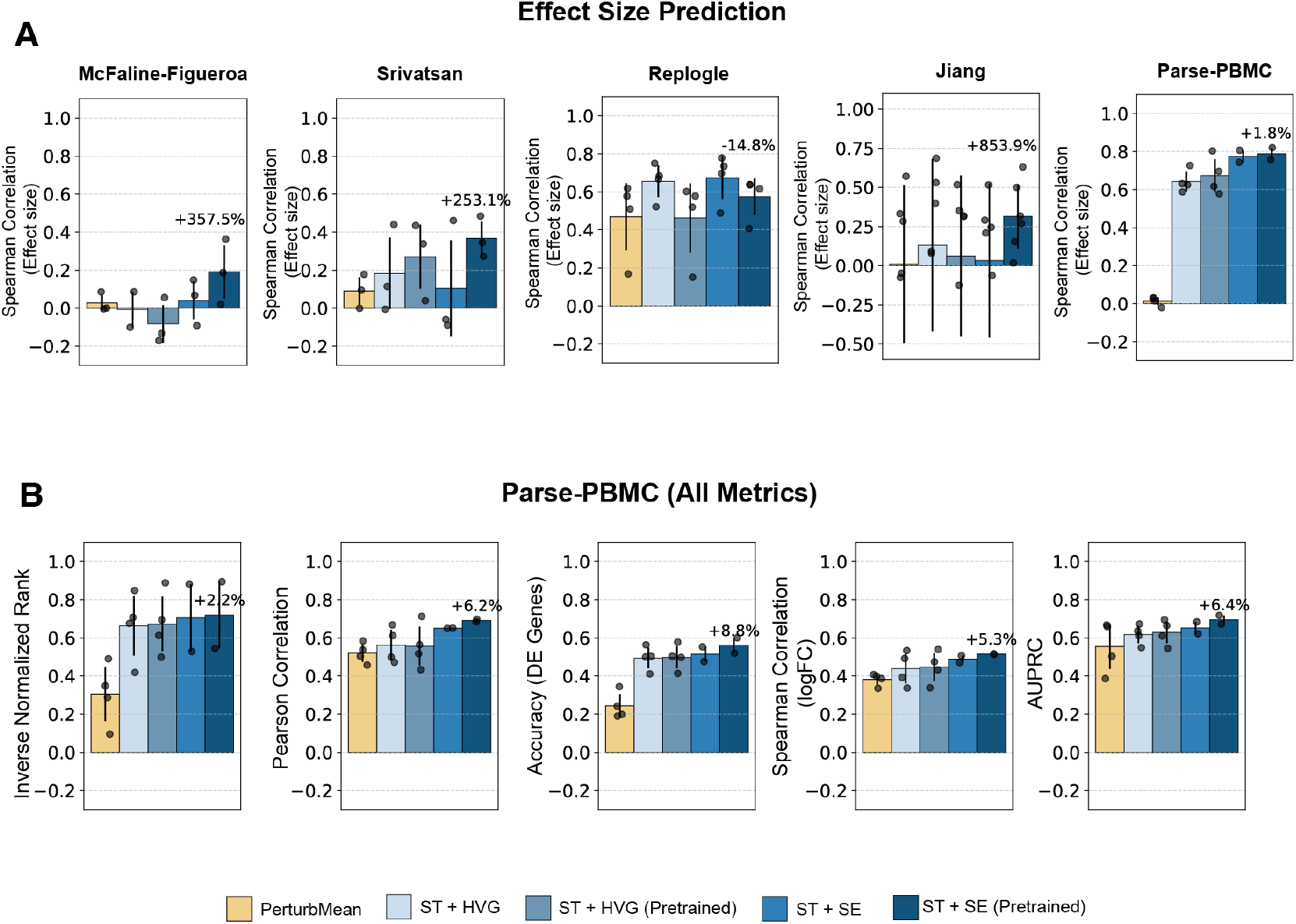
Impact of State embedding pretraining on zero-shot model performance. **(A)** Spearman correlation between model predicted and true perturbation effect sizes when predicting on a context for which perturbation data is completely unobserved during training. Performance improvements correspond to change between ST + SE and ST + SE (Pretrained). Five datasets are considered. **(B)** Model performance on perturbation effect prediction zero-shot on a context that is unobserved during training. Performance improvements correspond to change between ST + SE and ST + SE (Pretrained). Six metrics are considered (i) Perturbation Discrimination Score (ii) Pearson Correlation (iii) AUPRC (iv) Spearman correlation (log fold changes) (v) DE Gene Overlap Accuracy (vi) Spearman correlation (perturbation effect size).

**Figure S9:**
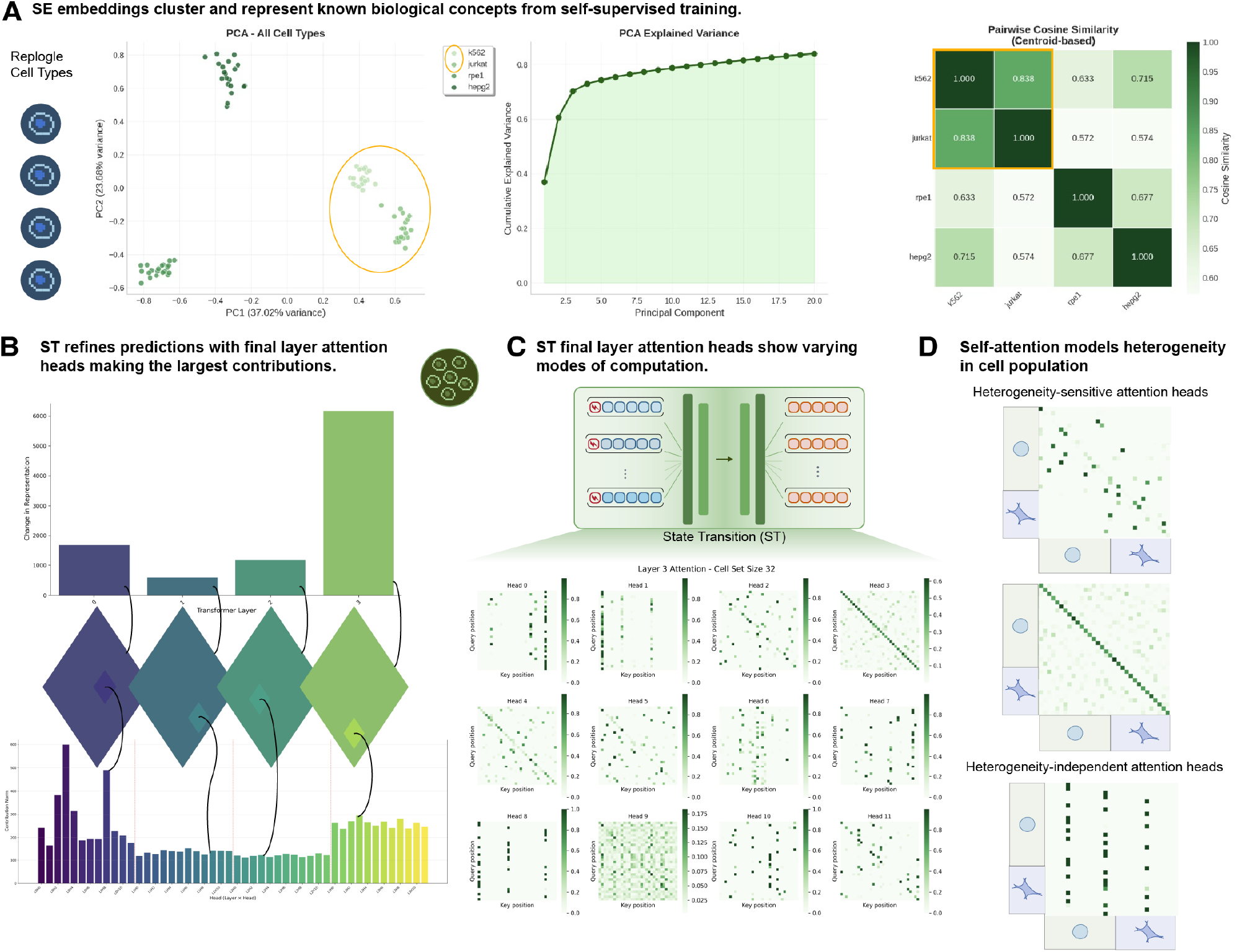
Initial interpretability analyses indicate how State leverages learnt biological information to make it’s predictions. **(A)** SE, without biological labels, learns that K-562 and Jurkat share similarities (both being from leukemia cell lines) just from gene expression data. **(B)** The final layer in ST contributes largely to the changing hidden state, across all attention heads. **(C)** Visualizing the attention heads (Query *×* Key) for the final layer displays striking variation across attention heads. **(D)** Constructing an inference-time example with a mixed cell set of K-562 and Jurkat visually displays how self-attention can model heterogeneity across the cells in the input set.

**Figure S10:**
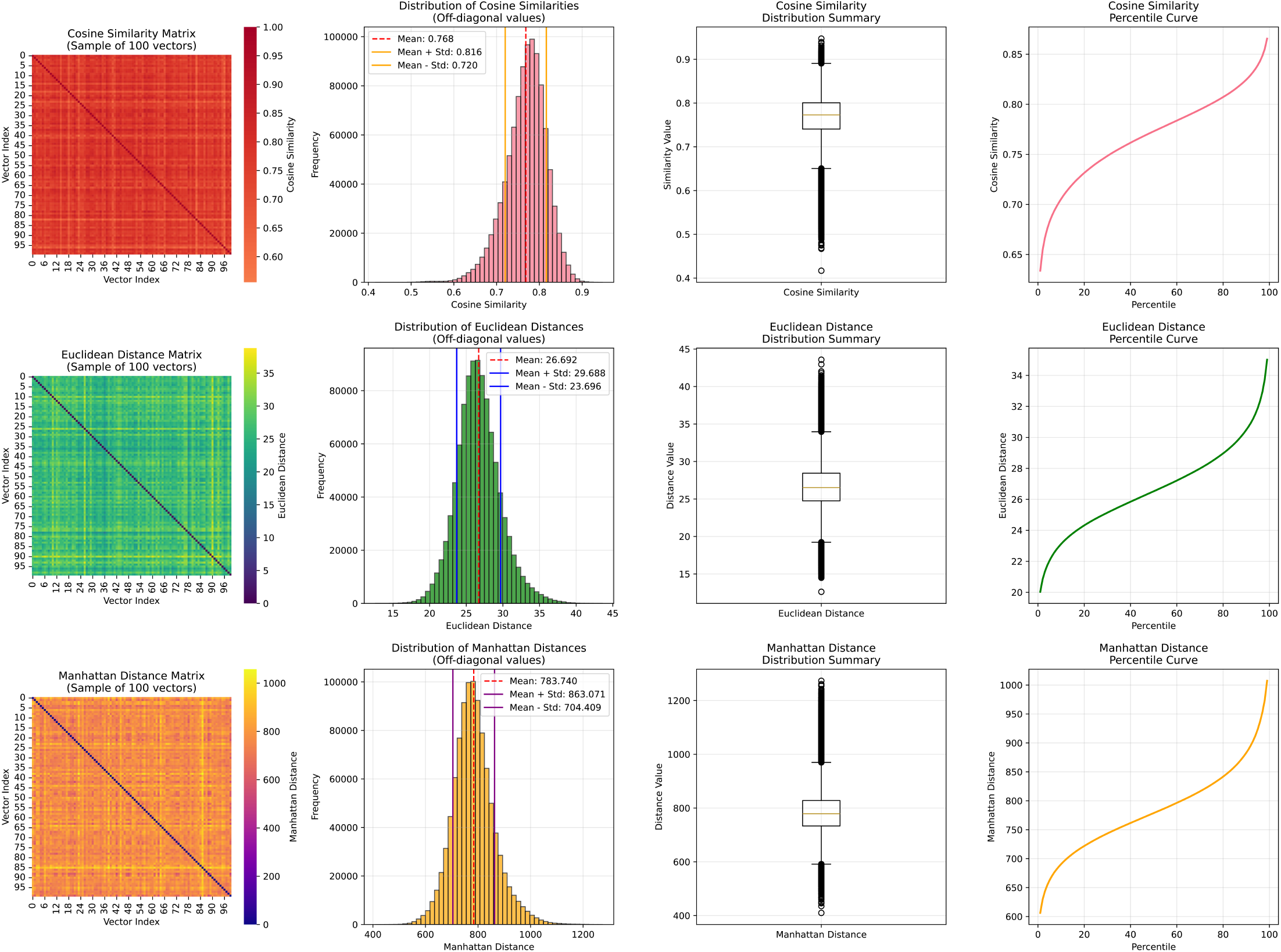
Distributions of real gene expression data with different discrimination operations displays how cosine similarity skews ranking metrics. We find that cosine similarity, by not accounting for the magnitudes of genes in expression data, reflects excessive similarity between real gene expression values and predicted values, which causes a distribution shift in downstream rank based metrics. Comparably, Euclidean distances (L2) and Manhattan distances (L1) better separate varying gene expressions within groups of cells.

## Notes

### Competing Interest Statement

A.K.A., Y.H.R., H.G. and D.P.B. are inventors on a patent filed by Arc Institute relevant to \textsc{State}. D.G. acknowledges outside interest as part of the founding team of the Autoscience Institute. D.P.B. acknowledges outside interest as a Google Advisor. H.G. acknowledges outside interest as a co-founder of Exai Bio, Tahoe Therapeutics, and Therna Therapeutics, serves on the board of directors at Exai Bio, and is a scientific advisory board member for Verge Genomics and Deep Forest Biosciences. L.A.G. is a co-founder of nChroma Bio and a Scientific Advisory Board member of Myllia Biotechnology. P.D.H. acknowledges outside interest as a co-founder of Monet AI, Terrain Biosciences, and Stylus Medicine, serves on the board of directors at Stylus Medicine, is a board observer at EvolutionaryScale and Terrain Biosciences, a scientific advisory board member at Arbor Biosciences and Veda Bio, and an advisor to NFDG, Varda Space, and Vial Health.

### Summary of Updates

Updated text, methods and description of model fine-tuning

## References

1. C. Ahlmann-Eltze, W. Huber, and S. Anders. Deep learning-based predictions of gene perturbation effects do not yet outperform simple linear methods. BioRxiv, pages 2024–09, 2024.

2. E. Ameisen, J. Lindsey, A. Pearce, W. Gurnee, N. L. Turner, B. Chen, C. Citro, D. Abrahams, S. Carter, B. Hosmer, J. Marcus, M. Sklar, A. Templeton, T. Bricken, C. McDougall, H. Cunningham, T. Henighan, A. Jermyn, A. Jones, A. Persic, Z. Qi, T. Ben Thompson, S. Zimmerman, K. Rivoire, T. Conerly, C. Olah, and J. Batson. Circuit tracing: Revealing computational graphs in language models. Transformer Circuits Thread, 2025.

3. K. Banach, J. Kowalska, Z. Rzepka, A. Beberok, J. Rok, and D. Wrześniok. The role of uva radiation in ketoprofen-mediated braf-mutant amelanotic melanoma cells death–a study at the cellular and molecular level. Toxicology in Vitro, 72:105108, 2021.

4. D. Bau, B. Zhou, A. Khosla, A. Oliva, and A. Torralba. Network dissection: Quantifying interpretability of deep visual representations. In Proceedings of the IEEE conference on computer vision and pattern recognition, 2017.

5. M. Bereket and T. Karaletsos. Modelling cellular perturbations with the sparse additive mechanism shift variational autoencoder. Advances in Neural Information Processing Systems,36, 2024.

6. A. Bhaskar, A. Wettig, D. Friedman, and D. Chen. Finding transformer circuits with edge pruning. Advances in Neural Information Processing Systems, 37, 2024.

7. S. Biswal, K. Elamvazhuthi, and R. Sonthalia. Identification of mean-field dynamics using transformers. arXiv preprint 2410.16295, 2024.

8. C. Bock, P. Datlinger, F. Chardon, M. A. Coelho, M. B. Dong, K. A. Lawson, T. Lu, L. Maroc, T. M. Norman, B. Song, et al. High-content crispr screening. Nature Reviews Methods Primers, 2(1):1–23, 2022.

9. P. Boyeau, J. Hong, A. Gayoso, M. Kim, J. L. McFaline-Figueroa, M. I. Jordan, E. Azizi, C. Ergen, and N. Yosef. Deep generative modeling of sample-level heterogeneity in singlecell genomics. BioRxiv, pages 2022–10, 2022.

10. Y. Brenier. Polar factorization and monotone rearrangement of vector-valued functions. Communications on pure and applied mathematics, 44(4):375–417, 1991.

11. T. Bricken, A. Templeton, J. Batson, B. Chen, A. Jermyn, T. Conerly, N. Turner, C. Anil, C. Denison, A. Askell, R. Lasenby, Y. Wu, S. Kravec, N. Schiefer, T. Maxwell, N. Joseph, Z. Hatfield-Dodds, A. Tamkin, K. Nguyen, B. McLean, J. E. Burke, T. Hume, S. Carter, T. Henighan, and C. Olah. Towards monosemanticity: Decomposing language models with dictionary learning. Transformer Circuits Thread, 2023.

12. G. Brixi, M. G. Durrant, J. Ku, M. Poli, G. Brockman, D. Chang, G. A. Gonzalez, S. H. King, D. B. Li, A. T. Merchant, et al. Genome modeling and design across all domains of life with evo 2. BioRxiv, pages 2025–02, 2025.

13. C. Bunne, S. G. Stark, G. Gut, J. S. Del Castillo, M. Levesque, K.-V. Lehmann, L. Pelkmans, A. Krause, and G. Rätsch. Learning single-cell perturbation responses using neural optimal transport. Nature methods, 20(11):1759–1768, 2023.

14. C. Bunne, Y. Roohani, Y. Rosen, A. Gupta, X. Zhang, M. Roed, T. Alexandrov, M. AlQuraishi, P. Brennan, D. B. Burkhardt, et al. How to build the virtual cell with artificial intelligence: Priorities and opportunities. Cell, 187(25):7045–7063, 2024a.

15. C. Bunne, G. Schiebinger, A. Krause, A. Regev, and M. Cuturi. Optimal transport for single-cell and spatial omics. Nature Reviews Methods Primers, 4(1):58, 2024b.

16. L. A. Caffarelli. The regularity of mappings with a convex potential. Journal of the American Mathematical Society, 5(1):99–104, 1992.

17. H. Chefer, S. Gur, and L. Wolf. Transformer interpretability beyond attention visualization. In Proceedings of the IEEE/CVF conference on computer vision and pattern recognition, pages 782–791, 2021.

18. Y. Chen and J. Zou. Genept: a simple but effective foundation model for genes and cells built from chatgpt. bioRxiv, pages 2023–10, 2024.

19. Y. Chen, Z. Hu, W. Chen, and H. Huang. Fast and scalable wasserstein-1 neural optimal transport solver for single-cell perturbation prediction. arXiv preprint 2411.00614, 2024.

20. M. Chevalley, Y. Roohani, A. Mehrjou, J. Leskovec, and P. Schwab. Causalbench: A largescale benchmark for network inference from single-cell perturbation data. arXiv preprint 2210.17283, 2022.

21. K. Clark, U. Khandelwal, O. Levy, and C. D. Manning. What does bert look at? an analysis of bert’s attention. arXiv preprint 1906.04341, 2019.

22. H. Cui, C. Wang, H. Maan, K. Pang, F. Luo, N. Duan, and B. Wang. scgpt: toward building a foundation model for single-cell multi-omics using generative ai. Nature Methods, 21(8): 1470–1480, 2024.

23. M. Cuturi. Sinkhorn distances: Lightspeed computation of optimal transport. Advances in neural information processing systems, 26, 2013.

24. H. Daneshmand. Provable optimal transport with transformers: The essence of depth and prompt engineering. arXiv preprint 2410.19931, 2024.

25. E. Dann, N. C. Henderson, S. A. Teichmann, M. D. Morgan, and J. C. Marioni. Differential abundance testing on single-cell data using k-nearest neighbor graphs. Nature biotechnology, 40(2):245–253, 2022.

26. T. Dao. Flashattention-2: Faster attention with better parallelism and work partitioning. In The Twelfth International Conference on Learning Representations, 2024.

27. P. Datlinger, A. F. Rendeiro, C. Schmidl, T. Krausgruber, P. Traxler, J. Klughammer, L. C. Schuster, A. Kuchler, D. Alpar, and C. Bock. Pooled crispr screening with single-cell transcriptome readout. Nature methods, 14(3):297–301, 2017.

28. V. De la Pena and E. Giné. Decoupling: from dependence to independence. Springer Science & Business Media, 2012.

29. A. Demir, E. Solovyeva, J. Boylan, M. Xiao, F. Serluca, S. Hoersch, J. Jenkins, M. Devarakonda, and B. Kiziltan. sc-otgm: Single-cell perturbation modeling by solving optimal mass transport on the manifold of gaussian mixtures. arXiv preprint 2405.03726, 2024.

30. J. Ding, J. Lin, S. Jiang, Y. Wang, Z. Miao, Z. Fang, J. Tang, M. Li, and X. Qiu. Toward a privacy-preserving predictive foundation model of single-cell transcriptomics with federated learning and tabular modeling. bioRxiv, pages 2025–01, 2025.

31. A. Dixit, O. Parnas, B. Li, J. Chen, C. P. Fulco, L. Jerby-Arnon, N. D. Marjanovic, D. Dionne, T. Burks, R. Raychowdhury, et al. Perturb-seq: dissecting molecular circuits with scalable single-cell rna profiling of pooled genetic screens. cell, 167(7):1853–1866, 2016.

32. M. Dong, B. Wang, J. Wei, A. H. de O. Fonseca, C. J. Perry, A. Frey, F. Ouerghi, E. F. Foxman, J. J. Ishizuka, R. M. Dhodapkar, et al. Causal identification of single-cell experimental perturbation effects with cinema-ot. Nature methods, 20(11):1769–1779, 2023.

33. J. Dunefsky, P. Chlenski, and N. Nanda. Transcoders find interpretable llm feature circuits. In The Thirty-eighth Annual Conference on Neural Information Processing Systems, 2024.

34. S. Elfwing, E. Uchibe, and K. Doya. Sigmoid-weighted linear units for neural network function approximation in reinforcement learning. Neural networks, 107:3–11, 2018.

35. C. Feng, E. M. Peets, Y. Zhou, L. Crepaldi, S. Usluer, A. Dunham, J. M. Braunger, J. Su, M. E. Strauss, D. Muraro, et al. A genome-scale single cell crispri map of trans gene regulation across human pluripotent stem cell lines. bioRxiv, pages 2024–11, 2024.

36. J. Feydy, T. Séjourné, F.-X. Vialard, S.-i. Amari, A. Trouve, and G. Peyré. Interpolating between optimal transport and mmd using sinkhorn divergences. In The 22nd International Conference on Artificial Intelligence and Statistics, pages 2681–2690, 2019.

37. F. Fischer, D. S. Fischer, R. Mukhin, A. Isaev, E. Biederstedt, A.-C. Villani, and F. J. Theis. sctab: scaling cross-tissue single-cell annotation models. Nature Communications, 15(1):6611, 2024.

38. T. Furuya, M. V. de Hoop, and G. Peyré. Transformers are universal in-context learners. arXiv preprint 2408.01367, 2024.

39. A. Gretton, K. M. Borgwardt, M. J. Rasch, B. Schölkopf, and A. Smola. A kernel two-sample test. Journal of Machine Learning Research, 13(25):723–773, 2012.

40. S. Gunasekar, J. Lee, D. Soudry, and N. Srebro. Characterizing implicit bias in terms of optimization geometry. In International Conference on Machine Learning, pages 1832– 1841. PMLR, 2018.

41. M. Hao, J. Gong, X. Zeng, C. Liu, Y. Guo, X. Cheng, T. Wang, J. Ma, X. Zhang, and L. Song. Large-scale foundation model on single-cell transcriptomics. Nature methods, 21 (8):1481–1491, 2024.

42. P. Hase, M. Bansal, B. Kim, and A. Ghandeharioun. Does localization inform editing? surprising differences in causality-based localization vs. knowledge editing in language models. Advances in Neural Information Processing Systems, 36, 2023.

43. T. Hayes, R. Rao, H. Akin, N. J. Sofroniew, D. Oktay, Z. Lin, R. Verkuil, V. Q. Tran, J. Deaton, M. Wiggert, et al. Simulating 500 million years of evolution with a language model. Science, page eads0018, 2025.

44. K. He, X. Zhang, S. Ren, and J. Sun. Delving deep into rectifiers: Surpassing human-level performance on imagenet classification. In Proceedings of the IEEE international conference on computer vision, pages 1026–1034, 2015.

45. G. Heimberg, R. Bhatnagar, H. El-Samad, and M. Thomson. Low dimensionality in gene expression data enables the accurate extraction of transcriptional programs from shallow sequencing. Cell systems, 2(4):239–250, 2016.

46. N. Ho, C. N. Ellington, J. Hou, S. Addagudi, S. Mo, T. Tao, D. Li, Y. Zhuang, H. Wang, X. Cheng, et al. Scaling dense representations for single cell with transcriptome-scale context. bioRxiv, pages 2024–11, 2024.

47. W. Hoeffding. The strong law of large numbers for u-statistics. Univ. of North Carolina Institute of statistics. Mimeo Series, 302, 1961.

48. A. C. Huang, T.-H. S. Hsieh, J. Zhu, J. Michuda, A. Teng, S. Kim, E. M. Rumsey, S. K. Lam, I. Anigbogu, P. Wright, et al. X-atlas/orion: Genome-wide perturb-seq datasets via a scalable fix-cryopreserve platform for training dose-dependent biological foundation models. bioRxiv, pages 2025–06, 2025a.

49. K. Huang, S. Zhang, H. Wang, Y. Qu, Y. Lu, Y. Roohani, R. Li, L. Qiu, J. Zhang, Y. Di, et al. Biomni: A general-purpose biomedical ai agent. bioRxiv, pages 2025–05, 2025b.

50. Y. Ji, M. Lotfollahi, F. A. Wolf, and F. J. Theis. Machine learning for perturbational single-cell omics. Cell Systems, 12(6):522–537, 2021.

51. J. Jiang, C. Wang, R. Qi, H. Fu, and Q. Ma. scread: a single-cell rna-seq database for alzheimer’s disease. Iscience, 23(11), 2020.

52. L. Jiang, C. Dalgarno, E. Papalexi, I. Mascio, H.-H. Wessels, H. Yun, N. Iremadze, G. Lithwick-Yanai, D. Lipson, and R. Satija. Systematic reconstruction of molecular pathway signatures using scalable single-cell perturbation screens. Nature Cell Biology, pages 1–13, 2025.

53. Q. Jiang, S. Chen, X. Chen, and R. Jiang. scpram accurately predicts single-cell gene expression perturbation response based on attention mechanism. Bioinformatics, 40(5): btae265, 2024.

54. D. Kalamkar, D. Mudigere, N. Mellempudi, D. Das, K. Banerjee, S. Avancha, D. T. Vooturi, N. Jammalamadaka, J. Huang, H. Yuen, J. Yang, J. Park, A. Heinecke, E. Georganas, S. Srinivasan, A. Kundu, M. Smelyanskiy, B. Kaul, and P. Dubey. A study of bfloat16 for deep learning training. arXiv preprint 1905.12322, 2019.

55. E. Kernfeld, Y. Yang, J. S. Weinstock, A. Battle, and P. Cahan. A systematic comparison of computational methods for expression forecasting. BioRxiv, pages 2023–07, 2023.

56. A. Kumar, L. Owen, N. R. Chowdhury, and F. Güra. Zclip: Adaptive spike mitigation for llm pre-training. arXiv preprint 2504.02507, 2025.

57. A. Lal, A. Karollus, L. Gunsalus, D. Garfield, S. Nair, A. M. Tseng, M. G. Gordon, J. Blischak, B. van de Geijn, T. Bhangale, et al. Decoding sequence determinants of gene expression in diverse cellular and disease states. bioRxiv, pages 2024–10, 2024.

58. D. Lara-Astiaso, A. Goñi-Salaverri, J. Mendieta-Esteban, N. Narayan, C. Del Valle, T. Gross, G. Giotopoulos, T. Beinortas, M. Navarro-Alonso, L. P. Aguado-Alvaro, et al. In vivo screening characterizes chromatin factor functions during normal and malignant hematopoiesis. Nature genetics, 55(9):1542–1554, 2023.

59. J. Lee, Y. Lee, J. Kim, A. Kosiorek, S. Choi, and Y. W. Teh. Set transformer: A framework for attention-based permutation-invariant neural networks. In International conference on machine learning, pages 3744–3753. PMLR, 2019.

60. C. Li, H. Gao, Y. She, H. Bian, Q. Chen, K. Liu, L. Wei, and X. Zhang. Benchmarking ai models for in silico gene perturbation of cells. bioRxiv, pages 2024–12, 2024a.

61. L. Li, Y. You, W. Liao, X. Fan, S. Lu, Y. Cao, B. Li, W. Ren, Y. Fu, J. Kong, et al. A systematic comparison of single-cell perturbation response prediction models. bioRxiv, pages 2024–12, 2024b.

62. Z. Lin, H. Akin, R. Rao, B. Hie, Z. Zhu, W. Lu, N. Smetanin, R. Verkuil, O. Kabeli, Y. Shmueli, et al. Evolutionary-scale prediction of atomic-level protein structure with a language model. Science, 379(6637):1123–1130, 2023.

63. R. Lopez, J. Regier, M. B. Cole, M. I. Jordan, and N. Yosef. Deep generative modeling for single-cell transcriptomics. Nature methods, 15(12):1053–1058, 2018.

64. R. Lopez, N. Tagasovska, S. Ra, K. Cho, J. Pritchard, and A. Regev. Learning causal representations of single cells via sparse mechanism shift modeling. In Conference on Causal Learning and Reasoning, pages 662–691. PMLR, 2023.

65. I. Loshchilov and F. Hutter. Decoupled weight decay regularization. In International Conference on Learning Representations, 2017.

66. M. Lotfollahi, F. A. Wolf, and F. J. Theis. scgen predicts single-cell perturbation responses. Nature methods, 16(8):715–721, 2019.

67. M. Lotfollahi, A. Klimovskaia Susmelj, C. De Donno, L. Hetzel, Y. Ji, I. L. Ibarra, S. R. Srivatsan, M. Naghipourfar, R. M. Daza, B. Martin, et al. Predicting cellular responses to complex perturbations in high-throughput screens. Molecular systems biology, 19(6): e11517, 2023.

68. M. D. Luecken, M. Büttner, K. Chaichoompu, A. Danese, M. Interlandi, M. F. Müller, D. C. Strobl, L. Zappia, M. Dugas, M. Colomé-Tatché, et al. Benchmarking atlas-level data integration in single-cell genomics. Nature methods, 19(1):41–50, 2022.

69. M. D. Luecken, S. Gigante, D. B. Burkhardt, R. Cannoodt, D. C. Strobl, N. S. Markov, L. Zappia, G. Palla, W. Lewis, D. Dimitrov, et al. Defining and benchmarking open problems in single-cell analysis. Nature Biotechnology, pages 1–6, 2025.

70. I. Lugowska, H. Koseła-Paterczyk, K. Kozak, and P. Rutkowski. Trametinib: a mek inhibitor for management of metastatic melanoma. OncoTargets and therapy, pages 2251–2259, 2015.

71. C. Lyle, Z. Zheng, K. Khetarpal, J. Martens, H. van Hasselt, R. Pascanu, and W. Dabney. Normalization and effective learning rates in reinforcement learning, 2024. URL https://arxiv.org/abs/2407.01800.

72. A. Makkuva, A. Taghvaei, S. Oh, and J. Lee. Optimal transport mapping via input convex neural networks. In International Conference on Machine Learning, pages 6672–6681. PMLR, 2020.

73. J. L. McFaline-Figueroa, A. J. Hill, X. Qiu, D. Jackson, J. Shendure, and C. Trapnell. A pooled single-cell genetic screen identifies regulatory checkpoints in the continuum of the epithelial-to-mesenchymal transition. Nature genetics, 51(9):1389–1398, 2019.

74. J. L. McFaline-Figueroa, S. Srivatsan, A. J. Hill, M. Gasperini, D. L. Jackson, L. Saunders, S. Domcke, S. G. Regalado, P. Lazarchuck, S. Alvarez, et al. Multiplex single-cell chemical genomics reveals the kinase dependence of the response to targeted therapy. Cell Genomics, 4(2), 2024.

75. A. Nadig, J. M. Replogle, A. N. Pogson, M. Murthy, S. A. McCarroll, J. S. Weissman, E. B. Robinson, and L. J. O’Connor. Transcriptome-wide analysis of differential expression in perturbation atlases. Nature Genetics, pages 1–10, 2025.

76. E. Nguyen, M. Poli, M. G. Durrant, B. Kang, D. Katrekar, D. B. Li, L. J. Bartie, A. W. Thomas, S. H. King, G. Brixi, et al. Sequence modeling and design from molecular to genome scale with evo. Science, 386(6723):eado9336, 2024.

77. T. M. Norman, M. A. Horlbeck, J. M. Replogle, A. Y. Ge, A. Xu, M. Jost, L. A. Gilbert, and J. S. Weissman. Exploring genetic interaction manifolds constructed from rich single-cell phenotypes. Science, 365(6455):786–793, 2019.

78. E. Noutahi, J. Hartford, P. Tossou, S. Whitfield, A. K. Denton, C. Wognum, K. Ulicna, J. Hsu, M. Cuccarese, E. Bengio, et al. Virtual cells: Predict, explain, discover. arXiv preprint 2505.14613, 2025.

79. C. Olah, A. Mordvintsev, and L. Schubert. Feature visualization. Distill, 2(11):e7, 2017.

80. E. Papalexi, E. P. Mimitou, A. W. Butler, S. Foster, B. Bracken, W. M. Mauck III, H.-H. Wessels, Y. Hao, B. Z. Yeung, P. Smibert, et al. Characterizing the molecular regulation of inhibitory immune checkpoints with multimodal single-cell screens. Nature genetics, 53 (3):322–331, 2021.

81. Parse Biosciences. 10 million human pbmcs in a single experiment, 2023. URL https://www.parsebiosciences.com/datasets/10-million-human-pbmcs-in-a-single-experiment/.

82. J. D. Pearce, S. E. Simmonds, G. Mahmoudabadi, L. Krishnan, G. Palla, A.-M. Istrate, A. Tarashansky, B. Nelson, O. Valenzuela, D. Li, et al. A cross-species generative cell atlas across 1.5 billion years of evolution: The transcriptformer single-cell model. bioRxiv, pages 2025–04, 2025.

83. S. Peidli, T. D. Green, C. Shen, T. Gross, J. Min, S. Garda, B. Yuan, L. J. Schumacher, J. P. Taylor-King, D. S. Marks, et al. scperturb: harmonized single-cell perturbation data. Nature Methods, 21(3):531–540, 2024.

84. S. Persad, Z.-N. Choo, C. Dien, N. Sohail, I. Masilionis, R. Chaligné, T. Nawy, C. C. Brown, R. Sharma, I. Pe’er, et al. Seacells infers transcriptional and epigenomic cellular states from single-cell genomics data. Nature Biotechnology, 41(12):1746–1757, 2023.

85. G. Peyré, M. Cuturi, et al. Computational optimal transport: With applications to data science. Foundations and Trends® in Machine Learning, 11(5-6):355–607, 2019.

86. Z. Piran, N. Cohen, Y. Hoshen, and M. Nitzan. Disentanglement of single-cell data with biolord. Nature Biotechnology, pages 1–6, 2024.

87. C. C. S. Program, S. Abdulla, B. Aevermann, P. Assis, S. Badajoz, S. M. Bell, E. Bezzi, B. Cakir, J. Chaffer, S. Chambers, et al. Cz cellxgene discover: a single-cell data platform for scalable exploration, analysis and modeling of aggregated data. Nucleic Acids Research, 53(D1):D886–D900, 2025.

88. L. Przybyla and L. A. Gilbert. A new era in functional genomics screens. Nature Reviews Genetics, 23(2):89–103, 2022.

89. A. Radford, J. Wu, R. Child, D. Luan, D. Amodei, I. Sutskever, et al. Language models are unsupervised multitask learners. OpenAI blog, 1(8):9, 2019.

90. J. M. Replogle, R. A. Saunders, A. N. Pogson, J. A. Hussmann, A. Lenail, A. Guna, L. Mascibroda, E. J. Wagner, K. Adelman, G. Lithwick-Yanai, et al. Mapping information-rich genotype-phenotype landscapes with genome-scale perturb-seq. Cell, 185(14):2559–2575, 2022.

91. S. A. Rizvi, D. Levine, A. Patel, S. Zhang, E. Wang, S. He, D. Zhang, C. Tang, Z. Lyu, R. Darji, et al. Scaling large language models for next-generation single-cell analysis. bioRxiv, pages 2025–04, 2025.

92. M. L. Rizzo and G.J. Székely. Energy distance. wiley interdisciplinary reviews: Computational statistics, 8(1):27–38, 2016.

93. J. E. Rood, A. Hupalowska, and A. Regev. Toward a foundation model of causal cell and tissue biology with a perturbation cell and tissue atlas. Cell, 187(17):4520–4545, 2024.

94. J. E. Rood, S. Wynne, L. Robson, A. Hupalowska, J. Randell, S. A. Teichmann, and A. Regev. The human cell atlas from a cell census to a unified foundation model. Nature, 637(8048):1065–1071, 2025.

95. Y. Roohani, K. Huang, and J. Leskovec. Predicting transcriptional outcomes of novel multigene perturbations with gears. Nature Biotechnology, 42(6):927–935, 2024a.

96. Y. Roohani, A. Lee, Q. Huang, J. Vora, Z. Steinhart, K. Huang, A. Marson, P. Liang, and J. Leskovec. Biodiscoveryagent: An ai agent for designing genetic perturbation experiments. arXiv preprint 2405.17631, 2024b.

97. Y. Rosen, Y. Roohani, A. Agarwal, L. Samotorčan, T. S. Consortium, S. R. Quake, and J. Leskovec. Universal cell embeddings: A foundation model for cell biology. bioRxiv, pages 2023–11, 2023.

98. J. Ryu, C. Bunne, L. Pinello, A. Regev, and R. Lopez. Cross-modality matching and prediction of perturbation responses with labeled gromov-wasserstein optimal transport. arXiv preprint 2405.00838, 2024.

99. R. A. Saunders, W. E. Allen, X. Pan, J. Sandhu, J. Lu, T. K. Lau, K. Smolyar, Z. A. Sullivan, C. Dulac, J. S. Weissman, et al. A platform for multimodal in vivo pooled genetic screens reveals regulators of liver function. bioRxiv, pages 2024–11, 2024.

100. B. Song, D. Liu, W. Dai, N. F. McMyn, Q. Wang, D. Yang, A. Krejci, A. Vasilyev, N. Untermoser, A. Loregger, et al. Decoding heterogeneous single-cell perturbation responses. Nature cell biology, pages 1–12, 2025.

101. D. Soudry, E. Hoffer, M. S. Nacson, S. Gunasekar, and N. Srebro. The implicit bias of gradient descent on separable data. Journal of Machine Learning Research, 19(70):1–57, 2018.

102. S. R. Srivatsan, J. L. McFaline-Figueroa, V. Ramani, L. Saunders, J. Cao, J. Packer, H. A. Pliner, D. L. Jackson, R. M. Daza, L. Christiansen, et al. Massively multiplex chemical transcriptomics at single-cell resolution. Science, 367(6473):45–51, 2020.

103. H. Steck, C. Ekanadham, and N. Kallus. Is cosine-similarity of embeddings really about similarity? In Companion Proceedings of the ACM Web Conference 2024, pages 887–890, 2024.

104. A. Szałata, K. Hrovatin, S. Becker, A. Tejada-Lapuerta, H. Cui, B. Wang, and F. J. Theis. Transformers in single-cell omics: a review and new perspectives. Nature methods, 21(8):1430–1443, 2024.

105. C. V. Theodoris, L. Xiao, A. Chopra, M. D. Chaffin, Z. R. Al Sayed, M. C. Hill, H. Mantineo, E. M. Brydon, Z. Zeng, X. S. Liu, et al. Transfer learning enables predictions in network biology. Nature, 618(7965):616–624, 2023.

106. H. Touvron, T. Lavril, G. Izacard, X. Martinet, M.-A. Lachaux, T. Lacroix, B. Rozière, N. Goyal, E. Hambro, F. Azhar, et al. Llama: Open and efficient foundation language models. arXiv preprint 2302.13971, 2023.

107. G. Vardi. On the implicit bias in deep-learning algorithms. Communications of the ACM, 66(6):86–93, 2023.

108. E. Weinberger, C. Lin, and S.-I. Lee. Isolating salient variations of interest in single-cell data with contrastivevi. Nature Methods, 20(9):1336–1345, 2023.

109. E. Weinberger, R. Conrad, and T. Ashuach. Modeling variable guide efficiency in pooled crispr screens with contrastivevi+. arXiv preprint 2411.08072, 2024.

110. A. Wenteler, M. Occhetta, N. Branson, M. Huebner, V. Curean, W. Dee, W. Connell, A. Hawkins-Hooker, P. Chung, Y. Ektefaie, et al. Perteval-scfm: Benchmarking single-cell foundation models for perturbation effect prediction. bioRxiv, pages 2024–10, 2024.

111. C. Wu, I. MacLeod, and A. I. Su. Biogps and mygene. info: organizing online, gene-centric information. Nucleic acids research, 41(D1):D561–D565, 2013.

112. M. Wu, R. Littman, J. Levine, L. Qiu, T. Biancalani, D. Richmond, and J.-C. Huetter. Contextualizing biological perturbation experiments through language. arXiv preprint 2502.21290, 2025.

113. Y. Wu, E. Wershof, S. M. Schmon, M. Nassar, B. Osiński, R. Eksi, K. Zhang, and T. Graepel. Perturbench: Benchmarking machine learning models for cellular perturbation analysis. arXiv preprint 2408.10609, 2024.

114. A. Yates, K. Beal, S. Keenan, W. McLaren, M. Pignatelli, G. R. Ritchie, M. Ruffier, K. Taylor, A. Vullo, and P. Flicek. The ensembl rest api: Ensembl data for any language. Bioinformatics, 31(1):143–145, 2015.

115. N. D. Youngblut, C. Carpenter, J. Prashar, C. Ricci-Tam, R. Ilango, N. Teyssier, S. Konermann, P. Hsu, A. Dobin, D. P. Burke, et al. scBaseCount: An AI agent-curated, uniformly processed, and continually expanding single cell data repository. bioRxiv, pages 2025–02, 2025.

116. J. X. Yu, J. M. Suh, K. D. Popova, K. Garcia, T. Joshi, B. Culbertson, J. B. Spinelli, V. Subramanyam, K. Lou, K. M. Shokat, J. Weissman, and H. Goodarzi. Multiplexed mosaic tumor models reveal natural phenotypic variations in drug response within and between populations. bioRxiv, 2024. doi: 10.1101/2024.12.13.628239.

117. M. Zaheer, S. Kottur, S. Ravanbakhsh, B. Poczos, R. R. Salakhutdinov, and A. J. Smola. Deep sets. Advances in neural information processing systems, 30, 2017.

118. M. D. Zeiler and R. Fergus. Visualizing and understanding convolutional networks. In Computer Vision–ECCV 2014: 13th European Conference, Zurich, Switzerland, September 6-12, 2014, Proceedings, Part I 13, pages 818–833. Springer, 2014.

119. J. Zhang, K. Greenewald, C. Squires, A. Srivastava, K. Shanmugam, and C. Uhler. Identifiability guarantees for causal disentanglement from soft interventions. Advances in Neural Information Processing Systems, 36:50254–50292, 2023.

120. J. Zhang, A. A. Ubas, R. de Borja, V. Svensson, N. Thomas, N. Thakar, I. Lai, A. Winters, U. Khan, M. G. Jones, J. D. Thompson, V. Tran, J. Pangallo, E. Papalexi, A. Sapre, H. Nguyen, O. Sanderson, M. Nigos, O. Kaplan, S. Schroeder, B. Hariadi, S. Marrujo, C. C. A. Salvino, G. Gallareta Olivares, R. Koehler, G. Geiss, A. Rosenberg, C. Roco, D. Merico, N. Alidoust, H. Goodarzi, and J. Yu. Tahoe-100m: A giga-scale single-cell perturbation atlas for context-dependent gene function and cellular modeling. bioRxiv, pages 2025–02, 2025.

